# Genetically encoded non-canonical amino acids reveal asynchronous dark reversion of chromophore, backbone and side-chains in EL222

**DOI:** 10.1101/2022.09.16.506679

**Authors:** Aditya S. Chaudhari, Aditi Chatterjee, Catarina A.O. Domingos, Prokopis C. Andrikopoulos, Yingliang Liu, Inger Andersson, Bohdan Schneider, Víctor A. Lórenz-Fonfría, Gustavo Fuertes

## Abstract

Photoreceptors containing the light-oxygen-voltage (LOV) domain elicit biological responses upon excitation of their flavin mononucleotide (FMN) chromophore by blue light. The mechanism and kinetics of dark-state recovery are not well understood. Here we incorporated the non-canonical amino acid p-cyanophenylalanine (CNF) by genetic code expansion technology at forty-five positions of the bacterial transcription factor EL222. Screening of light-induced changes in infrared (IR) absorption frequency, electric field and hydration of the nitrile groups identified residues CNF31 and CNF35 as reporters of monomer/oligomer and caged/decaged equilibria, respectively. Time-resolved multi-probe UV/Visible and IR spectroscopy experiments of the lit-to-dark transition revealed four dynamical events. Predominantly, rearrangements around the A’α helix interface (CNF31 and CNF35) precede FMN-cysteinyl adduct scission, folding of α-helices (amide bands), and relaxation of residue CNF151. This study illustrates the importance of characterizing all parts of a protein and suggests a key role for the N-terminal A’α extension of the LOV domain in controlling EL222 photocycle length.

**Significance:** The kinetics of fold switching between non-illuminated and blue-light-irradiated states in the transcription factor EL222 is important for understanding the signal transduction mechanism of LOV photoreceptors. Here we combine two native probes, the FMN chromophore (absorption bands in the UV/Visible region) and the protein backbone (amide bands in the infrared region), with genetically encoded cyano (C≡N)-containing phenylalanine residues as infrared reporters of protein microenvironments. EL222 structural dynamics is more complex than expected if using a single type of probe. Local changes around residues 31 and 35 precede FMN-protein adduct rupture, which in turn precedes the global protein conformational relaxation. Our findings point the way forward to obtaining comprehensive descriptions of kinetic transitions in LOV and other photosensors.

## Introduction

The conformation and dynamics of photoreceptor proteins are orchestrated by light, which regulates changes at the secondary, tertiary and sometimes quaternary levels of structure (*1*). In the photocontrolled transcription factor EL222 from *Erythrobacter litoralis*, the well-studied dark state is characterized by a monomeric “closed” conformation featuring extensive interactions between the light-oxygen-voltage (LOV) and helix-turn-helix (HTH) domains (*2*). Although high-resolution structural information of the lit state(s) of EL222 is missing, it is known that blue-light excitation of the flavin mononucleotide (FMN) chromophore embedded in the LOV domain leads to uncaging of the associated HTH module, oligomerization and subsequent DNA binding and gene expression (*2-6*). UV/Vis spectroscopy has been instrumental in monitoring the photoconversion of FMN (**Fig. 1** and **Fig. S1**). Non-irradiated LOV proteins absorb maximally at ∼450 nm while their irradiated counterparts peak at ∼390 nm (**Fig. 1B** left panel). This frequency shift is caused by photon absorption by the oxidized FMN cofactor, which triggers its partial reduction to a semiquinone radical and adduct formation with a conserved cysteine residue (*7*). FMN-Cys covalent bond formation takes place in the nanosecond to microsecond time scale (*5, 8*). The rupture of such an adduct state, sometimes referred to as A_390_ species (*5*), is much slower, typically returning to the resting state in seconds to hours, depending on the particular LOV domain (*9*). The dark recovery kinetics of isolated LOV domain suggests synchronous relaxation of chromophore and protein chain (*10*), but the behavior of multi-domain LOV-containing proteins is largely unknown.

**FIGURE 1.**
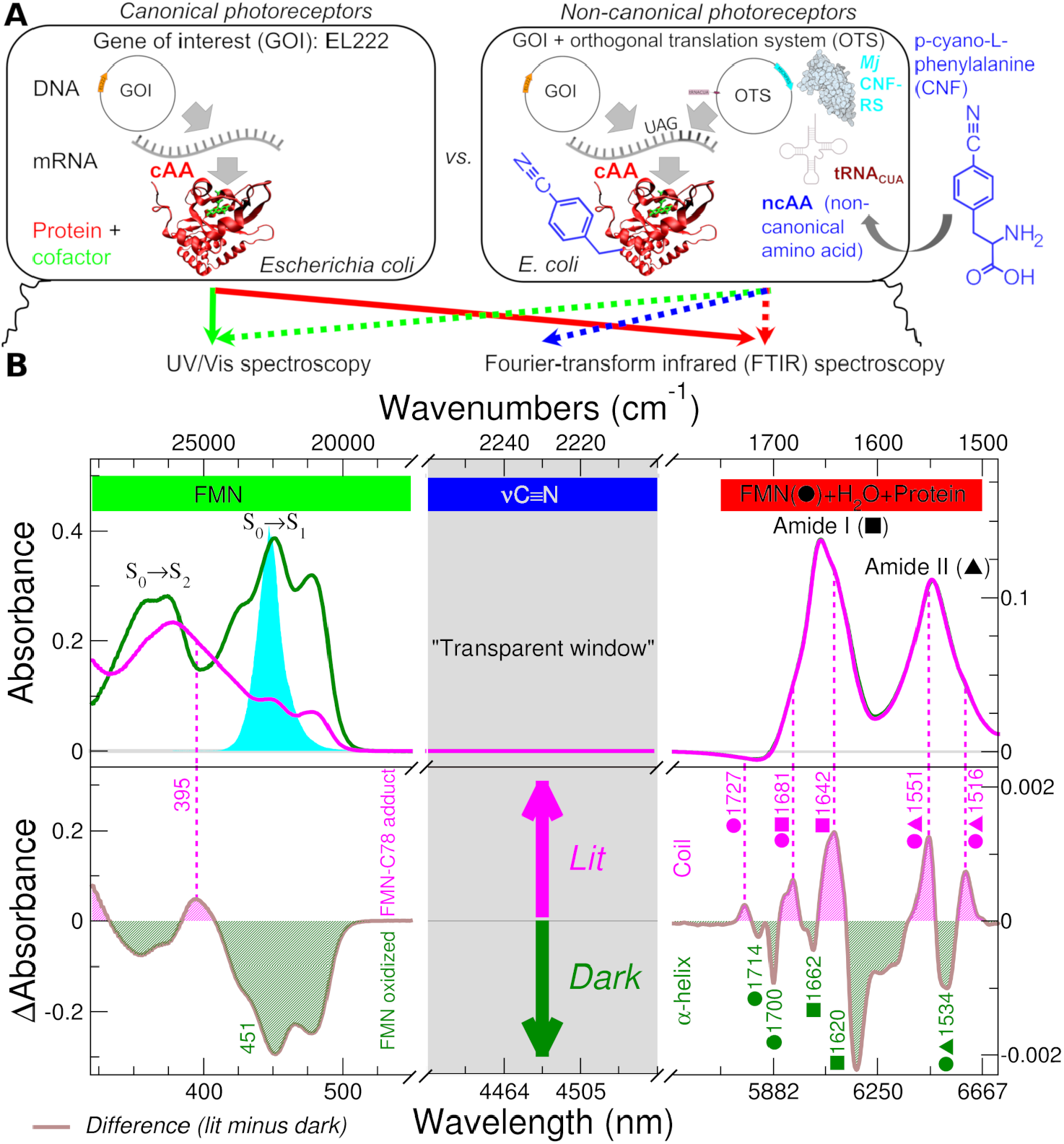
Protein engineering via genetic code expansion meets electronic/vibrational spectroscopy. **(A)** Comparison between photoreceptors based on canonical (left) and non-canonical (right) amino acids (ncAA) carrying vibrational probes. The ncAA p-cyano-L-phenylalanine (CNF) contains a triple bond C≡N group vibrating in the “transparent window” region (grey area in panel B) and delivers local residue-specific information. **(B)** Absolute (top) and difference (bottom) electronic and vibrational spectra of wt EL222 in H_2_O-based buffer. In the case of UV/Vis spectra (left), dark state (green) and lit state (magenta) samples have been collected before and immediately after blue-light illumination (spectral output in the top panel, cyan area), respectively. Transmission IR spectra of hydrated EL222 (right) were collected in the absence (dark state) and presence (lit state) of continuous blue-light illumination. The positions of the main bands discussed in the text are marked as dashed lines and numbers correspond to their wavelength/frequency.

Vibrational spectroscopies are *a priori* ideally suited to monitor protein-specific changes due to their intrinsic sensitivity to chemical bonds and inherently high time resolution, down to the picoseconds time range in the case of infrared experiments (*5, 11*). Light-induced difference infrared (IR) spectra of photosensitive proteins, where the signal from the dark state is subtracted from its lit state, are particularly informative (**Fig. 1B** bottom panel) (*12*). Analysis of the protein amide I band, mostly arising from coupled backbone C=O stretching vibrations (*13, 14*), has revealed secondary structure changes in other LOV photoreceptors, often interpreted as unfolding of the α-helices flanking the core LOV domain (N-terminal A’α and C-terminal Jα) (*15, 16*). In spite of its utility as a reporter for global structure changes in proteins (*13, 14*), the amide I vibration generally lacks spatial detail. Another difficulty is that even IR difference spectra of proteins are strongly congested, with multiple vibrations falling in common frequency regions, making it challenging to ascribe spectral changes to particular residues or vibrations.

Most studies to date have focused on “canonical” photoreceptors composed of the 20 standard amino acids (**Fig. 1A** left panel). As an alternative, chemical groups with absorption frequencies in the range from ∼1800 cm^-1^ to ∼2500 cm^-1^, known as the “transparent window” because most natural proteins do not absorb there (**Fig. 1B** grey area), make the assignment straightforward while providing valuable site-specific information (*17, 18*). These extrinsic non-native moieties, which allow researchers to spectrally single out individual residues, can be incorporated into proteins using a variety of methods (*19*). Genetic code expansion (GCE) technology is arguably the most versatile of such methods as it allows the introduction of non-canonical amino acids (ncAA) at virtually any desired position along the protein sequence (*20*). An engineered aminoacyl-tRNA synthetase recognizes a particular ncAA and attaches it to an orthogonal transfer RNA (tRNA), this one most often including the CUA anticodon (complementary to the amber stop codon, UAG), as illustrated in **Fig. 1A** (right panel). NcAAs can be utilized as chemical handles to introduce reporter groups, or as “spectator” vibrational probes that inform non-intrusively on local changes (*17, 18, 21*). Additionally, the system can be designed so that the included ncAA tunes protein functions and helps elucidating reaction mechanisms (*22*). To date, the three most popular ncAA used as IR reporters have been 4-cyanophenylalanine (CNF) (*23*), 4-azidophenylalanine (*24*), and cyanocysteine (*25*). Thiocyanates, which have the smallest transition dipole strength of the three, are generated by chemical cyanylation of cysteine residues. Azides are potentially unstable due to their chemical and photochemical reactivity (*26*), and their complex absorption profile makes interpretation challenging (*27*). For those reasons we chose CNF, a genetically encodable ncAA featuring a cyano (C≡N) group attached to a phenyl ring at the *para* position (**Fig. 1A** right), as our infrared reporter of local EL222 environments. Being appended to a side-chain, the vibrations of the reporter group are expected to report on local changes in the tertiary and quaternary protein structure, which might differ from global secondary structure changes experienced by the protein and sensed by amide backbone vibrations (*25, 28*). Due to its local nature, a sufficiently large number of tagged positions is imperative for the reporter group to render a complete picture of protein structural changes. However, previous IR studies on photoreceptors decorated with unnatural amino acids have focused mainly on a small number of residues and mainly proximal to the chromophore site (*23-25, 28-30*), with its associated limitations.

Here, we exploit the potential of merging protein engineering tools with spectroscopy by tagging in EL222 up to 45 different residue positions with the unnatural amino acid CNF. We characterized the spectral properties of CNF mutants and mapped them on the 1D (sequence) and 3D (global fold) structural levels of EL222. After a thorough screening phase to identify both intrusive and spectrally silent CNF probes, we choose the most promising EL222-CNF protein variants for kinetic studies by time-resolved UV/Vis and IR spectroscopies. We focused on the less understood transition from the “lit” state (or mixture of states) present under continuous illumination, to the “dark” state of EL222 repopulated in the absence of light. In total, four kinetic events were identified and interpreted in molecular terms. Thus, the joint use of spectral signals from the chromophore, the protein backbone and genetically encoded side-chain infrared probes enabled a multi-probe and multi-site characterization of non-equilibrium protein structural dynamics in EL222, a strategy that may benefit other LOV proteins and photosensory receptors in general.

## Results

### Preparation of EL222 variants and quality control

#### Mutant selection

Ideally, the IR reporters should cover as much of the EL222 sequence as possible while being minimally perturbative. CNF is chemically similar to the canonical amino acids phenylalanine and tyrosine, which can be envisioned as the natural choices for replacement. However, the four native phenylalanines and four native tyrosines of EL222 are all located in the photosensitive LOV domain and are therefore insufficient for a complete understanding of EL222 structural dynamics. Thus, we looked for other replacement candidates distributed in the protein structure. We selected residues both in the FMN binding pocket (**Fig. S2A**), to track changes in the active site, and along the LOV/HTH interface (**Fig. S2B**), to identify and characterize changes between the putative “decaged” (absence of LOV/HTH interactions) and “caged” (presence of LOV/HTH interactions) states of EL222. In total, we chose a set of 45 residue positions along the EL222 sequence. From the N-terminus to the C-terminus we have (**Table S1**): (i) one residue (A18) in the disordered N-terminal domain, (ii) 3 residues in the A’α helix, (iii) 27 residues in the LOV domain, (iv) 5 residues in the linker domain (including the bridging Jα helix), and (v) 9 residues in the HTH domain. Such an extensive labeling corresponds, on average, to one local probe inserted every six residues.

#### Mutant production

For amber suppression in *E. coli* we used an orthogonal translation system (OTS) comprised of an orthogonal aminoacyl-tRNA synthetase/tRNA pair derived from tyrosyl tRNA synthetase of the archaea *Methanocaldococcus jannaschii* (*Mj*CNFRS) and its cognate tRNA (**Fig. 1A** and **Table S2**). The modified suppressor tRNA incorporates the 5’-CUA-3’ anticodon (tRNA_CUA_) that is efficiently recognized by the modified synthetase. The enzyme is also able to attach phenylalanine derivatives (CNF and others) to the suppressor tRNA_CUA_, thus allowing decoding UAG codons during protein translation (*31, 32*). All proteins engineered using this strategy (see sequences in **Table S2**) were purified to > 95%. Mass spectrometry analyses confirmed the successful incorporation of CNF in all 45 single-mutant proteins (**Table S3**). Although the fidelity of MjCNFRS is known to be high, mis-incorporation of phenylalanine is still possible. Mass spectrometry confirmed that mis-incorporation of Phe was only significant in two mutants (W89CNF, I94CNF), where the phenylalanine-carrying EL222 was the most abundant species detected. In these cases, we were able to minimize the presence of Phe by re-expression of these constructs in minimal media (**Table S3**). In another mutant (A42CNF) we only found evidence of truncated products, which cannot be attributed to the UAG codon being recognized as STOP since the purification tag is found at the C-terminus, but to post-translational protein instability enhanced by the introduced mutation, consistently with the known tendency of EL222 to undergo proteolytic degradation under certain conditions (*2*).

While for double-CNF mutants the amount and purity of protein was still sufficient for subsequent studies, both the yield and protein translation fidelity of triple-CNF mutants were unsatisfactory low. This a known issue of nonsense suppression methods and the reason why the large majority of studies limit themselves to typically one or exceptionally two target positions for ncAA replacement. We turned to the *E. coli* strain C321.ΔA.exp(DE3) as the expression host, a genomically recoded organism that supports multiple instances of UAG codon suppression with high efficiency, at least in model proteins (*33*). Gratifyingly, we could purify a few milligrams of high-quality EL222 (**Table S3**) and subject it to further experiments.

#### Initial screening of mutants

Much alike site-directed mutagenesis, the introduction of an unnatural side-chain may give rise to non-native interactions that can alter the structure, thermodynamics and kinetics of the system under study (*34*). Altered mutants need not be discarded since they can provide valuable mechanistic insights, but their interpretation must be done with extreme care. To ascertain the perturbative effect of the introduced CNF residues, we initially monitored two parameters: the level of incorporation of FMN, used to discard gross structural perturbations, and the oligomeric state of EL222 in the dark, to discard constitutive protein-protein interactions in the absence of illumination. Further screening of CNF mutants criteria based on quantifying their degree of similarity with respect to wt EL222 (vide infra) allowed us to improve their classification as intrusive/non-intrusive.

Most of the 44 full-length mutants of EL222 were able to incorporate FMN stoichiometrically (one mole of FMN per mole of protein). However, four mutants did not bind FMN at all (**Table S3**): V44CNF, S46CNF, C78CNF, and N110CNF. The mutations C78CNF and N110CNF involve residues participating in FMN binding (**Fig. S2A**). Indeed, both mutations were originally intended as negative controls: C78 is the critical residue that makes a covalent adduct with the C4a atom of FMN upon blue-light illumination, and N110 makes two hydrogen-bonds with N3 and C2=O2 of FMN that might be critical for the binding of this cofactor (*2*). Additionally, the bulky CNF side-chain, compared to asparagine, would reduce the space available to FMN in the binding pocket in the N110CNF mutant. Although V44 and S46 are not involved directly in FMN binding, the mutation of these small residues by CNF may introduce detrimental steric clashes, probably leading to long-range structure changes that may reach the binding site of FMN and prevent its incorporation. Out of the 40 EL222 mutants able to incorporate FMN, in four cases (L55CNF, N120CNF, V122CNF and V124CNF) we found a significant drop (less than 50%) in FMN loading. Thus, for them, we expect sample heterogeneity in the form of a mixture of apo-EL222 and holo-EL222.

Concerning the oligomeric state of holo-EL222 wt, it is a monomer in the dark but readily oligomerizes upon light stimulation (*3, 4, 6*). We verified the oligomeric state of wt EL222 by size-exclusion chromatography (SEC) and found that its elution volume, related to the particle size, was in agreement with the expected molecular weight of a monomer. Similarly, nearly all mutants were found to exist as monomers in the dark, with similar elution volumes as wt EL222 (**Table S3**). Three notable exceptions were S140CNF, L123CNF and Q141CNF. The elution volume of S140CNF was significantly smaller than the elution volume of wt EL222, suggesting a multimeric organization in the dark. The latter two showed elution volumes in between those expected for monomeric and dimeric assemblies. This behavior is akin to those previously reported for L123K and S140Y mutations, which resulted in the assembly of oligomers (*2, 6*). Hence, our results suggest that the three mentioned mutants (L123CNF, S140CNF and Q141CNF) may partially or fully adopt in the dark a structure resembling the conformation competent for DNA binding adopted by wt EL222 after illumination, i.e., resembling the lit state.

### Steady-state spectroscopy

EL222 samples were measured in the dark as well as under continuous blue-light illumination, achieving a photostationary state. We focused on three spectral regions. The first is the 320-550 nm (UV/Visible) region, which is due to absorption bands of the FMN chromophore (S_0_→S_1_ and S_0_→S_2_ electronic transitions, **Fig. 1B** top left panel). The second is the 1750 to 1500 cm^-1^ mid-IR region which includes absorption bands from both the protein backbone (amide I and II), FMN (C=O and C=C stretching vibrations among others) and water (**Fig. 1B** top right panel). The third region includes the band around 2230 cm^-1^ from the selectively introduced nitrile moieties. Peaks in the latter region are absent in wt EL222 (compare **Figs. 1** and **2**).

#### UV/Visible absorption region of FMN

We first compared the UV/Vis spectra of wt EL222 and CNF mutants to identify those mutations with a preserved FMN spectrum and, by extension, with protein-chromophore interactions similar to those in the wt (**Fig. S3**). We characterized the spectral similarity in the 400-550 nm region of the different mutants with respect to wt by a linear correlation analysis (**Fig. S3A**), which we complemented by determining shifts in maximum absorbance wavelength (Δλ_max_) with respect to wt (**Fig. S3B**). With few exceptions, all variants showed a high spectral similarity (correlation coefficient above 0.9) and Δλ_max_ less than 2 nm, sustaining a preserved environment of the FMN chromophore in the dark. However, three mutants showed clear indications of an altered FMN environment (L55CNF, N120CNF, and Q141CNF), with a correlation coefficient below 0.7 and Δλ_max_ larger than 4 nm.

#### IR absorption region of native protein and FMN

The main band in this region is the amide I band (1700 to 1600 cm^-1^), primary arising from peptide backbone carbonyl C=O stretching modes, known by its secondary structure sensitivity (*35*). The nearby amide II band (1600 to 1500 cm^-1^) is due to C-N stretching and N-H bending modes, which can be used to report on solvent accessibility of the protein backbone via hydrogen/deuterium exchange experiments (*36*). The two carbonyl stretching modes of flavins (C2=O2 and C4=O2, see **Fig. S2A**) also show up in the 1750 to 1650 cm^-1^ spectral region (*8, 37, 38*). Absolute FTIR spectra in the 1750-1500 cm^-1^ region of dark-adapted and lit-adapted wt EL222 in H_2_O are hardly distinguishable (**Fig. 1B** top right). In contrast, lit-minus-dark difference spectra show significant changes between both (**Fig. 1B** bottom right). The positive/negative features at 1727 cm^-1^ (+)/1714 cm^-1^ (-) and 1681 cm^-1^ (+)/1700 cm^-1^ (-) can be attributed to the C4=O4 and C2=O2 stretches of FMN, respectively (*8, 39, 40*). The positive-negative-positive feature at 1551 cm^-1^(+)/1534 cm^-1^ (-)/1516 cm^-1^ (+) is probably due to changes in the protein amide II band, but potential contributions from side-chains (e.g. tyrosine ring vibration) or FMN modes (particularly the C4A-C10 ring stretching vibration, see numbering in **Fig. S2A**) cannot be ruled out (*5, 16, 41*). Even with a clear assignment of the above bands to amide II vibrations, their interpretation in structural terms would be difficult as a result of the complex dependence of amide II frequency on secondary structure (42). The negative bands (bleaches) at 1662 cm^-1^ (-) and 1620 cm^-1^ (-) and the positive bands at 1681 cm^-1^ (+) and 1642 cm^-1^ (+) cm^-1^ could be assigned to the secondary structure-sensitive amide I vibrations from the protein backbone. However, the O-H scissoring mode of liquid H_2_O has a strong background absorbance in the amide I region, potentially distorting the intensity of bands in this region when working in solution as in the present case. The background absorbance from water in the amide I region can be dramatically reduced by dissolving the protein into deuterated water (D_2_O), which also induces the H/D exchange of labile hydrogen atoms in the protein and causes bands from vibrations involving these hydrogens to shift significantly. Absolute and difference IR spectra of wt EL222 in D_2_O are shown in **Fig. S1A**. Although a definitive assignment of all observed bands is out of the scope of this study, some interpretations in the amide I’ (I in H_2_O) region are possible. The negative band (bleach) at 1647 cm^-1^ (-) (1662 cm^-1^ in H_2_O) can be attributed to loss of α-helices, with concomitant positive bands at 1661 cm^-1^ (+) and 1630 cm^-1^ (+) (1681 and 1642 cm^-1^ in H_2_O) reflecting gain in unordered structures and/or β-sheet elements. Thus, from the point of view of secondary structure changes in the protein, IR spectroscopy suggests a light-induced helix-to-coil transition *somewhere* in EL222. In other LOV domains, a similar IR pattern has been interpreted as unwinding/unfolding of the Jα helix (*15, 43*), and unfolding of the A’α helix (*44*). Hence, we assume that in the EL222 protein, the Jα and A’α helices are, individually or in combination, the likely candidates to unfold upon blue-light irradiation.

#### IR absorption region of genetically encoded nitrile groups

We tagged EL222 with cyano probes at different positions to address light-induced structural changes residue-by-residue. In **Fig. 2A** (top panels) we show the absolute spectra of the C≡N stretching vibration of the 40 single-CNF EL222 mutants able to incorporate substantial amounts of FMN. Exceptionally, the EL222-W89CNF mutant showed no measurable peak in that spectral region despite all quality-control analyses confirming the presence of the nitrile moiety (**Table S3**). We characterized the vibration frequency of the C≡N stretching from the peak maximum (ν_max_). We also looked at the band areas, proportional to the integrated extinction coefficient: larger areas indicate a larger transition dipole moment for the C≡N bond. Recently, it has been proposed that this dipole senses the local electric field by a Stark effect and, thus, the average local electric field 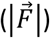 projected on the C≡N direction can be derived from the peak area of the C≡N stretch (*45*). Hence, we finally extracted two parameters from the C≡N band of 39 single-CNF mutants: ν_max_ and 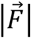 (**Note S1** and **Table S4**). Peak maxima span ∼14 cm^-1^, from 2223 cm^-1^ for N120CNF to 2237 cm^-1^ for Y68CNF, a variability similar to that reported before for nitrile groups in other proteins (∼18 cm^-1^) (*46*). As a reference, for model compounds the nitrile frequency in aprotic solvents decreases with the solvent polarity, from 2233 cm^-1^ in hexane to 2227 cm^-1^ in DMSO. However, for polar protic solvents (able to form H-bonds), the nitrile frequency increases, being 2236 cm^-1^ in water (*47*). Thus, as a first approximation, one could infer that residue CNF68 experiences a very polar and H-bonding environment, and CNF120 a rather apolar and non-H bonding environment. Indeed, CNF120 is the only residue whose peak maxima can be predicted by electric field-frequency calibration curves valid for non-protic solvents (**Fig. S4**) suggesting that all other CNF residues of EL222 are engaged in H-bonding to one extent or another (see further results below).

**FIGURE 2.**
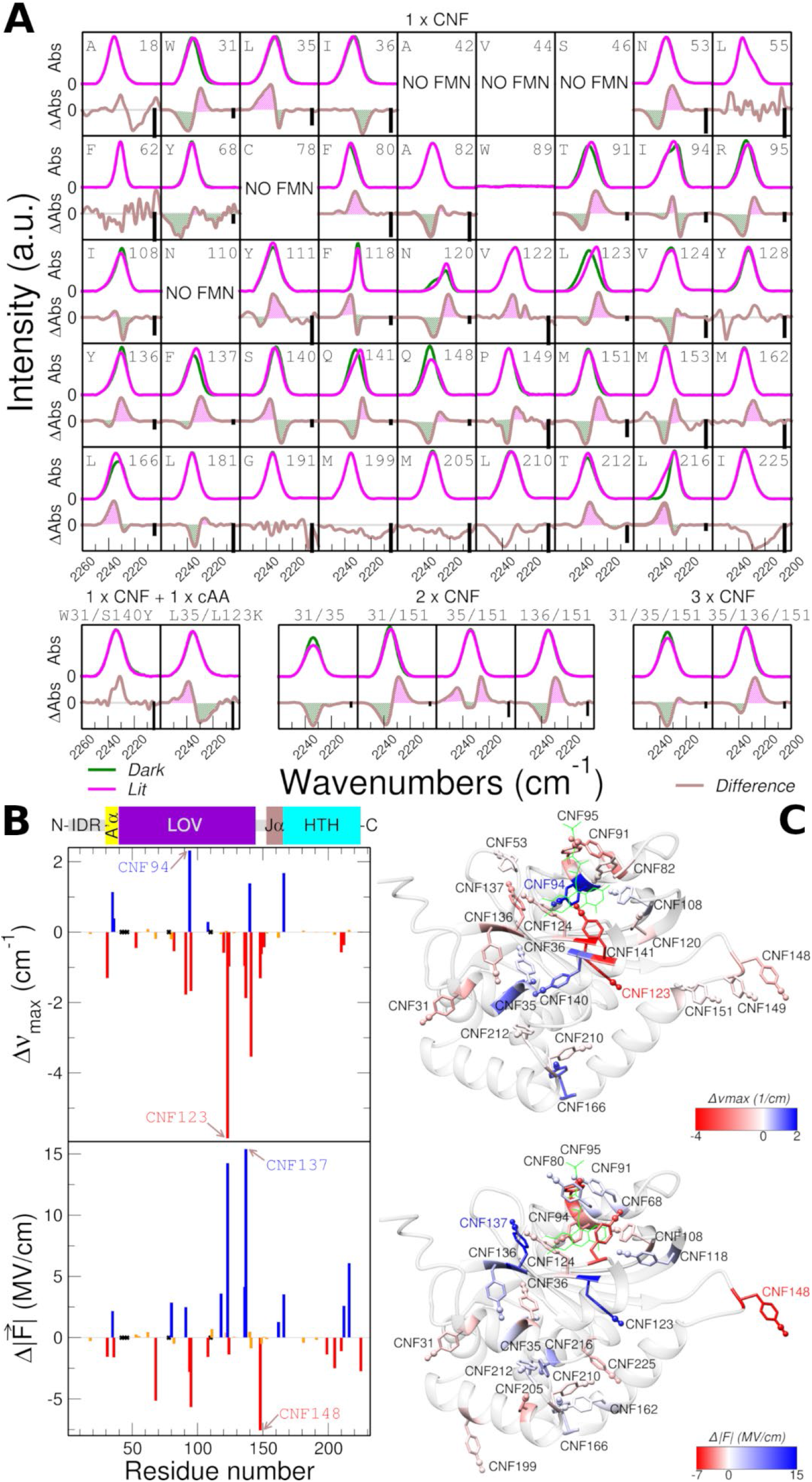
Site-specific infrared spectroscopy of EL222 mutants in the dark and lit states. **(A)** Absolute (top) and steady-state light-induced difference (bottom) IR spectra in the nitrile absorption region of forty-five single-CNF EL222 mutants (each carrying a single CNF ncAA in place of the indicated native residue), together with four double-CNF, two triple-CNF and two single-CNF/single cAA mutants (bottom row). Negative bands in the lit-minus-dark difference spectra correspond to the dark state (green areas) while positive bands indicate the lit state (magenta areas). Difference spectra without shaded areas indicate absence of significant changes (less than 0.25 cm^-1^ frequency shifts and less than 1 % area difference). The vertical scale bar (black) shown in the difference spectra amounts to 0.5 mOD. **(B)** Mapping site-specific changes reported by the CNF probe along the sequence. The light-minus-dark difference between the frequency of the maximum absorbance (Δν_max_) and changes in electric fields 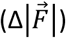 are plotted as a function of the residue number. Residues with the largest differences are indicated. A scheme of different EL222 regions is shown on top, displaying, from the N-to the C-terminus, an intrinsically disordered region (IDR, grey), the A’α extension (yellow), the LOV domain (purple), another IDR (grey), the Jα bridging helix (brown), and the HTH domain (cyan). **(C)** Mapping site-specific changes reported by the CNF probe onto the known 3D structure of dark-adapted EL222 (PDB: 3P7N). Separate models with CNF residues colored according to the two criteria in panel B, Δν_max_ (top) and 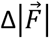 (bottom) are shown.

As done above for the amide I band, a simple procedure to observe changes upon light stimulation is to record a difference spectrum. Difference IR spectra in the nitrile stretching region are shown in **Fig. 2A** (bottom panels) for each individual EL222 mutant. Mutants were classified in three categories based on the observed frequency shift (Δν_max_), and the estimated change in electric field 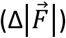 of the nitrile band upon illumination. Sixteen mutants experienced a red-shift in ν_max_, i.e., the illuminated samples had lower mean frequency for its nitrile group after illumination. Six proteins experienced, in contrast, a blue-shift, and seventeen mutants did not show significant frequency shifts (below 0.25 cm^-1^). For both red-shifted and blue-shifted difference FTIR spectra, a simple pattern is typically observed with a single negative band (i.e. the disappearing dark-state) and a single positive one (i.e. the appearing lit-state). One outlier is L35CNF whose difference IR spectra suggest two components in the lit state. Concerning the changes in the electric field between dark (non-irradiated) and lit (irradiated) states, 12 CNF labels sensed a smaller electric field upon illumination, 11 a larger field, and another 16 sensed virtually no change (less than 1 MV/cm difference).

### Visualization and interpretation of CNF spectra

#### Mapping absolute and light-induced CNF-derived shifts

We started first by mapping the characterized CNF microenvironments in the dark state (frequency and electric field) into the primary structure of EL222 (**Fig. S5A**). No evident sequence-dependent pattern was found for the nitrile frequency. However, the larger electric fields tend to concentrate in the middle of the sequence of EL222, between residues 122 and 141 (Hβ and Iβ Strands of the LOV domain). In order to better interpret the spectral changes at the atomic level, we next mapped the nitrile frequencies and electric fields into the dark state crystal structure of EL222 (**Fig. S5B**) (*2*). To this end, we created structural models of the mutants with substitution of the native residues by CNF, considering only the most energetically favored side-chain rotamer (**Note S2** and **Table S5**). Although in these structural models the positions of the cyano side-chains are approximate, it seemed that residues close in space behaved spectrally similar. Exceptions were found. For instance, while CNF148 and CNF151 display similar areas for their nitrile vibration, the peak area of CNF149, and therefore the sensed electric field, is significantly different.

Next, we performed a similar analysis to quantify the extent to which the local environment sensed by the CNF probe changes upon illumination. The light-induced differences in the C≡N stretching wavenumber (Δν_max_), and local electric field 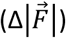 were plotted as a function of the residue number for all the variants (**Fig. 2B**). Significant light-induced IR shifts were observed throughout the EL222 sequence from the N-terminal A’α helix (CNF31) to the C-terminal HTH DNA-binding domain (CNF216). Propagation of conformational changes far beyond the FMN locus has been also proposed based on the NMR chemical shift differences between dark and lit EL222 states (*2*). Our results using the cyano groups have unveiled a “hot spot” between residues 136 and 153, where on average half of the residues showed notable light-induced frequency and/or area shifts. This region encompasses the C-terminal part of the LOV domain (Iβ strand), the intrinsically disordered region between the folded domains, and the N-terminal part of the bridging Jα helix. In other LOV-containing proteins, this part of the β-sheet has been proposed as an interaction interface between LOV domains (*48-50*), and the Jα helix has been shown to unfold upon blue-light irradiation (*15, 43*).

The magnitude of the photoinduced changes in frequency or electric fields (areas) does not correlate with closeness to the light-harvesting site (the FMN, see **Fig. S6**). On the other hand, spatial proximity of residues does not always imply a similar spectral response of the nitrile group of CNF to the protein structural changes induced by light (**Fig. 2C**). For instance, CNF151 and CNF153 (two residues in the Jα helix) experience comparable red shifts in the lit-state without clear area changes, possibly connected to a partial Jα unfolding. However, the three residues sitting on A’α (CNF31, CNF35, and CNF36) respond quite differently in the lit-state: red shift and area decrease for CNF31, blue shift and area increase for CNF35, and blue shift and area decrease for CNF36, thus complicating the interpretation about the nature of light-induced structural changes in the A’α helix. This is in line with the multiple roles played by A’α in other LOV domains, including unfolding (*44*) and participation in protein-protein interaction interfaces (*48-50*).

It is informative to compare our IR results with the published NMR assignments of EL222 (*2*). We should keep in mind that the ^1^H-^15^N chemical shifts report primarily on the environment around peptide backbone amide groups while site-specific IR frequency shifts monitor the environment around side-chain C≡N triple bonds. Additionally, the NMR shifts correspond to the fraction of EL222 molecules that remain monomeric under illumination, while IR is sensitive to all species (monomeric or else) present in solution. Some residues were only accessible to IR (e.g. W31) or NMR (e.g. A42). For some positions both techniques agree and point to changes in the A’α (L35), LOV domain (N120), 1α-2α loop (G181), and 4α helix (L216) of the HTH domain. Other locations show differential sensitivity. For instance, F80 is the residue with the largest chemical shift difference but the nitrile frequency of CNF80 did not vary substantially with light and only a modest 2 MV/cm change in electric field was observed. On the other side, IR clearly detects a change in the surroundings of residues CNF118 (LOV domain) and CNF166 (HTH domain), that is largely absent in NMR studies for the corresponding native residue F118 and L166. These results emphasize the complementarity between NMR and IR spectroscopies for the interpretation of structural dynamics in complex biological machines.

#### Towards a molecular-level interpretation of nitrile frequencies

Introducing CNF groups at many different positions in EL222 provided us with information about light-induced changes on the local electric field from variations in the nitrile band area. We also resolved changes in the nitrile frequency, whose interpretation is more challenging by its dual dependence on local electric fields and H-bonding (*45*). In H-bonding environments, e.g. water, a blue-shift of the nitrile stretching frequency upon increasing the magnitude of the electric field is observed (*45, 47*). In contrast, in non-H bonding environments, e.g. DMSO, the nitrile stretching frequency red-shifts as the electric field increases (*47*). A combined analysis of peak frequency and band area of nitrile probes has been recently proposed to characterize their H-bonding (**Note S1**) (*45*). We applied this analysis to our data, and the estimated frequency shifts ascribable to H-bonding (Δν_HB_) are listed in **Table S4** for each CNF group. Surprisingly, little or no significant changes in the H-bonding strength of nitrile groups are predicted between dark and lit states of EL222 by such analysis, CNF123 being one notable exception. Our results do not dismiss the utility of such a method but rather suggest that further benchmarking is necessary.

Another more established way to identify changes in the hydrogen bonding interactions of nitrile groups, even in the dynamic and complex environment found in a protein (*46*), is through frequency-temperature line slope (FTLS) plots (*51*). In FTLS analysis, the vibrational frequency of a selected bond is recorded as a function of temperature. For model nitrile groups dissolved in aprotic solvents, unable to form H-bonds with the nitrile group, the slope is zero. In contrast, when dissolved in water, a protic solvent, the slope is approximately -0.04 cm^-1^/°C (equivalent to ∼1 cm-^1^ downshift every 24 °C increase) (*46*). Thus, the slope in FTLS plots for CNF groups is related to the degree of exposure to water, with a zero slope indicative of a buried residue (*51*). Even more, the slope is directly related to molecular-level details of the specific hydrogen bonding interactions to the nitrile probe. In particular, for a given CNF residue there is a strong correlation between the FTLS and the solvent accessible surface area, a structure-sensitive parameter (*46*). A disadvantage of this method is that it requires spectral measurements at different temperatures and is therefore time-consuming when applied to a large number of proteins.

We concentrated on four positions that cover different parts of the sequence, show wt-like FMN spectra and display clear light-induced spectral differences: W31CNF (A’α, red-shift and smaller electric field), L35CNF (A’α, blue-shift and larger electric field), Y136CNF (LOV, red-shift and larger electric field), and M151CNF (Jα, red-shift and no change in electric field). The experimentally measured spectra in the nitrile absorption region of several EL222 variants is plotted in **Fig. S7**. The chosen temperature range (5-35 °C) is well below the melting temperature of wt EL222 (45 °C). Two residues (Y136CNF and M151CNF) show relatively small but clear changes in slope between their dark and lit states (**Fig. 3A**). In the absence of illumination, CNF151 displays a slope indicative of full solvent exposure, while for CNF136 the slope lies in between that expected for a fully exposed and a fully buried residue. Both residues become less exposed to water upon perturbation by light. On the other hand, a large change in slope occurring between the dark and lit states of EL222 W31CNF (**Fig. 3B**) indicates a dramatic change in polarity around the residue 31, from a slope corresponding to full solvation in the dark state to a null slope indicative of a solvent-excluded residue in the lit state. Since EL222 is known to oligomerize in the lit state (*3, 4, 6*), we suggest that CNF31, and by extension W31, participates in a protein-protein interaction surface. In fact, the dimer interface seen in the X-ray structures of three LOV photosensors crystallized in the lit state is largely constituted by interactions involving the A’α helix (*48-50*). To test such hypothesis we designed two EL222 mutants as positive (constitutive oligomerization) and negative (diminished oligomerization) controls. First, we prepared the S140Y mutation, known to promote EL222 oligomerization already in the dark-adapted state (*6*), in combination with W31CNF as a reporter. Wt EL222 was nearly 100% monomeric without light stimulation (**Table S3**). We found that the EL222 W31CNF/S140Y mutant had enhanced oligomerization propensity in the absence of illumination with respect to the wt version (**Table S3**), and its light-induced difference IR spectra in the C≡N region did not show appreciable changes (**Fig. 2A**). Moreover, FTLS analysis of W31CNF/S140Y displayed a reduced slope compared to W31CNF, suggesting a lower degree of solvent accessibility around CNF31 in the dark state of the EL222 W31CNF/S140Y variant compared to the dark state of EL222 W31CNF, albeit not as dehydrated as the lit state of EL222 W31CNF (**Fig. 3B**). As a second control, we produced a truncated EL222 lacking the HTH domain, known to drive homo-dimerization (*2*), again including the mutation W31CNF as a reporter (LOV-W31CNF). This variant was mostly monomeric in the dark (**Table S3**) and showed no clear light-induced spectral changes (**Fig. S7**). Accordingly, the slope in the FTLS experiments were similar between LOV-W31CNF and the parental two-domain EL222 W31CNF mutant in the dark state, suggesting a similar environment around residue CNF31 (**Fig. 3B**). Taken altogether, these results are consistent with the nitrile moiety at position 31 reporting primarily on the oligomerization state of EL222.

**FIGURE 3.**
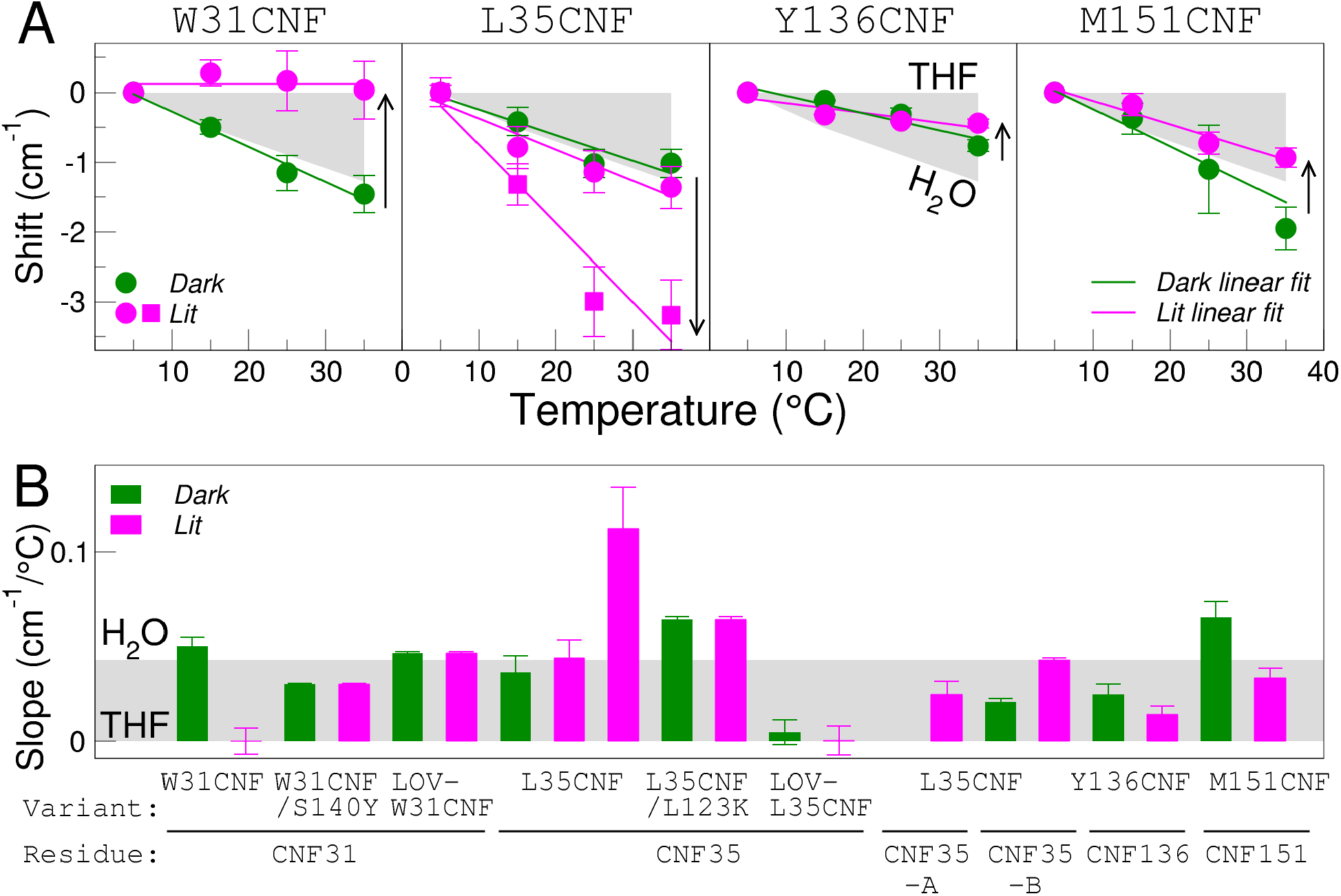
Light-induced changes in solvent exposure and H-bonding around selected CNF residues. **(A)** Frequency-temperature line slope (FTLS) analyses of the C≡N absorption bands of four single-CNF EL222 variants (indicated on top of the panels). Frequency shifts, calculated with respect to spectra acquired at a temperature of 5 °C, of non-illuminated and illuminated samples are indicated as green and magenta circles, respectively. In the EL222 L35CNF panel, the shifts of the additional lit species found for this mutant are represented as magenta squares. Arrows indicate the magnitude of the change in site-specific hydration for the dark-to-lit transition. **(B)** Slopes calculated from linear fits to FTLS plots for the indicated EL222 variants and CNF residues. EL222 W31CNF/S140Y is a constitutively oligomeric variant (A’α-LOV/A’α-LOV interactions in the absence of illumination). EL222 L35CNF/L123K is constitutively “open” variant (lack of interaction between A’α-LOV and DNA-binding domains in the absence of illumination). LOV-W31CNF and LOV-L35CNF are both truncated versions of EL222 lacking the C-terminal DNA-binding domain. CNF35-A and CNF35-B are the “fast” and “slow” kinetic components of the cyano relaxation of L35CNF variant (see **Fig. 5**). As a reference, the grey area represents the behavior of free CNF dissolved in tetrahydrofuran (THF, zero-slope) versus water (H_2_O, negative slope of 0.043 cm^-1^/°C). Raw data can be found in **Fig. S7**. Error bars represent the standard deviation of 3 independent measurements.

In the case of the L35CNF mutant, the observed asymmetry in the C≡N band (**Fig. 2A**), possibly indicating the presence of more than one CNF environment, complicated the analysis. Indeed, three bands (one negative and two positive) were clearly resolved at all temperatures in the photostationary light-minus-dark difference IR spectra of EL222 L35CNF (**Fig. S7**). In principle, two bands in the absolute spectra should give rise to two pairs of positive/negative bands in the difference spectra. Hence, the missing band could be possibly due to cancellation of one negative band by others. We fitted multiple Gaussian distributions to the difference spectra of L35CNF, and used the mean peak positions to construct FTLS plots (**Fig. S7**). The slope of the negative band (corresponding to the dark state) agrees with a fully water exposed residue. The slope of one of the positive bands was similar to the slope of the dark state, while the other positive band had a steeper slope, more than twice that reported before for model CNF groups dissolved in water (**Fig. 3B**). Therefore, we conclude that in one of the two environments (or conformations) of L35CNF populated in the light, the CNF probe experiences a stronger hydrogen-bonding environment than in liquid water. Because L35 resides in A’α element (residues 30-37) and its side-chain lies at the interface between the A’α and HTH domains (**Fig. S1B**), we asked ourselves whether CNF35 could be sensing the folding status (helix *vs*. coil) of A’α or the proximity between the A’α-LOV and HTH halves of EL222. To test the latter hypothesis, we designed two EL222 variants as negative controls. First, we prepared a double mutant containing L35CNF and L123K. The latter mutation has been reported to increase the size (hydrodynamic radius) and proteolytic sensitivity of EL222, probably by favoring an “open” conformation (HTH decaged from the LOV domain) even in the absence of illumination (*2*). The elution time of L35CNF/L123K was intermediate between that of monomeric and dimeric wt EL222 (**Table S3**), compatible with monomeric EL222 in a more extended conformation than that seen in the crystal structure. The absolute and difference IR spectra of EL222 L35CNF/L123K are shown in **Fig. 2A**. The amplitude of the difference spectrum was too low and noisy to allow further analysis. Nevertheless, FTLS analysis of L35CNF/L123K rendered a slope different from that seen in the dark state of L35CNF (**Fig. 3B**). Secondly, we introduced the L35CNF mutation in an EL222 variant lacking the HTH domain (LOV-L35CNF). There was evidence of only a single lit environment for residue CNF35 in LOV-L35CNF (**Fig. S7**), which did not differ significantly from the dark-state (**Fig. 3B**). We conclude that in one of the two environments experienced by CNF35 in the EL222 L35CNF mutant, the CNF probe might be sensing the light-induced open/close (decaged/caged) conformational equilibrium of the protein.

### Time-resolved spectroscopy of EL222

To characterize the transition from lit to dark states with single-residue resolution, we measured the photorecovery kinetics of EL222 using three different spectral regions: 320-550 nm (FMN probe), 1500-1750 cm^-1^ (amide probe), and 2200-2270 cm^-1^ (CNF probe) (**Fig. 4A**). The UV/Visible spectra mainly reveal adduct rupture (A_390_-to-D_450_ transition), bands in the 1500-1750 cm^-1^ region disclose global protein secondary structure changes (Amide_coil_-to-Amide_helix_ transition) among others, and cyano bands in the “transparent window” region report on local environment changes around a particular CNF residue (CNF_lit-environment_-to-CNF_dark-environment_ transition) (**Fig. 4B**). Thus, as opposed to FMN and amide bands, which report on local changes around the chromophore and global protein backbone changes, respectively, the nitrile group witnesses local changes around the C≡N group of the CNF residue to which it is attached. Of note, a direct comparison of all these three probes can only be accomplished using H_2_O as a solvent. While D_2_O as a solvent facilitates the study of the structure-sensitive amide I region, as already mentioned, it literally wipes out the tiny infrared signal from the nitrile stretching vibration (**Fig. S1A**). Measuring secondary structure changes of soluble proteins in bulk H_2_O is possible but technically challenging by the strong background absorption from H_2_O (see **Note S1**). Prior to studies with CNF mutants, we thoroughly studied the recovery kinetics of wt EL222, natively devoid of nitrile bands, by UV/Vis/IR spectroscopies using two intrinsic probes: FMN and amide.

**FIGURE 4.**
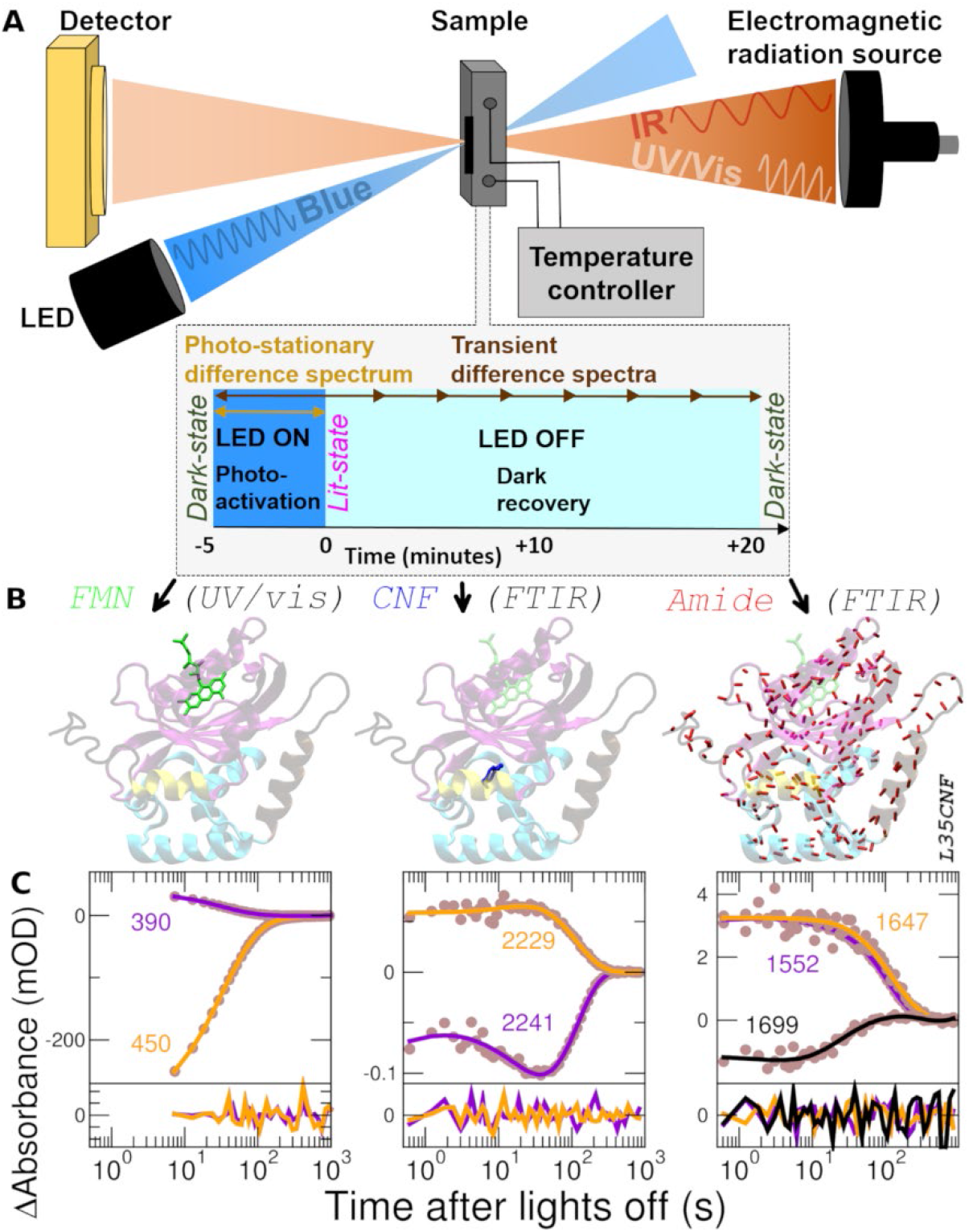
Time-resolved spectroscopy of the lit-to-dark transition of EL222 using multiple probes (FMN, CNF, amide). **(A)** Data were recorded in solution (typically H_2_O-based buffer but a few experiments were conducted with D_2_O-based buffer) at 20 °C (except for the Arrhenius plots) by first accumulating the lit state population under continuous blue-light irradiation (5 minutes at 25 mW/cm^2^), then switching lights off (time zero) and recording rapid-scan spectra for 10 minutes. The same protein was subjected to three independent experiments. **(B)** *Left*, local chromophore-specific time-resolved UV/Vis spectra (FMN probe, green) were taken between 320 and 550 nm in a cell of 10 mm path-length. *Middle*, local residue-specific time-resolved IR spectra (CNF probe, blue) were taken between 2270 and 2200 cm^-1^ with a cell of 50 μm path-length. *Right*, global protein time-resolved IR spectra (amide probe, mostly backbone carbonyls, red) were taken between 1750 and 1500 cm^-1^ (this region includes also contributions from FMN and other vibrations, see main text) with a cell of 8 μm path-length. Molecular models are color-coded as in the scheme of **Fig. 2B. (C)** Kinetic traces of differential absorption (circles) and fits (lines) at two or three selected frequencies as a function of time after ceasing illumination for the three probes (EL222-L35CNF mutant): FMN (left), CNF (middle), and amide (right). Fits were done by maximum entropy lifetime distribution method. Fit residuals are shown in the bottom panels.

#### WT EL222

Full kinetic traces for wt EL222 in H_2_O and D_2_O buffer (wavelength or frequency *vs*. time) can be found in **Fig. S8-S9** (panel A), respectively. Experimental traces were analyzed by a maximum entropy lifetime distribution analysis (LDA) (52). The commonly employed global exponential fit and related kinetic analyses use a fixed number of exponential terms with adjustable but common time constants, which implicitly assume a discrete and known number of intermediary states (*53*). In contrast, LDA uses a distribution of exponential terms, able to adapt to more complex scenarios such as when the number of intermediates is unknown or even not well-defined, when the exponential terms are not discrete but distributed, or when the same time constants are not shared at all frequencies (*54*). An example of experimental and fitted time traces can be found in **Fig. 4C**. The output of LDA is typically a lifetime density map (**Fig. S8-S9** panel B) where each point represents the exponential amplitude for a given lifetime and frequency (either wavenumber in the case of IR or wavelength in the case of UV/Vis spectroscopy). One way to summarize the information content of a lifetime density map is to calculate the square root of the sum over the squared amplitudes for all frequencies, providing a lifetime distribution known as the “average dynamical content”, hereafter abbreviated as *D* (**Note S1**) (*52, 55*). The amplitudes of *D* lifetime distributions provide a visual summary about the magnitude of the spectroscopic changes that occur at a given time constant. *D* lifetime distributions for the UV/Vis and IR data are plotted in **Fig. 5A** (top panel) and lifetimes (τ) with the largest *D* values are listed in **Table S6**.

**FIGURE 5.**
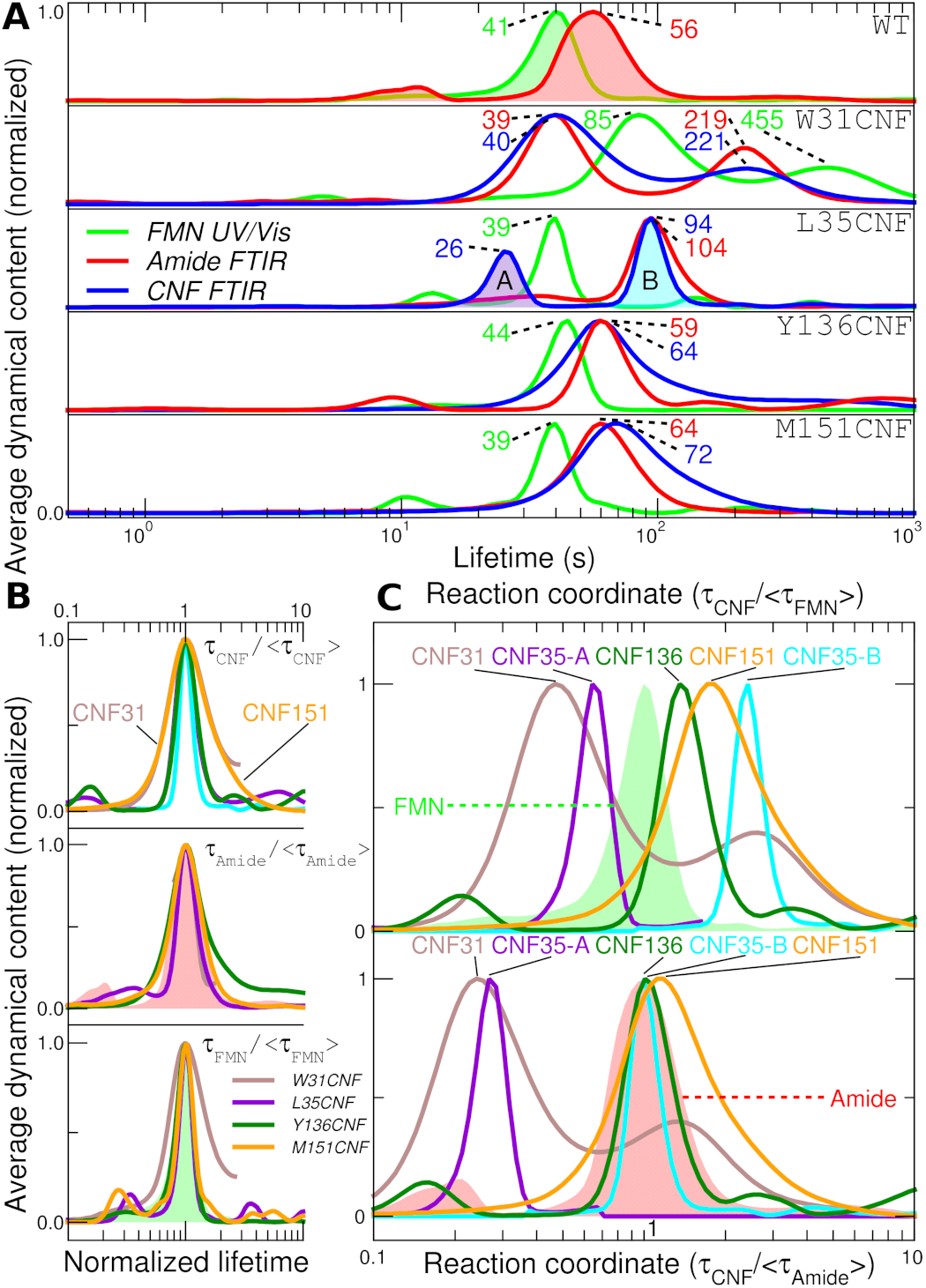
Lifetime distribution analysis (LDA) of the dark recovery kinetics of single-CNF EL222 variants using the maximum entropy method. **(A)** Normalized average dynamical content (*D*) as a function of lifetime for five variants (wt EL222 and single-CNF W31CNF, L35CNF, Y136CNF, and M151CNF mutants) using three probes: FMN (green), amide (red) and the cyano moiety of the CNF residue (blue). The numbers report the lifetimes of the main dynamical events (mean peak value) again color-coded as green, red and blue for FMN, amide, and CNF probes, respectively. Two clear cases of bimodal lifetime distributions are found for the CNF relaxation kinetics of L35CNF, named CNF35-A (violet-shaded area) and CNF35-B (cyan-shaded area). **(B)** Normalized average dynamical content (*D*) as a function of normalized lifetime calculated as the ratio between a given lifetime and its mean value (peak maxima). The top, middle and bottom panels show the CNF-, amide- and FMN-derived distributions, respectively. CNF31 and CNF151 stand out as the moieties whose distributions have the largest width. **(C)** Normalized average dynamical content as a function of the reaction coordinate calculated as the ratio between a given CNF-derived lifetime and the mean peak value of FMN-derived *D* lifetime distribution (top panel), or the mean peak value of amide-derived *D* lifetime distribution (bottom panel). Mean peak values for all *D* lifetime distributions can be found in **Table S6**. For easy referencing, the *D* lifetime distributions of the amide and FMN native probes of wt EL222 are shown as filled areas in red and green colors, respectively.

Regarding the FTIR data, we also obtained *D* lifetime distributions for three spectral sub-ranges: a) 1750-1680 cm^-1^, which is dominated by the two bands from FMN carbonyls, b) 1680-1600 cm^-1^, with bands mainly ascribable to the protein backbone amide I mode, and c) 1600-1500 cm^-1^, which has mixed contributions from the protein backbone amide II bands and FMN ring modes. For wt EL222 in H_2_O, one dominant photorecovery time constant was found for all triplicates at 41 s for UV/Vis (320-550 nm), 44 s for FMN C=O (1750-1680 cm^-1^), 47 s for the Amide II (1600-1500 cm^-1^), and 59 s for the Amide I (1680-1600 cm^-1^) (**Fig. S10A** top). Although the recovery lifetime in the UV/Vis clearly preceded that in the amide I region, it might be unclear if this was a reliable feature or induced by the higher noise and baseline fluctuations in the latter region caused by the strong background absorbance from liquid water. To resolve this point, as well as to gain more insight into the mechanism of dark reversion, we repeated the kinetics in D_2_O-based buffer, as a function of temperature and protein concentration.

In D_2_O-based buffer, dark recovery kinetics of wt EL222 was clearly slower than in H_2_O-based buffer (**Fig. S10A** bottom). The kinetic isotope effect (KIE) for the main kinetic event was 4.6 for the amide II, 4.3 for the FMN, 4.2 for the FMN C=O, and 3.8 for the amide I. Importantly, the decoupling between UV/Vis-based (320-550 nm) and FTIR-based (1750-1500 cm^-1^) kinetics is confirmed in D_2_O buffer, with FMN recovering to the dark state in ∼178 s and the protein backbone in ∼220 s (**Fig. S10A** bottom): FMN recovery occurs prior to protein backbone recovery also in D_2_O solvent.

From recovery kinetics measured in the UV/Vis and IR regions at various temperatures, we prepared Arrhenius plots (natural logarithm of rate constant *vs*. reciprocal temperature). As shown in **Fig. S11A**, the rate constants were basically independent of the IR probed frequency range at all tested temperatures. Therefore, from here on, *D* lifetime distributions and associated lifetimes (τ_Amide_) will refer to the whole 1750-1500 cm^-1^ region. The recovery kinetics in the UV/Vis region was faster i.e. smaller lifetimes (τ_FMN_) than in the IR region at all temperatures, both in H_2_O and D_2_O, with the difference being reduced as the temperature increased, both converging at 35 °C (**Fig. S11B**). The activation energies from the IR and UV/Vis regions, derived from the slopes in Arrhenius plots, were similar but not identical (∼18 kcal/mol and ∼15 kcal/mol, respectively), both H_2_O and D_2_O (**Table S7**). We conclude that acquisition of secondary structure and adduct breakage are two separate events.

Due to the large difference in extinction coefficients and concentrations between the FMN chromophore, peptide carbonyls groups, and nitrile groups, together with the necessity to minimize the background absorption of liquid water in the IR, we had to use lower protein concentrations in UV/Vis (50 μM) than in FTIR (1-2 mM) experiments. Because such difference in concentration could be partially behind the faster recovery kinetics measured in the UV/Vis than in the IR, we measured the dark recovery kinetics of wt EL222 at various protein concentrations by both UV/Vis and FTIR time-resolved spectroscopy (**Fig. S12**). A concentration-independent time constant of ∼40 s was found in the 320-550 nm region from 10 μM to 400 μM, while a concentration-independent time constant of ∼55 s was obtained in the 1750-1500 cm^-1^ region from 400 μM to 2 mM. Therefore, the recovery reaction of EL222 is approximately zero^th^-order from 10 μM to 2 mM. We also confirm that the kinetic parameters derived by UV/Vis and FTIR spectroscopies can be safely compared to each other even if derived from experiments at dissimilar protein concentration, and we reaffirm the conclusion that the recovery of the FMN chromophore monitored by the UV/Vis is faster than the recovery of the protein backbone monitored by IR spectroscopy.

Since protein domains are independent folding units, the observed lit-to-dark refolding kinetics of EL222 could be affected by the presence of two domains (LOV and HTH). We thus prepared and measured a variant (EL222-LOV) containing the A’α and Jα extensions flanking the LOV domain but lacking the C-terminal DNA-binding module. The steady-state difference spectra of EL222-LOV (**Fig. S1B** in H_2_O and **S1C** in D_2_O) were similar to that of wt EL222 suggesting that most of the secondary structure changes are localized in the A’α-LOV-Jα region of EL222. In addition, a similar lit-to-dark recovery kinetics is still present, with FMN relaxation clearly preceding the relaxation of the protein backbone, although with the difference that the rates were accelerated for both H_2_O and D_2_O experiments (**Fig. S10B** and **Table S6**). Therefore, we dismiss the assumption that the asynchronous relaxation of FMN and the protein backbone in EL222 could be a consequence of EL222 being a multi-domain protein.

All these results taken together, three key conclusions can be extracted from time-resolved studies using the native chromophores present in wt EL222 and its LOV domain. The first is that recovery kinetics manifest a single dominant kinetic process, although additional but minor events cannot be excluded. The second is that the main kinetic event reported by UV/Vis spectroscopy (FMN-C78 adduct rupture) seems to precede the main kinetic event probed by IR spectroscopy (global protein refolding to the dark-state conformation), at least at temperatures below 35 °C (**Fig. 5A** top row). The third is that sub-dividing the 1750-1500 cm^-1^ IR region into smaller zones does not lead to a clear increase in the information content i.e. the number of kinetic components stays unchanged. We next asked whether the non-native CNF probes could resolve additional kinetic events in the lit-to-dark transition of EL222.

#### Single-CNF EL222 mutants

To facilitate the kinetic comparison among the CNF mutants, as well as with respect to wt EL222, it is desirable that the introduced CNF residues minimally alter the thermodynamics and kinetics of the photochemical reaction. To this end, we first screened single-CNF mutants according to their photoconversion yields and FMN photorecovery kinetics.

The fraction of EL222 molecules in the lit-state (the adduct state or A_390_ species) upon illumination at 450 nm was estimated based on UV/Vis absorbance measurements (**Note S3**). For wt EL222, the photoconversion yield was close to one (**Table S8**), indicating that EL222 molecules covalently bound to FMN via C78 predominate under steady-state conditions. Photoconversion yields were also large for essentially all mutants (except F80CNF), suggesting that CNF residues minimally perturbed the equilibrium composition of the photostationary state.

The FMN recovery lifetimes for single-CNF mutants, obtained from the main components in the lifetime *D* distribution from time-resolved UV/Vis spectroscopy, are collected in **Table S6**. Almost all variants feature a single kinetic component. One important exception is the W31CNF mutant. Out of the 39 CNF mutants (excluding FMN-less and CNF-less variants), 14 mutants displayed recovery times similar to the wt (∼41 s); 13 mutants were found to recover significantly faster than the wt; and 12 of the mutants returned to the dark state at much lower rates than the wt **Fig. S13A**). The fastest mutant was found to be F62CNF (∼8 s) while the slowest mutant was N120CNF (∼370 s), suggesting a strong destabilization of the lit-state in the former case and a strong stabilization of the lit-state in the latter case. To the best of our knowledge, N120CNF is the slowest single mutant of EL222 so far described and may be useful in optogenetic applications (*56*). Strikingly, five of the “fast” mutations are found in the DNA-binding domain, far away from the FMN (**Fig. S13B**). These results add to the number of studies reporting on the extreme sensitivity of FMN recovery times to even minute perturbations in the LOV folding status and raises concerns about how such perturbations are allosterically coupled to the, often distant, FMN site (*9, 57*).

From the pool of mutants that are kinetically non-perturbative (similar FMN lifetimes as wt), nine of them (L35CNF, N53CNF, I108CNF, Y136CNF, P149CNF, M151CNF, M153CNF, L181CNF, and L216CNF) show a clear frequency shift in the CNF stretch region upon illumination and could therefore be suitable for time-resolved IR studies. Unfortunately, the signal-to-noise ratio of the CNF difference spectra of L108CNF, P149CNF, M153CNF and L181CNF was too low to allow accurate time-resolved measurements. As a result, we performed kinetic analyses with five mutants (L35CNF, N53CNF, Y136CNF, M151CNF, and L216CNF). Exceptionally, we included also the W31CNF variant because, although the kinetics of its FMN group is perturbed, the kinetics of its nitrile group may be sensitive to oligomer dissociation, as we concluded in static FTLS experiments (see above).

We studied the dark recovery kinetics of the six selected EL222 CNF mutants using two different IR frequency ranges: 1750-1500 cm^-1^, plus 2270-2200 cm^-1^ (see **Fig. 4C** for the case of L35CNF mutant). Complete kinetic traces for the six single-CNF mutants can be found in **Fig. S14-S19** (panel A). Most single-CNF mutants displayed a single dominant component in the CNF-derived *D* lifetime distributions, equivalent to a mono-exponential behavior, which was used to calculate the recovery lifetime of the CNF moiety (τ_CNF_) (**Fig. 5A** and **Table S6**). In these cases, the transient decay-associated difference spectra were similar to the steady-state difference spectra (**Fig. S20A** for the nitrile absorption region and **Fig. S20B** for the amide bands). Unlike wt EL222, two mutants (N53CNF and L216CNF, **Fig. S21**) showed a similar recovery kinetics for all three probes (FMN, amide and CNF), suggesting a mechanistic perturbation induced by the CNF label. For the remaining four CNF variants (W31CNF, L35CNF, Y136CNF, M151CNF), the FMN recovery happens before the protein backbone recovery, similar to the behavior observed for wt EL222 (**Fig. 5A** and **Table S6**).

##### Two mutants showed particular behaviors: W31CNF and L35CNF

EL222 W31CNF featured the most intricate kinetics of all single-CNF variants, with two events clearly distinguishable by all three probes, although with unequal amplitudes (**Fig. 5A**). For UV/Vis and CNF relaxation there was a major kinetic event, while for amide-derived kinetics the two components had similar amplitudes. Indeed, the slowest relaxation event of CNF31, which has also the lowest amplitude, was not resolved at all temperatures, so it will be ignored in the following. The spectrum from the slowest kinetic event of W31CNF resembled more the spectrum of W31CNF under photostationary conditions (**Fig. S20B**). The temperature-dependence of the main relaxation component of CNF31 showed non-Arrhenius kinetic behavior, in sharp contrast to all other dynamical processes (**Fig. S11C**), suggesting that CNF31 monitors a unique kinetic event. Therefore, it seems that in the W31CNF mutant, apart from protein backbone and FMN relaxations, an additional kinetic event is taking place and is sensed by both the amide and CNF probes. In combination with the previous mutagenesis and FTLS results, we propose that such an extra process reports on the monomer/oligomer equilibrium of EL222.

The EL222 L35CNF mutant showed two similarly intense dynamical events linked to the relaxation of CNF35 residue, with associated lifetimes of ∼26 and ∼94 seconds (labeled “A” and “B”, respectively in **Fig. 5A** middle). The slowest component correlates with the protein backbone recovery, while the FMN recovery sensed by UV/Vis largely precedes it. On the other hand, the fastest CNF component takes places before the FMN moiety relaxes. To facilitate the interpretation of the two dynamical processes of CNF35, CNF35-A as the fastest and CNF35-B as the slowest, we looked at their transient spectra. The two events have distinct associated spectra and thus report on dissimilar changes around the CNF residue: a red shift from ∼2241 to ∼2230 cm^-1^ for the fastest component, and a blue-shift from ∼2229 to ∼2240 cm^-1^ for the slowest one (**Fig. S20A**). The transient difference spectrum of CNF35-B resembles more the steady-state difference spectrum of the whole CNF population, where gain in α-helicity occurs, than the transient difference spectra of CNF35-A (**Fig. S20A**). Because the CNF35 label is located in the A’α helix, we previously concluded from FTLS analysis of steady-state difference spectra that one of the two spectral components of L35CNF could be monitoring the decaged (“open”)/caged (“close”) conformational equilibrium of EL222 but the folding state of A’α could not be ruled out. To link stationary and kinetic evidence, we performed time-resolved experiments at different temperatures. First, we did FTLS analyses on transient decay-associated difference spectra (DADS) obtained upon lifetime distribution analysis (**Fig. S7**). The main positive band in the DADS of CNF35-B had a similar FTLS slope as one of the lit states of L35CNF, while the main positive band of the DADS of CNF35-A had a flatter slope (lower solvent exposure) than either state of L35CNF (**Fig. 3B**). Thus, the positive band in the “slow” component (CNF35-B) may correspond to one of the lit environments previously found for CNF35 under continuous illumination. On the other hand, hydrogen bonding around the CNF environment corresponding to the “fast” kinetic component (CNF35-A) seems to deviate from the environment sensed under photostationary conditions. Secondly, we prepared Arrhenius plots for the two kinetic events (**Fig. S11D**) and found that the CNF35-B component had similar activation energy as the backbone relaxation calculated from the amide region of wt EL222 but 1.2 times higher than CNF35-A (**Table S7**). Therefore, we conclude that the slowest CNF35 component of L35CNF (CNF35-B) reports on the same (or similar) relaxation event as the backbone amide vibrations i.e. the global increase in α-helical secondary structure content. In contrast, the fastest CNF35 component of L35CNF (CNF35-A) monitors another dynamical process. Although the precise nature of such a process is difficult to define, considering all the mutagenesis and FTLS results, we tentatively propose that CNF35-A could be sensing the interaction between A’α-LOV and HTH halves of EL222 but not the folding state of A’α.

We gained additional information about how homogenous the relaxation kinetics was for the different mutants through comparison of width of the distribution of *D* values, appropriately normalized by dividing the time constant axis by the mean lifetime of a probed range. In such a way, all probes were centered at 1, thus facilitating their comparison when the absolute lifetimes differ. As can be seen from **Fig. 5B**, the lifetime distributions of the nitrile groups in the W31CNF and M151CNF mutants were wider than the rest. Such an effect is unlikely to be an artefact, since all data sets were processed alike and we used essentially the same regularization values in order to obtain the lifetime distributions (**Table S9**). It might rather suggest a heterogeneity in the kinetic barrier for the dark relaxation of CNF31 and CNF151 residues. Actually, non-Arrhenius kinetic behavior was observed for both CNF31 and CNF151 (**Fig. S11D** and **S11E**) in contrast with UV/Vis- and amide-derived kinetics. In the particular case of CNF151 we found a nearly zero activation energy at temperatures beyond 15 °C thus suggesting a barrierless transition between its lit and dark environments (**Table S7**).

To compare the temporal evolution among mutants, given the altered and complex kinetics in some cases, we defined two reaction coordinates, one relative to the FMN relaxation and the other relative to amide relaxation. Subsequently, we computed normalized lifetimes (**Table 1**) and displayed *D* distributions against normalized lifetimes (**Fig. 5C**). In such representations, the relaxation events monitored by CNF31 and CNF35-A, which partially overlap due to the broad nature of the CNF31 distribution, clearly take place before any other processes (**Fig. 5C**). CNF136, CNF35-B, and CNF151 relax after the FMN adduct breaks (**Fig. 5C** top). The relaxation of CNF136 and CNF35-B are nearly simultaneous with the global protein relaxation (**Fig. 5C** bottom). Finally, the relaxation of CNF151 is slower and the distribution of rates broader than the protein backbone relaxation (**Fig. 5C** bottom). Therefore, from the point of view of the CNF probes, the results above point to at least three distinct kinetic classes of residue-specific relaxation times in EL222. First, a fast relaxation preceding both changes in the FMN and protein backbone and sensed as a red shift of CNF35-A and a red shift of CNF31. Second, a relaxation simultaneous with the protein backbone relaxation and sensed as a red shift of CNF136 and a blue shift of CNF35-B. Third, a slow relaxation posterior to the relaxation of the protein backbone and sensed as a red shift of CNF151.

**TABLE 1.** Normalized lit-to-dark recovery lifetimes. Normalized CNF-derived recovery times of single-CNF EL222 variants measured in H_2_O buffer and a possible explanation of the corresponding dynamical event based on mutagenesis and FTLS evidence. Absolute (non-normalized) lit-to-dark lifetimes appear in **Table S6**

| Probe | $\tau_{\text{CNF}}/\tau_{\text{FMN}}$ | $\tau_{\text{CNF}}/\tau_{\text{Amide}}$ | Tentative assignment |
| --- | --- | --- | --- |
| CNF31 | 0.5 | 0.2 | Disruption of A'α-LOV/ A'α-LOV interactions (oligomer dissociation) |
| CNF35-A | 0.6 | 0.2 | A'$\alpha$ packing against HTH (formation of caged conformation) |
| CNF35-B | 2.5 | 0.9 | Fold switching from disordered to α-helical (affecting primarily A'$\alpha$ and Jα elements) |
| CNF136 | 1.4 | 1.1 |  |
| CNF151 | 2.0 | 1.3 | Reconfiguration disordered loop |

We conclude that CNF labels located in different parts of the protein track several aspects of the relaxation to the dark-state that are not concurrent with changes in the FMN nor in the protein backbone. To directly compare the kinetics sensed by more than one CNF probe, without the need of using normalized lifetime distributions, we measured the dark recovery of EL222 variants carrying multiple CNF residues.

#### Multiple-CNF EL222 mutants

First, we prepared four double-CNF EL222 mutants: W31CNF/L35CNF, W31CNF/M151CNF, L35CNF/M151CNF, and Y136CNF/M151CNF (**Table S2** and **S3**). Absolute and difference infrared spectra in the CNF region are displayed in **Fig. 2A** bottom row. The lit-minus-dark steady-state difference spectra (SSDS) of three mutants were simple, with a single negative/positive signal pair. Only the pattern of the difference IR spectrum of the L35CNF/M151CNF mutant was more convoluted, presenting two positive bands at both edges together with a nearly zero differential absorbance in the central spectral region. This is consistent with our previous observation that the corresponding single-CNF mutants, L35CNF and M151CNF, showed frequency shifts in opposite directions: blue shifted and red shifted, respectively. Actually, the SSDS in the nitrile region of double-CNF mutants could be well reproduced as a linear combination of the two constituent single-CNF SSDS (**Fig. S22**), suggesting that the two introduced CNF groups worked as two independent probes. The single exception was W31CNF/L35CNF, probably as a result of the close spatial proximity of the two CNF moieties in the double mutant.

We next monitored the recovery kinetics after ceasing the illumination by time-resolved UV/Vis and IR spectroscopy (see full datasets in **Fig. S23-S26**). The *D* lifetime distributions of FMN and amide were more complicated than for single-CNF mutants, featuring shoulders, asymmetry and double peaks in some cases (**Fig. 6A**). Regarding the C≡N stretching mode, two dynamical events were clearly identified in three out of four cases (**Fig. 6A**). We attempted to assign the observed time-resolved spectra by assuming that each transient spectrum of the double-CNF mutants is the result of a linear combination of the two transient spectra of the corresponding single-CNF mutants (**Fig. 6B**), as explained in **Note S1**. In all three cases, the slowest dynamical event seen upon analysis of the time traces in the CNF absorption region is enriched in CNF151 relative to the fastest event (see molecular models and fractional contributions in **Fig. 6C**). Thus, the double-CNF mutants suggest that CNF151 returns to its dark state environment at a slower pace compared to the other three CNF probes (CNF31, CNF35 and CNF136), and that the relaxation times of CNF31 and CNF35 are similar. These observations are in agreement with the results obtained with single-CNF mutants.

**FIGURE 6.**
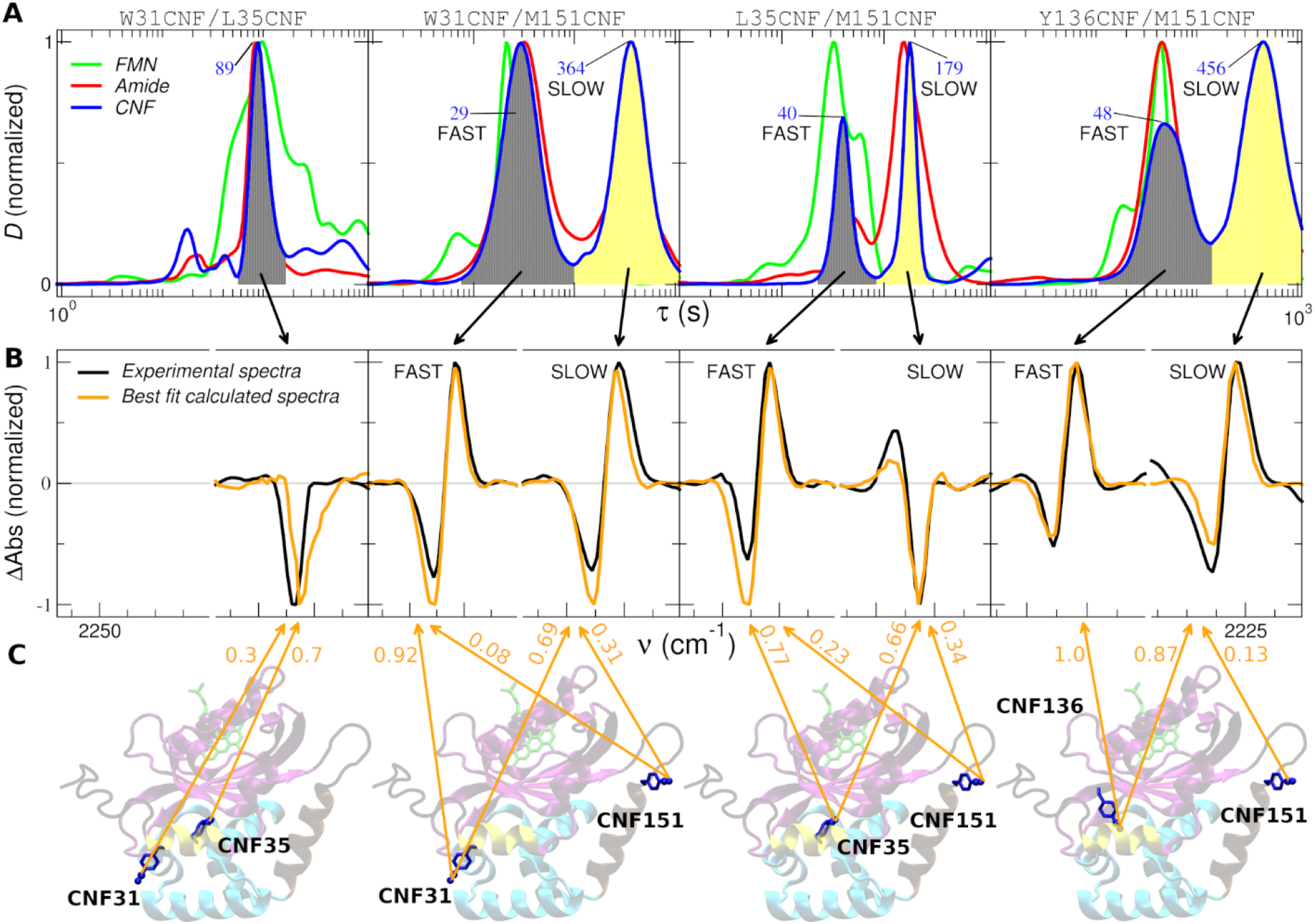
Disentangling residue-by-residue relaxation rates with double-CNF EL222 mutants. **(A)** Average dynamical content (*D*) as a function of lifetime (τ), derived from lifetime distribution analysis using the maximum entropy method for four double-CNF mutants (W31CNF/L35CNF, W31CNF/M151CNF, L35CNF/M151CNF, and Y136CNF/M151CNF). Curves were calculated for the relaxation of the FMN (green lines), amide bands (red lines), and the cyano moiety of the CNF residue (blue lines). The peak maxima, corresponding to the most probable lifetime (in seconds) are indicated in the case of CNF relaxation. **(B)** The transient decay-associated difference spectra of the kinetic events seen in the lit-to-dark transition of the cyano moiety were derived by integration over the shaded areas (black for the fastest event and yellow for the slowest event). Assuming that the transient spectra of double-CNF mutants are a linear combination of the corresponding single-CNF transient spectra, the relative contribution of each residue was retrieved by minimizing the discrepancy between experimental (black lines) and calculated (orange lines) spectra. The orange numbers on top of the structures (in C) report the fractional contribution of each CNF site to the best-fit calculated spectra. **(C)** Structural models of the double-CNF mutants in the dark state showing the location of the CNF residues and nitrile moiety (Van der Waals representation).

Is it feasible to follow the relaxation for more than two CNF groups simultaneously in a single EL222 protein? To test this, we prepared two triple-CNF mutants: W31CNF/L35CNF/M151CNF and L35CNF/Y136CNF/M151CNF (**Table S2** and **S3**). Steady-state absolute and difference infrared spectra in the CNF region are displayed in **Fig. 2A** bottom row. Regarding the lit-to-dark recovery kinetics, two events were resolved for W31CNF/L35CNF/M1151CNF while one event, with an asymmetric and broad peak in the lifetime distribution *D*, was resolved for L35CNF/Y136CNF/M151CNF (**Fig. S27-S29**). Hence, increasing the number of CNF reporters did not lead to a concomitant increase in the number of peaks in the *D* lifetime distribution. On the contrary, in the CNF relaxation in the triple-CNF mutant L35CNF/Y136CNF/M151CNF we resolved less kinetic events (one) than in the double-CNF mutant Y136CNF/M151CNF (two). To clarify to what extent this effect could be due to analytical limitations of the lifetime distribution analysis when several kinetic components overlap, we generated realistic synthetic datasets of multiple-CNF variants by averaging the individual experimental time-resolved matrices of the corresponding single-CNF variants (**Fig. S30**). Only two major dynamical events were retrieved by LDA using the maximum entropy method despite the fact that, individually, the data contain four different dynamical events. Even more revealing is the synthetic dataset corresponding to a hypothetical quadruple-CNF mutant (W31CNF/L35CNF/Y136CNF/M151CNF). In such a case, only a single dynamical event, corresponding to the fastest relaxation of CNF35, is found. Hence, it seems that there is a “dilution” effect in multiple-CNF mutants where the response of some CNF residues may be spectrally masked by other CNF sites depending on their relative intensities and their temporal overlap with other probes. Therefore, the experiments with mutants carrying three CNF probes do not rule out the presence of multiple (more than two) CNF-associated dynamical events, but rather point to limitations of the current lifetime distribution analysis when many kinetic components of different intensities spectrally and temporally overlap. Other limitations of our approach are detailed in **Note S4**.

Taken together, our integrative time-resolved UV/Vis/IR approach suggests that after removal of the light stimulus, the side-chain C≡N moieties, FMN chromophore and protein backbone do not return to the initial state simultaneously. Specifically, the recovery lifetimes follow the order: τ_CNF31_ ∼ τ_CNF35-A_ < τ_FMN_ < τ_Amide_ ∼ τ_CNF35-B_ ∼ τ_CNF136_ ≤ τ_CNF151_. Thus, CNF31, and partially CNF35 (CNF35-A), sense the relaxation path back to the dark state prior to all other probes, although the full recovery of CNF35 takes place later together with the protein backbone relaxation. The recovery of the FMN precedes that of the protein backbone. CNF136 and partially CNF35 (CNF35-B) relax simultaneously with backbone relaxation. Finally, the relaxation of CNF151 is delayed with respect to that of the protein backbone. Therefore, we have evidence that some CNF sites sense their recovery to the initial dark state environment at a rate significantly different from either the protein backbone structure or the FMN chromophore. With the help of FTLS and mutational analyses we tentatively assume that CNF31 reports on the A’α-LOV/A’α-LOV interface (oligomer/monomer EL222 equilibrium), and the fastest relaxation event of CNF35 reports on changes at the A’α-LOV/HTH interface (open/close equilibrium between the lit and dark state of EL222).

Critically, inclusion of CNF labels disclosed three additional dynamical events including the relaxation around residues CNF31, CNF35 and CNF151. CNF kinetics tend to resemble more amide kinetics than FMN kinetics. Indeed, CNF and amide probes rendered partially redundant kinetic information for several mutants. However, in the absence of information from amide relaxation, the interpretation of CNF-derived kinetics would have been challenging if not impossible to achieve. Lifetime distributions from CNF and FMN probes only overlapped each other in the case of two mutants. In short, none of the probes considered in isolation (side-chain C≡N, backbone C=O, and FMN) can deliver a complete picture of EL222 dark recovery mechanism and kinetics, thus making a case for the complementarity of the three.

#### Improved photocycle model of EL222

Based on the synergistic utilization of multiple probes (FMN, backbone, and CNF) and multiple sites (one, two, and three CNF residues), we propose the following model of the dark recovery reaction of the LOV-based transcription factor EL222 (**Fig. 7**). The dark state conformation can be safely assumed to represent the structure solved by X-ray crystallography (*2*). While the lit-state is presently unknown, it should be more oligomeric (dimer held together through interactions at least between the HTH domains), “open” (HTH domain decaged from LOV domain) and have less α-helical content (i.e. more random coil content) than the dark-state. Our data are in agreement with multiple parallel refolding pathways where different equilibria are present, each one with a distinct rate constant, but do not exclude the presence of sequential intermediates or more complex kinetic schemes, particularly concerning CNF35 relaxation. Indeed, not all kinetic events will be experienced by all proteins present in solution. For instance, the fraction of EL222 molecules that stay as monomers upon illumination may be spectrally silent with respect to CNF31 relaxation. Specifically, the lit-to-dark transition of EL222 can be described by up to seven equilibria, including one for the backbone relaxation, another one for the FMN and five for the residues (environments) CNF31, CNF35-A, CNF35-B, CNF136 and CNF151. These seven equilibria can be grouped into four categories based on their measured average recovery lifetimes in H_2_O. Ordered from fastest to slowest kinetics but without assuming any dependence among them, we have the following molecular events. In ∼25 s, EL222 collapses and the A’α-LOV fragment engages in interactions with the HTH domain (sensed by CNF35-A probe). The oligomer falls apart in similar time scales (CNF31 probe). In ∼40 s, the FMN-C78 covalent bond is broken (sensed by the UV/Vis absorption of the FMN probe). The protein backbone (most likely Jα, A’α, or both) increases its α-helical secondary structure content in ∼60 s (sensed by amide, CNF35-B and CNF136 probes). Equilibration of the environment around CNF151 takes place ∼80 s upon stopping illumination.

**FIGURE 7.**
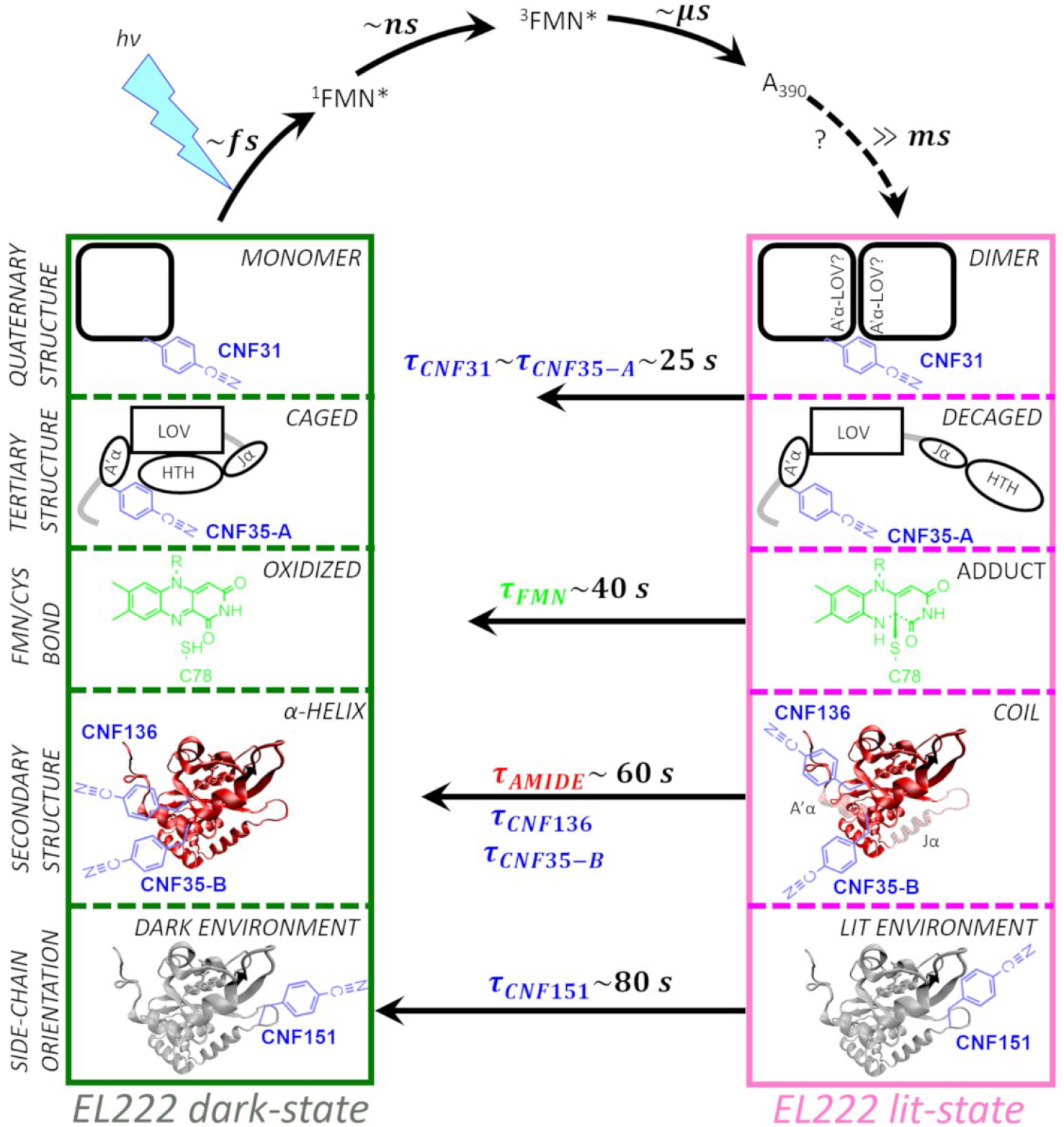
Kinetic-molecular model of EL222 photocycle. The model is defined in terms of five loosely coupled equilibria: monomer/oligomer, caged/decaged conformation, oxidized/C78-bound FMN, caged/decaged, helix/coil, and side-chain dark environment/lit environment. For the dark-to-lit transition, several intermediate species are formed (sequential model): the singlet (^1^FMN*), the triplet (^3^FMN*), the adduct (A_390_), and potentially others. For the lit-to-dark transition, each equilibrium exchanges at a specific rate (parallel model). The relative time scale of events appears to follow the order (from fastest to slowest): 1^st^) CNF31∼CNF35-A (the fastest kinetic event monitored by CNF35), 2^nd^) FMN, 3^rd^) Amide∼CNF136∼CNF35-B (the slowest kinetic event of CNF35), 4^th^) CNF151. Assuming that CNF31 informs on the A’α-LOV/A’α-LOV interface and that the “fast” CNF35 (CNF35-A) process reports on the A’α-LOV/HTH interface, a molecular interpretation of our results would be as follows. On average (population level), the fastest event is oligomer dissociation and formation of the “caged” conformation. The second step is the rupture of the FMN-C78 adduct. The third event is the increase in α-helical content of the protein backbone, which happens at a similar rate as the equilibration of CNF136 and some CNF35 side-chains. Finally, the slowest phase is the reorganization of the CNF151 side-chain. CNF, FMN, and amide probes are colored blue, green, and red, respectively.

In the dark-to-lit transition of EL222 measured in D_2_O, the time-resolved IR data is typically analyzed using a sequential kinetic model according to which a singlet species (^1^FMN*) is nearly instantaneously formed upon photon absorption, followed by the triplet species in ∼3 ns (^3^FMN*), and finally the adduct state in ∼5 μs (A_390_) (*5, 8*). Because the steady-state difference spectra are different from the transient spectra of the adduct state obtained a few microseconds after irradiation (**Fig. S31**), the existence of an additional intermediate species in the multi-millisecond time scale is envisioned, as previously postulated (*5*). Moreover, since some of the largest apparent CNF spectral changes are observed for those mutants that have higher tendency to self-associate than the wt (L123CNF, S140CNF, and Q141CNF), we hypothesize that such an additional intermediate may correspond to sparsely populated oligomeric species. This is in line with the low photo-dimerization yield of EL222 in the absence of its target DNA (*3, 4*).

## Discussion

### Non-canonical amino acids for site-specific infrared structural biology

We have studied the dark-state relaxation of EL222 from changes in the UV/Vis absorption spectra of the FMN chromophore and in the infrared absorption spectra resulting from protein backbone amide vibrations, complemented by spectral changes in the “transparent window” region of EL222 mutants containing the genetically encoded ncAA CNF. Major advantages of the co-translational incorporation of ncAA carrying vibrational reporters include: i) the freedom of choice regarding the labeling positions (we successfully introduced CNF at 45 different positions out of 45 attempts); ii) the high degree of incorporation efficiency (only two cases out of 45 showed clear levels of mis-incorporation); and iii) the possibility of simultaneous introduction of several reporters (so far we achieved triple CNF labeling in EL222). General benefits of using CNF are the sensitivity of the nitrile vibration to changes in the local electric field and solvent exposure, and its spectral simplicity. The CNF region typically contains one or at most two peaks, a situation that contrasts with the multitude of overlapping bands in the amide region, facilitating the isolation of dynamic changes as illustrated by cleaner lifetime distributions obtained for the former region. A drawback of the CNF region is its intrinsically local information content, which we compensated for by labelling many different residue locations and by simultaneously studying vibrational changes in the amide region (reporting on global secondary changes in structure), and changes in UV/Vis spectra (reporting on covalent bond formation between the FMN chromophore and C78). In the future, simultaneous use of CNF and other ncAA may provide access to even more dynamical events (**Note S4**).

Our results indicate that CNF residues inserted in selected positions of EL222 can, after a careful scrutiny, provide residue-specific information on the photocycle not attainable by other probes naturally present in EL222, like the FMN chromophore or the protein backbone. This observation is in line with previous studies on other photoreceptors that, in contrast to our work, focused on a relatively low number of unnatural residues (typically less than ten) or did not comprehensively address their potential perturbation (*23, 24, 28-30, 58*). Based on our results with a large collection of mutants (45 for a protein of 210 residues), a major caveat of genetically encoded non-canonical side-chains as site-specific infrared reporters is the risk of mis- or over-interpretations when only a limited number of such probes are tested. As a matter of fact, only two (Y136CNF, M151CNF) out of forty-five CNF residues faithfully recapitulated the kinetics seen in wt EL222. Others, like W31CNF, departed kinetically from wt behavior although they could still provide useful mechanistic information as reporters for changes in solvent accessibility (addressed by FTLS experiments). In total, only three CNF-derived dynamical events (CNF31, the fastest component of CNF35 and CNF151) provided genuinely unique kinetic information.

Additionally, looking at many CNF residues gives an overview of spectral and structural changes that can be mapped on the 1D (sequence level) and the 3D (conformational level) structure of EL222, the latter possible thanks to the availability of a high-resolution structure in its dark-state. Analysis of the infrared spectra in the “transparent window” region of CNF clearly shows long-distance signaling (allosteric communication) inside EL222. Perturbation of the FMN chromophore and binding pocket (the allosteric site) by photon absorption is transferred to the HTH module (the active site where DNA binding occurs) and other regions like A’α and Jα elements. This observation is in agreement with previous results based on NMR spectroscopy, although the residue-by-residue sensitivity of the two techniques differs (*2*).

Importantly, the spectral observables (e.g. peak positions and areas) may be transformed into physical variables, like local electric fields and H-bonding, thus providing new avenues for the synergistic use of experimental and physics-based computational approaches. In summary, ncAA-assisted optical spectroscopy complements the information content provided by common structural biology techniques and helps delineating signal transduction pathways in EL222.

### Dark reversion kinetics as witnessed by the chromophore, protein and residues

Previous studies on LOV photorecovery have largely concentrated on the UV/Vis region (*9*) rather than on the IR region (*43*). Here we measured the thermal recovery of EL222 by both UV/Vis and FTIR spectroscopies. In principle, the relaxation times of different structural elements in proteins may be different, which may be inferred by independently analyzing different IR frequencies (*54, 59*). This was not the case for EL222 because the Arrhenius plots were linear and essentially identical at all three probed frequency ranges (1750-1680 cm^-1^, 1680-1600 cm^-1^ and 1600-1500 cm^-1^). In turn, these results suggest that the dark recovery of EL222 conforms to a two-state folding transition (*54*). Previously, non-Arrhenius behavior (two different slopes) was observed for the recovery kinetics of EL222 based on FMN absorbance (*60*). Here we restricted ourselves to temperatures at or below 35 °C to avoid partial unfolding of the protein (melting temperature is 55 °C). Indeed, if we compare only the data below 35 °C, the two studies agree on the same activation energy (∼15 kcal/mol) for the rupture of the flavin-cysteinyl bond. The ∼18 kcal/mol barrier seen in the IR region agrees with its slower kinetics and indicates a larger energy expenditure involved in rearranging the structure of EL222 in the transition state relative to adduct breakage. The observed KIE (more than 4-fold slower relaxation kinetics in D_2_O than H_2_O) is in agreement with previous reports of similar LOV domains based on UV/Vis data (*9, 61*). The high KIE can be explained because the adduct breakage requires releasing a proton and an electron, and the folding of α-helices necessitates the breaking of protein-water H-bonds and formation of new intramolecular H-bonds.

For wt EL222 at 20 °C, *D* values suggest a decoupling between FMN and protein relaxation where, on average, the FMN-C78 bond is broken earlier (∼41 s in H_2_O or ∼178 s in D_2_O) and then the protein recovers its secondary structure content (∼57 s in H_2_O or ∼220 s in D_2_O). These results contrast with the synchronous relaxation of protein and FMN structures seen in other LOV domains by circular dichroism (CD), NMR and UV/Vis spectroscopies (*10, 61, 62*). Such discrepancies are rather puzzling since CD, NMR and IR techniques are all sensitive to protein secondary structure changes. One could argue that the spectral overlap between the protein backbone and the FMN absorption bands in the far-UV region could contaminate the CD kinetics, biasing it toward the UV/Vis-determined rates, which is solely due to FMN. Concerning the previous NMR study, we notice that the method is sensitive to backbone dynamics whose relaxation kinetics could be simpler than the side-chain relaxation kinetics probed by the CNF. Nevertheless, asynchronous dark relaxation of chromophore, backbone and native side-chains was previously reported for Slr1694, a related photoreceptor belonging to the BLUF (blue-light using flavin adenine dinucleotide) family, based on a hybrid UV/Vis/IR approach (*63*). Also in agreement with our observations, a delayed recovery of the protein structure with respect to the flavin adduct decay has been recently described for a dimeric two-domain LOV photoreceptor based on small-angle X-ray scattering and UV/Vis (*64*).

### Implications for protein folding dynamics and signal transduction

The lit-to-dark transition of EL222 resembles the denatured-to-native refolding of unfolded proteins, where secondary structure acquisition ensues chain collapse (*65*). High cooperativity would imply that once the rate-limiting step has occurred, all subsequent steps would follow at essentially the same time. Such a situation is only observed in the case of N53CNF and L216CNF variants. For the rest of the CNF mutants, our kinetic results indicate a more complex picture where (i) CNF dark recovery rates are residue-specific and (ii) CNF recovery rates need not match neither the local FMN relaxation nor the global protein relaxation rates. These results add to the growing body of reports on protein folding dynamics showing different relaxation times for different residues (*66, 67*), even between the main-chain and side-chain of the same residue (*68*), or between global and local probes (*69, 70*). Because our methodological approach is restricted to the ensemble level, the proposed relative order of events in our four-step kinetic model (**Fig. 7**) is the most probable one. In other words, at the single-molecule level, another order of events would be a priori possible, as shown by molecular dynamics simulations of protein folding (*71*).

If the intrinsic time scales of thio-adduct breakage (picoseconds) (72), chain collapse (microseconds) (*73*), protein monomerization (milliseconds) (*74*), helix-coil transitions (nanoseconds) (*75*), and side-chain rotation (picoseconds) (*76*), are all much faster than the experimentally determined dark recovery times of the whole population of EL222, what is then the reason for observed lifetimes (multi-seconds) and what is the nature of the rate-limiting step? Light-induced protein structural dynamics and allostery typically follows a defined set of sequential conformational transitions (*41, 77, 78*). The light-induced dark-to-lit transition of EL222 is not an exception: FMN photochemistry drives, and therefore precedes, all subsequent conformational changes through a cascade of intermediates. Indeed, secondary structure changes (α-helix unfolding) are observed concomitantly with adduct formation (*5, 8*). In contrast, the dark-induced lit-to-dark transition of EL222 is not necessarily caused by adduct breakage since other sites (CNF31, CNF35) experience a change in their surroundings at earlier times. The escape from heterogeneous traps rather than the nucleation rate has been proposed to account for the residue-specific folding rates of α-helices (*66*). In other LOV sensors, dynamic interactions at protein/protein interfaces are considered additional rate-limiting factors for the conformational recovery (*64*). Our results highlight the A’α element, where both CNF31 and CNF35 reside, as a central hub in controlling the dark recovery rate of EL222. Furthermore, we hypothesize that the dissociation of the A’α-LOV/A’α-LOV interface i.e. dimer disassembly, and/or the formation of A’α-LOV/HTH interface might be the rate-determining step(s) in the lit-to-dark transition of EL222. Our kinetic model is also compatible with weakly coupled equilibria between the chromophore, local protein structure and global protein conformation, as recently described for another photoreceptor (*79*). This would imply that photosensory receptors have redundant signaling pathways that may be important to ensure robust responses to light or absence of it (*79*). Overall, our work stresses again the importance of integrative approaches using multiple probes, residues and techniques to comprehensively characterize protein folding landscapes (*59, 80*).

## Materials and Methods

### Materials

CNF was purchased from Bide Pharmatech. The plasmid encoding EL222 was a kind gift of Dr. Kevin Gardner. The plasmid encoding the intein-chitin binding domain-12his fusion protein was a kind gift of Dr. Edward Lemke. The final plasmid for genetic code expansion encoded an N-terminal EL222 protein (amino acids 17 to 225) followed by a C-terminal intein-CBD-12his purification tag. The plasmid pDule2-pCNF encoding the orthogonal translation system (OTS) composed of a polyspecific aminoacyl tRNA synthetase (*Mj*CNFRS) and suppressor tRNA (*Mj*tRNA_CUA_) was a kind gift from Dr. Ryan Mehl. All other materials can be found in the **Key Resource Table**.

### Protein expression

TAG sequences were introduced by site-directed mutagenesis. *Escherichia coli* BL21(DE3) (or C321.ΔA.exp(DE3) in case of the triple mutants) were co-transformed with a plasmid encoding EL222 (with one, two or three TAG triplets at desired positions) and another plasmid encoding an OTS able to incorporate CNF in response to UAG codons. Bacteria were grown in terrific broth (TB) media (or exceptionally glucose-based minimal media in the case of W89CNF and I94CNF mutants) at 37 °C up to an optical density at 600 nm of 0.4 at which point CNF (freshly dissolved in 3M NaOH) was added to the medium at a final concentration of 1 mM. Cells were grown for thirty more minutes, then the temperature was lowered to 20 °C and gene expression was induced with 0.5 mM IPTG in the dark.

### Protein purification and characterization

All protein purification steps were conducted under very dim light and 4 °C unless otherwise stated. After 16 hours, cells were harvested by centrifugation at 6000 g and disrupted by sonication for 4 minutes on ice (50% of duty cycle, 50 mW). Cell suspensions were clarified by centrifugation at 60000 g for 30 minutes and the supernatant was purified by immobilized metal affinity chromatography (IMAC) with a Ni^2+^ resin. Loosely bound proteins were removed with 40 mM imidazole washes and fusion proteins were eluted with 500 mM imidazole. Addition of 100 mM DTT activated the self-splicing intein moiety that cleaved off itself from EL222 (48 hours at 23 °C). Cleavage mixtures were purified by a second IMAC step to remove non-cleaved proteins. The flow-through was polished by size-exclusion chromatography (SEC) in a Superdex75Increase column with 50 mM MES, 100 mM NaCl, pH=6.8 as running buffer. The oligomeric states of all the purified proteins were estimated by interpolating the measured elution volumes on a calibration curve constructed using proteins of known molecular weight. In cases where the oligomeric state could not be unambiguously identified the mutants are labeled as multimers. Fractions were analyzed by SDS-PAGE, pooled and concentrated (**Table S3**) using 10 kDa molecular-weight cut-off filters. Proteins were quantified by two methods: UV/visible spectroscopy (FMN concentration) assuming an extinction coefficient at 450 nm of 13000 M^-1^cm^-1^, and IR spectroscopy (protein concentration). The latter is proportional to the concentration of C=O bonds (mostly arising from the protein backbone, with minor contributions from the two FMN carbonyls and side-chain carbonyls). The yields of CNF mutants were variable and in general lower by at least one order of magnitude (**Table S3**).

All proteins were analyzed by mass spectrometry to determine the molecular weight and check for the presence of the CNF residue (**Table S3**). Proteins were diluted with 100 μL of 5% acetic acid in water and loaded onto Opti-trap C4 cartridge (Optimize Technologies), washed 4 x with 250 μL of 5% acetic acid in water and eluted with 100 μL of 80% acetonitrile, 5% acetic acid. Proteins were analyzed by direct infusion using syringe pump at a flow rate 2 μL/min connected with an electrospray ion source of 15T solariX XR FT-ICR mass spectrometer (Bruker Daltonics). The mass spectrometer was externally calibrated using 1% (w/w) sodium trifluoracetate. Proteins were measured in positive mode with 2M data acquisition. The data were processed using SNAP algorithm, a part of DataAnalysis 4.4 software (Bruker Daltonics).

### Steady-state FTIR

The stationary infrared spectra of all the EL222 variants were measured with a Bruker Vertex 70v FTIR spectrometer equipped with a globar source, KBr beamsplitter and a liquid nitrogen cooled mercury cadmium telluride (MCT) detector. The protein samples were centrifuged at 18000 g for 15 minutes to remove any precipitants before measurements. D_2_O buffer exchange was done by centrifugal concentration method using 10 kDa cutoff filters. The aperture value was set in accordance to get maximum signal without saturating the detector. For measurements in the “transparent window” region or in D_2_O buffer, ∼25 μl of sample was loaded into a temperature controllable demountable liquid cell with CaF2 windows and a 50 μm Teflon spacer. For measuring amide region in H_2_O buffer, ∼10 μl of sample and 6 μm Teflon spacer was used instead. For each sample, three spectra (dark state, lit state and difference) were recorded at 20 °C and 200-300 scans were averaged in the spectral range of 4000 to 900 cm^-1^ with a resolution of 2 cm^-1^. Protein lit state was induced by illuminating blue light of 450 nm LED with power of 25 mW/cm^2^ on the sample throughout the measurement. Dark and lit samples were measured against buffer and the difference spectra were recorded as the lit spectra against the dark spectra. Protein samples had concentration of 1-2 mM in storage buffer (**Table S3**).

The resulting spectra where processed with SpectraGryph software. Baseline correction and Gaussian curve fitting was done with OriginPro (OriginLab).

Same setup of FTIR spectrometer mentioned above was used for frequency-temperature line slope (FTLS) measurements. Spectra were recorded at 5, 15, 25, and 35 °C averaging over 300 scans at a resolution of 1 cm^-1^. First, the reference spectrum for each temperature was measured followed by the protein samples. The samples were equilibrated for 10 minutes before measurement.

### Time-resolved rapid-scan FTIR experiments

Transient infrared spectra with a time resolution of tens of milliseconds were recorded on a Bruker Vertex 70v FTIR spectrometer. For the nitrile region, we used a 4.50 μm bandpass filter. Same sample amount, concentration, path length and temperature was used as in steady state FTIR. For the amide region, no filter was used and a 6 μm path-length was used to minimize the contribution of water absorption. The spectral resolution was 4 cm^-1^. At a single time point, 10 scans were averaged and for the cyano region it was further averaged 3 times. Samples were first illuminated for three minutes under continuous irradiation to accumulate the lit state. Then the LED was switched off and recovery kinetics were recorded over 10 minutes in the case of H_2_O-based experiments or 15 min for D_2_O-based experiments.

### Steady-state UV/Vis spectroscopy

UV-visible absorbance spectra in the range of 320 to 550 nm of EL222 variants were measured with a Specord 50 Plus spectrometer (Analytik Jena) using a quartz cuvette (Hellma Analytics) having 10 mm path length at 20 °C and 70 μL sample volume. Protein samples were diluted to get approximately 0.5 optical density at 450 nm.

### Time-resolved UV/Vis spectroscopy

Dark state recovery kinetics of protein variants were measured by first continuously illuminating the sample for 60 seconds with blue light of 450 nm LED with power of approximately 25 mW/cm^2^ (M450L1, Thorlabs). LED was then turned off and spectra were subsequently taken for 10 min in the case of H_2_O-based experiments or 15 min for D_2_O-based experiments. The samples were the same as for the FTIR experiments but they were diluted ∼70 times to a concentration between 10 and 30 μM, equivalent to an absorbance at 450 nm ∼0.5.

Additional methods can be found in Supplementary **Notes S1, S2**, and **S3**.

## Author contributions

Conceptualization: VALF, GF

Formal analysis, methodology: ASC, VALF, GF

Funding acquisition: BS

Investigation: ASC, AC, CAOD, GF

Resources: PCA, YL

Software: VALF Visualization: ASC, GF

Writing—original draft: GF

Writing—review & editing: ASC, IA, VALF, GF

## Acknowledgments

The work was supported by the project Structural dynamics of biomolecular systems (ELIBIO) (CZ.02.1.01/0.0/0.0/15_003/0000447) from the European Regional Development Fund (ERDF) and the Ministry of Education, Youth and Sports (MEYS) of the Czech Republic. The Institute of Biotechnology of the Czech Academy of Sciences acknowledges the institutional grant RVO86652036. We acknowledge CF Biophysic, CF SMS of CIISB, Instruct-CZ Centre BIOCEV, supported by MEYS CR (LM2018127)) and ERDF-Project "UP CIISB” (No. CZ.02.1.01/0.0/0.0/18_046/0015974). We thank the financial support provided by the Ministerio de Ciencia e Innovación (MICINN) - Agencia Estatal de Investigación (AEI) through projects BFU2016-768050-P, BFU2017-91559-EXP, PID2019-106103GBI00 and CTQ2017-87372-P, and by the Generalitat Valenciana through the project PROMETEU/2019/066.

## Data and materials availability

Steady-state and time-resolved datasets will be deposited in Zenodo (<u>10.5281/zenodo.7086623</u>). Other data, software and materials are available from the corresponding authors upon reasonable request. Plasmids can be shared upon signing of a Material Transfer Agreement (MTA).

