## Supplementary information for "Genetically encoded non-canonical amino acids reveal asynchronous dark reversion of chromophore, backbone and side-chains in EL222"

**Figure S19. Time-resolved UV/VIS/IR spectroscopy of EL222 L216CNF**.....33
**Figure S20. Analysis of time-resolved spectroscopy data of other single-CNF variants**.....35
**Figure S21. Lifetime distribution analysis of other single-CNF variants.**.....36
**Figure S22. Predictions of steady-state difference spectra (SSDS) of double-CNF variants.**.....37
**Figure S23. Time-resolved UV/VIS/IR spectroscopy of EL222 W31CNF/L35CNF**.....38
**Figure S24. Time-resolved UV/VIS/IR spectroscopy of EL222 W31CNF/M151CNF**.....38
**Figure S25. Time-resolved UV/VIS/IR spectroscopy of EL222 L35CNF/M151CNF**.....39
**Figure S26. Time-resolved UV/VIS/IR spectroscopy of EL222 Y136CNF/M151CNF**.....39
**Figure S27. Time-resolved UV/VIS/IR spectroscopy of EL222 W31CNF/L35CNF/M151CNF**.....40
**Figure S28. Time-resolved UV/VIS/IR spectroscopy of EL222 L35CNF/Y136CNF/M151CNF**.....40
**Figure S29. Disentangling residue-specific relaxation rates with triple-CNF EL222 mutants**.....41
**Figure S30. Simulations of lifetime density maps and averaged dynamical contents (*D*)**.....42
**Figure S31. Comparison of transient and stationary IR spectra of wt EL222 in $D_2O$**.....43
**Supplementary references**.....44

### NOTE S1: Analysis of spectroscopic data.

#### Steady-state spectroscopy.

Baseline correction was done by applying a second order polynomial using the method of least absolute value residuals (FTIR datasets) or a line (UV/Visible datasets). If necessary, an offset was applied by setting to zero the absorbance at 440-450 nm (UV/Vis datasets) or 1760-1800 cm$^{-1}$ (FTIR datasets in the amide region). No offset was applied in the CNF region. Offset correction is particularly important for time-resolved datasets in order to avoid baseline fluctuations leaking into the kinetics.

In case of CNF spectra, we noticed that the "direct" lit-minus-dark IR difference spectra, where the dark state is used as the reference spectrum for FTIR recording, used to be less noisy than the "manual" difference spectrum, where the dark-state spectrum is subtracted from the lit-state spectrum both being recorded against buffer. We therefore reconstructed the lit-state spectra by summing the difference spectra to the dark-state spectra. The peak maxima ($\nu_{max}$), areas and electric fields ($|\vec{F}|$) in the cyano stretching region (2250 to 2215 cm$^{-1}$) were calculated for both dark and lit states using custom scripts. Subsequently, light-induced changes were calculated as:

$$\Delta\nu_{max} = \nu_{max}^{lit} - \nu_{max}^{dark} \qquad \text{EQ. 1}$$

$$\Delta|\vec{F}| = |\vec{F}|^{lit} - |\vec{F}|^{dark} \qquad \text{EQ. 2}$$

The electric fields were calculated from the peak areas as described in (*45*). Briefly, transition dipole moment ($\vec{m}$) were obtained from the concentration-normalized peak areas by:

$$|\vec{m}| = \sqrt{\frac{3\,\epsilon_0 hc\,ln10}{2\,\pi^2\,N_A}\int\frac{\varepsilon(\bar{\nu})d\bar{\nu}}{\bar{\nu}}} \qquad \text{EQ. 3}$$

where $\varepsilon(\bar{\nu})$ is the extinction coefficient, $\varepsilon(\bar{\nu})d\bar{\nu}$ is the integrated peak area, and $\epsilon_0$, $h$, $c$, $N_A$, and $\bar{\nu}$ are the vacuum permittivity, Planck's constant, the speed of light, Avogadro's number, and the wavenumber frequency, respectively. Transition dipole moments were then converted into electric fields by using the $|\vec{m}|$-$|\vec{F}|$ calibration curves for ortho-nitrotoluene (*45*):

$$|\vec{m}| = -0.00092|\vec{F}| + 0.0388 \qquad \text{EQ. 4}$$

Frequency shifts due to H-bonding ($\Delta\nu_{HB}$) where calculated as the difference between the measured frequencies and theoretical frequencies expected according to the aprotic solvent field-frequency correlation curve of ortho-nitrotoluene in polarizable force-fields (*45*):

$$\Delta\nu_{HB} = \nu_{max} - (0.19|\vec{F}| + 2231.4) \qquad \text{EQ. 5}$$

For FTLS experiments, baseline-corrected absolute spectra were fitted to one Gaussian-shaped band in OriginPro (OriginLab). The L35CNF mutant necessitated two Gaussian bands to achieve reasonable fits but the fits themselves were not consistent at all temperatures. In this case, multiple Gaussian bands were fitted to the steady-state difference spectra of L35CNF and the peaks labeled in Figure S7 were tracked. The difference in the Gaussian-fitted frequency ($\nu_G$) between a given temperature and 5 °C ($\nu_G^{T=5°C}$) was calculated:

$$\Delta\nu_G = \nu_G^T - \nu_G^{T=5°C} \qquad \text{EQ. 6}$$

Multiple-CNF spectra were assumed to be described as a linear combination of single-CNF spectra. For a double-CNF mutant containing CNF residues X and Y, the expected transient difference spectra (DADS) were calculated as follows:

$$DADS_{XY}^{EXPECTED} = f * DADS_X^{OBSERVED} + (1-f) * DADS_Y^{OBSERVED} \qquad \text{EQ. 7}$$

where f is the fractional contribution of each single-CNF spectra. Best-fit reconstructed spectra (Figure 6B) were selected as the one whose f would give with the lowest deviation ($X^2$) with respect to the observed (experimental) DADS:

$$X^2 = \sum_{vmin}^{vmax}\frac{\left(DADS_{XY}^{OBSERVED} - DADS_{XY}^{EXPECTED}\right)^2}{DADS_{XY}^{EXPECTED}} \qquad \text{EQ. 8}$$

#### Time-resolved spectroscopy.

Data pre-processing prior to kinetic analyses comprised up to three steps: logarithmic averaging, removal of water vapor contributions, and baseline correction.

Logarithmic averaging with 20 points per decade was conducted for all time-resolved data. This step not only reduces the size of the dataset, speeding up calculations, but makes the data more evenly distributed in logarithmic scale, which is more appropriate than linearly distributed data when kinetic analysis is conducted over several orders of time.

Digital subtraction of water vapor contribution was only necessary for the time-resolved data in the 1800-1500 $cm^{-1}$ region. Digital subtraction was done using a water vapor spectrum recorded by letting air from the room into the sample compartment, with a time-dependent subtraction factor automatically determined to cancel the area of a water vapor band between 1765 to 1780 $cm^{-1}$.

Time-resolved UV/vis data were baseline corrected by making an offset in the 540-550 nm region, free from absorption changes. For time-resolved FTIR data in the CNF region, baseline correction was not considered necessary. In the amide region, there were important baseline fluctuations in the 1700-1600 $cm^{-1}$ region because of the strong background absorption from liquid $H_2O$. An offset in the absorption-free 1760-1800 $cm^{-1}$ region partially reduce these baseline fluctuations.

Lifetime distribution analysis (LDA) (*81*) was applied to pre-processed time-resolved datasets (A panels of Fig. S8-9, S14-S19, and S23-S28). Corrected differential absorbance (light-minus-dark) as a function of frequency $\omega_i$ (or wavelength in the case of UV/Vis data) and time delay $t_j$ (after switching lights off) were then fitted to a quasi-distributed sum of exponentials plus a constant term:

$$\Delta A(\omega_i, t_j) = a_0(\omega_i) + \sum_{k=1}^{n} a(\omega_i, \tau_k) e^{-t_j/\tau_k} \quad \text{EQ. 9}$$

where the index $k$ refers to a kinetic component with time constant $\tau_k$, and $a$ (or $a_o$) are the pre-exponential factors (amplitudes). Time-resolved datasets were numerically transformed from the time domain to the lifetime domain by finding the lifetime distribution that maximizes an entropy function as well as minimizes the squared sum of the residuals between the data and the fit (*52*). To this end, we used a quasi-continuous distribution of $\tau_k$ (200 lifetimes from $10^{-1}$ seconds to $10^{4}$ seconds spread on a logarithmic scale). The main advantage of the maximum entropy method with respect to the Tikhonov regularization, another common method for the analysis of time-resolved spectra (*82*), is in effectively suppressing artificial oscillations in the estimated lifetime distribution that can be confused with the actual kinetic components (*83*). As a second advantage, the maximum entropy method typically provides better-resolved lifetime distributions than Tikhonov regularization does for the same level of noise in the data (84). When the standard deviation of the noise in the data depends on time and/or on spectral frequency, such dependence needs to be accounted in the analysis by appropriately weighting the residuals between the data and the fit. We assumed that the noise standard deviation dependence on time and spectral frequency are independent of each other, and estimated these dependences iteratively from the residual matrix between the data and a fit. The balance between the two opposite goals (maximizing entropy and minimizing differences between the data and fit) is controlled by a scalar, the so-called regularization factor ($\lambda$). As $\lambda$ gets smaller the obtained lifetime distributions are more detailed and fit better the data, but eventually they start to include unreliable features. An optimal regularization factor ($\lambda_{opt}$), needed to balance the detail of the lifetime distribution with its sensitivity to noise in the data, is key for an optimal estimation of the lifetime distribution. $\lambda_{opt}$ was selected based on the L-curve (*85*), which is a plot of the residual norm (goodness of fit) as a function of the smoothing norm (solution smoothness), as the one corresponding to its corner (Table S9). The resulting 2D lifetime distributions, matrices of $a(\omega_i, \tau_k)$, are shown in the B panels of Fig. S8-9, S14-S19, and S23-S28).

Finally, the average dynamical content *D* was calculated from the 2D lifetime density maps as the integral (summation) over the squared amplitudes $a(\omega_i, \tau_k)$ for a given frequency (or wavelength) interval:

$$D = \sqrt{\sum_i a(\omega_i, \tau_k)^2} \qquad \text{EQ. 10}$$

*D* values as a function of lifetime are shown in Fig. 5, Fig. 6A, Fig. S10 and Fig. S21. Lifetimes with the largest associated *D* values (peak maxima) are listed in Table S6.

The transient spectra, equivalent to a decay-associated difference spectra (DADS) (*81*), shown in Fig. S20 and S21 were obtained by integration (summation) of the differential absorbance over time constant intervals corresponding to the peaks of the *D* lifetime distribution:

$$DADS(\omega_i) = \sum_{t_{j,min}=\tau_{k,min}}^{t_{j,max}=\tau_{k,max}} \Delta A(\omega_i) \qquad \text{EQ. 11}$$

All time-resolved experiments were done in triplicate. In some replicates, the recovery kinetics featured additional events. However, their small amplitude, unrealistic associated spectra, and lack of reproducibility, make their interpretation difficult. For the sake of simplicity and to avoid over-interpretation, peaks in the *D* lifetime distribution with a relative area smaller than 15 % were not further discussed.

#### NOTE S2: Molecular modeling.

EL222 models were constructed starting from the dark-state structure (Protein Data Bank ID: 3P7N) in two steps. First, residues missing in the crystal structure were added. Second, native residues were replaced by the ncAA CNF.

Ten residues in the N-terminal domain (Ala17 to Pro28) and seven residues in the loop connecting the LOV and HTH domains (Asp144 to Met151) were added using Modeller software (*86*). 1000 models were created for the missing N-terminal residues and other 1000 models were created for the missing loops. Results were then evaluated according to scoring functions. For chain A, both root mean square deviation (RMSD) and Estimated Overlap (3.5 Å) functions predict a model with a Chi-squared value of 1.35. This model was used for subsequent steps.

Structural data for 4-cyano-L-phenylalanine (entry 4CF) was obtained from SwissSidechain (*87*), a database of non-natural side-chains (**https://www.swisssidechain.ch/**). 4CF was inserted into EL222 at 45 different positions using the provided plugin for the Chimera suite. Only the most probable rotamer was kept (Table S5). Molecular models of dark-state EL222 colored according to the experimental light-induced shifts (Fig. 2C, Fig. S5B, and Fig. S13B) were done in Chimera (version 1.14) using the "Render by Attribute" structure analysis tool. Molecular models shown in Fig. 1A, Fig. 4B, Fig. 6C, and Fig. 7 were created in Visual Molecular Dynamics (*88*) and rendered using POV-ray. Note that all residue numbers are consistent with the UniprotKB entry **Q2NB98**, which differs from the corresponding PDB entry **3P7N** by a factor of +3.

#### NOTE S3: Light-dependent equilibria and photoconversion yields.

The light-dependent equilibria between “dark” and “lit” structural states of EL222 can be expressed as:

$$EL222_{DARK} \rightleftharpoons EL222_{LIT} \quad \text{EQ. 12}$$

where light shifts the equilibria towards the right and darkness favors the left-hand side. The dark state can be assumed to correspond to the crystal structure and is characterized by a monomeric “closed” conformation (LOV domain caged by the HTH domain) with FMN being held in place by non-covalent interactions. The lit state(s) is presently unknown at atomic resolution but accumulated evidence indicates that such a state is more oligomeric, “open” (LOV domain detached from HTH domain), and features the FMN chromophore covalently bound to Cys78 (adduct state).

The above equilibria can be decomposed into three elemental equilibria depending on the probing group: FMN (320-450 nm), protein backbone amide bands (1750-1500 cm$^{-1}$) or the nitrile side-chain of CNF (2270-2200 cm$^{-1}$):

$FMN_{450} \rightleftharpoons FMN_{390}$, more commonly referred to as $D_{450} \rightleftharpoons A_{390}$ EQ. 13

$$Amide_{HELIX} \rightleftharpoons Amide_{COIL} \quad \text{EQ. 14}$$

$$CNF^{i}_{DARK-ENVIRONMENT} \rightleftharpoons CNF^{i}_{LIT-ENVIRONMENT} \quad \text{EQ. 15}$$

While equilibria 13 and 14 are unique, there are as many EQ. 15 as residues *i* in the protein. Unfolding affects only a few residues, unknown so far but most likely belonging to A’α and/or Jα structural elements. Hence, the extent of coupling between EQ. 13-15 depends on the particular residue being investigated. For instance, a residue that does not sense adduct formation and that does not undergo a secondary structure change upon illumination, may still experience a change in its surroundings, due to oligomerization or tertiary structure changes:

$$CNF^{i}_{MONOMER} \rightleftharpoons CNF^{i}_{OLIGOMER} \quad \text{EQ. 16}$$

$$CNF^{i}_{CAGED} \rightleftharpoons CNF^{i}_{DECAGED} \quad \text{EQ. 17}$$

Assuming a two-state transition for the embedded FMN chromophore, the photoconversion yield between EL222 can be estimated as the fraction of EL222 molecules with semireduced FMN covalently bound to Cys78 upon illumination. This corresponds to the $A_{390}$ species (equivalent to 1 – $D_{450}$, the latter being the fraction of EL222 molecules with oxidized FMN non-covalently bound):

$$f_{A_{390}} = 1 - f_{D_{450}} = \frac{1-A^{SAMPLE}_{450}}{1-A^{L}_{450}} \ (UV/Vis) \quad \text{EQ. 18}$$

where $A^{SAMPLE}_{450}$ is the absorbance at 450 nm of a given variant under continuous illumination. Since light scattering from the LED source prevents UV/Vis spectra acquisition and simultaneous irradiation, $A^{SAMPLE}_{450}$ was estimated as the extrapolated absorbance at 450 nm at time zero using the fitted pre-exponential factors (Note S1). $A^{L}_{450}$ is the absorbance at 450 nm of the lit state (0.06), that was estimated from the spectra of EL222 AQTRIP (a quadruple-mutation variant that has a UV/Vis lifetime of ~60 min in $D_2O$ and thus the spectra taken a few seconds after illumination contain >99% of EL222 molecules in the lit state. Photoconversion yields can be found in Table S8**.**

Unfortunately, a similar procedure cannot be applied in the IR spectral region in order to calculate the photoconversion yield of protein backbone and CNF side-chains. This is because positive and negative bands partially cancel out in FTIR difference spectra, making the area smaller the higher the overlap between such bands. Moreover, in the amide I (I’) region all residues contribute (besides water if not correctly subtracted). Finally, in the CNF region, distinct extinction coefficients in the dark and lit states may confound this type of analysis.

### NOTE S4: Limitations, ideas and speculations.

Four main limitations of our hybrid protein engineering/spectroscopy approach are outlined. The first is that being non-native residues, CNF probes will be in some cases sufficiently intrusive to impede revealing a common mechanism (*34*). Their *intrusiveness* can sometimes be easily identified. For instance, a few of our CNF mutant solutions are transparent, in contrast with the yellow color of wt EL222, thus indicating the absence of FMN cofactor. In other cases, CNF induces a subtle perturbation. This is the case of N53CNF and L216CNF mutants that resemble the wt behavior in everything except for the amide band, which relaxes synchronously with the FMN and CNF, indicating that the kinetic mechanism under study has been affected. Therefore, we urge researchers using this approach not to extract conclusions uncritically without first doing all reasonable control experiments. Generally speaking, CNF kinetics are meaningless if not interpreted in the context of FMN and amide kinetics. Overall, we started with 45 CNF mutants and only two (Y136CNF, M151CNF) met the criteria of being non-kinetically perturbing and sufficient signal-to-noise ratio for demanding time-resolved experiments. If the introduced mutations do actually alter the kinetics, it may still be possible to draw some conclusions, provided that sufficient data is gathered. This is the case of our mutants W31CNF and L35CNF, whose absolute CNF kinetics are difficult to extrapolate to the wt in the absence of information about amide- and FMN-derived kinetics.

The second limitation is the *low level of throughput*: residues must be investigated one-by-one, i.e., 1 protein -> 1 ncAA. This is in contrast to the 1 protein -> multiple cAA approach in the NMR field where multidimensional experiments of uniformly labeled proteins can give access to several residues at once (*89*). In principle, two-dimensional (and higher order) IR spectroscopies could permit the assignment of proteins labeled simultaneously with several IR probes, but these methods are still limited to small peptides (*90*). This limitation is not only on the technical (spectroscopy) side but also on the protein synthesis side. For EL222, three simultaneous CNF labels is the maximum that we have attempted and looking at the protein yields and fidelities, we would be surprised if we could tag more than 4 positions. Nevertheless, the GCE method has been shown to permit the introduction of up to 30 copies of the same ncAA (*91*). A promising alternative would be to incorporate two frequency-resolved probes (18). So far, up to 4 distinct ncAA have been genetically encoded in a single protein (*92*). Moreover, UV/Vis and infrared spectroscopies typically require two separate set-ups. However, recent studies suggest the feasibility of simultaneous spectral acquisition in the UV/Vis and IR spectral ranges (*93*). While cyano and amide bands are better recorded using different pathlenghts, other probes with larger extinction coefficients, e.g. azidophenylalanine, may be monitored concomitantly with global protein bands (*28*).

The third limitation is the *interpretation* of the observed CNF shifts. In principle, both large-scale protein conformational changes and short-range side-chain "tuning" motions could give rise to the same frequency shifts and area changes. We notice that peak areas were converted to electric fields using a calibration curve developed for ortho-tolunitrile in a chosen single force field (*45*). Therefore, the applicability of this model to (para-)CNF in EL222 and other proteins remains to be determined. Two cases deserve special attention: CNF31 and CNF35. Through the joint analysis of FTLS, mutational and kinetics data, we speculate that CNF31 reports on oligomerization (interactions between A'α-LOV regions) and CNF35 (the fastest component) reports on tertiary structure contacts (interactions between LOV and HTH domains). These two hypotheses could be directly tested by small-angle scattering, FRET or electron paramagnetic resonance (*94-96*).

The fourth limitation lies in the *assignment* of IR spectral changes when more than one CNF group is incorporated. The steady-state spectra of double- and triple-CNF mutants contain two or three overlapping bands, respectively. While static spectral overlap may be partially alleviated in kinetic studies, as seen in the L35CNF/M151CNF and W31CNF/M151CNF mutants, we are cautious about our interpretation. We believe that the use of two frequency-resolved probes will facilitate the unambiguous and simultaneous tracking of the two side-chains.

### Table S0. Key resource table.

| REAGENT/RESOURCE | SOURCE | CATALOG NUMBER/IDENTIFIER |
|---|---|---|
| **Strains, plasmids and chemicals** | | |
| BL21 (DE3) *E. coli* strain | NA | NA |
| C321.ΔA.exp *E. coli* strain | Addgene | 49018 |
| C321.ΔA.exp(DE3) *E. coli* strain | This paper | NA |
| pParallel1-EL222-Intein-CBD-12His plasmid (x58) | This paper | NA |
| pDule2-pCNF plasmid | Addgene (http://n2t.net/addgene:85495) | 85495 |
| λDE3 Lysogenization Kit | Novagen | 69734 |
| p-cyanophenylalanine (CNF) | BLD Pharmatech | BD18790 |
| Isopropyl β- d-1-thiogalactopyranoside (IPTG) | Formedium | IPTG100 |
| Carbenicillin | Formedium | CAR0025 |
| Spectinomycin | Duchefa | S0188 |
| Dithiothreitol (DTT) | Formedium | DTT010 |
| Ni-NTA Resin | Thermo Scientific | 88222 |
| **Software** | | |
| Matlab R2021a | https://www.mathworks.com | NA |
| SpectraGryph 1.2 | https://www.effemm2.de/ | NA |
| Glotaran 1.5.1 | https://glotaran.org/ | NA |
| OriginPro 2020 | https://www.originlab.com | NA |
| MaxEnt_iLT2D | This paper | NA |
| QtGrace v0.2.6 | https://sourceforge.net/projects/qtgrace/ | NA |
| GIMP 2.10.8 | https://www.gimp.org/ | NA |
| VMD 1.9.3 | https://www.ks.uiuc.edu/Research/vmd/ | NA |
| Chimera 1.14 | https://www.cgl.ucsf.edu/chimera/ | NA |
| **Instruments and accessories** | | |
| FTIR spectrometer | Bruker | Vertex v70 |
| Sample cell | Specac | GS20512 |
| $CaF_2$ windows | Specac | GS20522 |
| Mounted LED | Thorlabs | M455L4 |
| LED Driver | Thorlabs | LEDD1B |
| Collimator lens | Thorlabs | LB1092 |
| Collimator lens holder | Thorlabs | SM1L10, SM1A6T |
| IR bandpass filter | Thorlabs | FB4500-500 |
| Temperature controller for IR | Huber | Ministat 125 |
| Direct Detect IR spectrometer | Millipore | DDHW00010-WW |
| UV-Vis spectrometer | Analytik Jena | Specord 50 Plus |
| UV-Vis spectrometer (nanodrop) | DeNovix | DS-11 FX + |
| Gel filtration chromatography system | Biorad | NGC system |
| Gel filtration column | Cytiva | Superdex 75 Increase 10/300 GL |
| Centrifuges | Beckman Coulter | Allegra X-30R, Avanti JXN-30 |
| Centrifugal concentrator | Sartorius | VS0201, VS0102, VS15RH02 |
| Sonicator | Qsonica sonicators | Q700 Sonicator |
| Microfluidizer/Homogenizer | Avestin | Emulsiflex C3 |
| Mass spectrometer | Bruker Daltonics | solariX XR |

#### TABLE S1. Target residues.

| Mutant | Role |
|---|---|
| *A18CNF* | Disordered N-terminal loop |
| *W31CNF* | Similarity to CNF, putative light-induced A’α unfolding |
| *L35CNF* | LOV-HTH interface, NMR shifts, putative light-induced A’aα unfolding |
| *I36CNF* | NMR shifts, putative light-induced A’α unfolding |
| *A42CNF* | Signal transduction |
| *V44CNF* | Altered photocycle in AQtrip |
| *S46CNF* | FMN binding pocket |
| *N53CNF* | FMN binding pocket |
| *L55CNF* | Altered photocycle in AQtrip |
| *F62CNF* | Similarity to CNF |
| *Y68CNF* | Similarity to CNF |
| *C78CNF* | Adduct formation with FMN |
| *F80CNF* | Similarity to CNF |
| *A82CNF* | FMN binding pocket |
| *W89CNF* | Similarity to CNF |
| *T91CNF* | FMN binding pocket |
| *I94CNF* | FMN binding pocket |
| *R95CNF* | FMN binding pocket |
| *I108CNF* | FMN binding pocket |
| *N110CNF* | FMN binding pocket |
| *Y111CNF* | Similarity to CNF |
| *F118CNF* | Similarity to CNF |
| *N120CNF* | FMN binding pocket |
| *V122CNF* | Role in dimerization in other LOV domains |
| *L123CNF* | LOV-HTH interface, |
| *V124CNF* | FMN binding pocket, altered photocycle in AQTrip |
| *Y128CNF* | LOV-HTH interface, similarity to CNF |
| *Y136CNF* | Similarity to CNF |
| *F137CNF* | FMN binding pocket |
| *S140CNF* | LOV-HTH interface, role in dimerization |
| *Q141CNF* | FMN binding pocket, signal transduction |
| *Q148CNF* | Disordered linker |
| *P149CNF* | Disordered linker |
| *M151CNF* | Putative light-induced Jα unfolding |
| *M153CNF* | Putative light-induced Jα unfolding |
| *M162CNF* | Putative light-induced Jα unfolding |
| *L166CNF* | Putative light-induced Jα unfolding |
| *L181CNF* | NMR shifts |
| *G191CNF* | α2-α3 loop |
| *M199CNF* | DNA-binding helix |
| *M205CNF* | DNA-binding helix |
| *L210CNF* | α3-α4 loop |
| *T212CNF* | LOV-HTH interface |
| *L216CNF* | NMR shifts |
| *I225CNF* | Disordered C-terminus |

### TABLE S2. Protein/RNA sequences.

| Biopolymer | | Sequence |
|---|---|---|
| **EL222** | *WT* | GADDTRVEVQPPAQWVLDLIEASPIASVVSDPRLADNPLIAINQAFTDLTGYSEEECVGRNCRFLAGSGTEPWLTDKIR<br>QGVREHKPVLVEILNYKKDGTPFRNAVLVAPIYDDDDELLYFLGSQVEVDDDQPNMGMARRERAAEMLKTLSPRQLEVT<br>TLVASGLRNKEVAARLGLSEKTVKMHRGLVMEKLNLKTSADLVRIAVEAGIA |
| **Single-CNF** | *A18CNF* | G*DDTRVEVQPPAQWVLDLIEASPIASVVSDPRLADNPLIAINQAFTDLTGYSEEECVGRNCRFLAGSGTEPWLTDKIR<br>QGVREHKPVLVEILNYKKDGTPFRNAVLVAPIYDDDDELLYFLGSQVEVDDDQPNMGMARRERAAEMLKTLSPRQLEVT<br>TLVASGLRNKEVAARLGLSEKTVKMHRGLVMEKLNLKTSADLVRIAVEAGIA |
| | *W31CNF* | GADDTRVEVQPPAQ*VLDLIEASPIASVVSDPRLADNPLIAINQAFTDLTGYSEEECVGRNCRFLAGSGTEPWLTDKIR<br>QGVREHKPVLVEILNYKKDGTPFRNAVLVAPIYDDDDELLYFLGSQVEVDDDQPNMGMARRERAAEMLKTLSPRQLEVT<br>TLVASGLRNKEVAARLGLSEKTVKMHRGLVMEKLNLKTSADLVRIAVEAGIA |
| | *L35CNF* | GADDTRVEVQPPAQWVLD*IEASPIASVVSDPRLADNPLIAINQAFTDLTGYSEEECVGRNCRFLAGSGTEPWLTDKIR<br>QGVREHKPVLVEILNYKKDGTPFRNAVLVAPIYDDDDELLYFLGSQVEVDDDQPNMGMARRERAAEMLKTLSPRQLEVT<br>TLVASGLRNKEVAARLGLSEKTVKMHRGLVMEKLNLKTSADLVRIAVEAGIA |
| | *I36CNF* | GADDTRVEVQPPAQWVLDL*EASPIASVVSDPRLADNPLIAINQAFTDLTGYSEEECVGRNCRFLAGSGTEPWLTDKIR<br>QGVREHKPVLVEILNYKKDGTPFRNAVLVAPIYDDDDELLYFLGSQVEVDDDQPNMGMARRERAAEMLKTLSPRQLEVT<br>TLVASGLRNKEVAARLGLSEKTVKMHRGLVMEKLNLKTSADLVRIAVEAGI |
| | *A42CNF* | GADDTRVEVQPPAQWVLDLIEASPI*SVVSDPRLADNPLIAINQAFTDLTGYSEEECVGRNCRFLAGSGTEPWLTDKIR<br>QGVREHKPVLVEILNYKKDGTPFRNAVLVAPIYDDDDELLYFLGSQVEVDDDQPNMGMARRERAAEMLKTLSPRQLEVT<br>TLVASGLRNKEVAARLGLSEKTVKMHRGLVMEKLNLKTSADLVRIAVEAGIA |
| | *V44CNF* | GADDTRVEVQPPAQWVLDLIEASPIAS*VSDPRLADNPLIAINQAFTDLTGYSEEECVGRNCRFLAGSGTEPWLTDKIR<br>QGVREHKPVLVEILNYKKDGTPFRNAVLVAPIYDDDDELLYFLGSQVEVDDDQPNMGMARRERAAEMLKTLSPRQLEVT<br>TLVASGLRNKEVAARLGLSEKTVKMHRGLVMEKLNLKTSADLVRIAVEAGIA |
| | *S46CNF* | GADDTRVEVQPPAQWVLDLIEASPIASVV*DPRLADNPLIAINQAFTDLTGYSEEECVGRNCRFLAGSGTEPWLTDKIR<br>QGVREHKPVLVEILNYKKDGTPFRNAVLVAPIYDDDDELLYFLGSQVEVDDDQPNMGMARRERAAEMLKTLSPRQLEVT<br>TLVASGLRNKEVAARLGLSEKTVKMHRGLVMEKLNLKTSADLVRIAVEAGIA |
| | *N53CNF* | GADDTRVEVQPPAQWVLDLIEASPIASVVSDPRLAD*PLIAINQAFTDLTGYSEEECVGRNCRFLAGSGTEPWLTDKIR<br>QGVREHKPVLVEILNYKKDGTPFRNAVLVAPIYDDDDELLYFLGSQVEVDDDQPNMGMARRERAAEMLKTLSPRQLEVT<br>TLVASGLRNKEVAARLGLSEKTVKMHRGLVMEKLNLKTSADLVRIAVEAGIA |
| | *L55CNF* | GADDTRVEVQPPAQWVLDLIEASPIASVVSDPRLADNP*IAINQAFTDLTGYSEEECVGRNCRFLAGSGTEPWLTDKIR<br>QGVREHKPVLVEILNYKKDGTPFRNAVLVAPIYDDDDELLYFLGSQVEVDDDQPNMGMARRERAAEMLKTLSPRQLEVT<br>TLVASGLRNKEVAARLGLSEKTVKMHRGLVMEKLNLKTSADLVRIAVEAGIA |
| | *F62CNF* | GADDTRVEVQPPAQWVLDLIEASPIASVVSDPRLADNPLIAINQA*TDLTGYSEEECVGRNCRFLAGSGTEPWLTDKIR<br>QGVREHKPVLVEILNYKKDGTPFRNAVLVAPIYDDDDELLYFLGSQVEVDDDQPNMGMARRERAAEMLKTLSPRQLEVT<br>TLVASGLRNKEVAARLGLSEKTVKMHRGLVMEKLNLKTSADLVRIAVEAGIA |
| | *Y68CNF* | GADDTRVEVQPPAQWVLDLIEASPIASVVSDPRLADNPLIAINQAFTDLTG*SEEECVGRNCRFLAGSGTEPWLTDKIR<br>QGVREHKPVLVEILNYKKDGTPFRNAVLVAPIYDDDDELLYFLGSQVEVDDDQPNMGMARRERAAEMLKTLSPRQLEVT<br>TLVASGLRNKEVAARLGLSEKTVKMHRGLVMEKLNLKTSADLVRIAVEAGIA |
| | *C78CNF* | GADDTRVEVQPPAQWVLDLIEASPIASVVSDPRLADNPLIAINQAFTDLTGYSEEECVGRN*RFLAGSGTEPWLTDKIR<br>QGVREHKPVLVEILNYKKDGTPFRNAVLVAPIYDDDDELLYFLGSQVEVDDDQPNMGMARRERAAEMLKTLSPRQLEVT<br>TLVASGLRNKEVAARLGLSEKTVKMHRGLVMEKLNLKTSADLVRIAVEAGIA |
| | *F80CNF* | GADDTRVEVQPPAQWVLDLIEASPIASVVSDPRLADNPLIAINQAFTDLTGYSEEECVGRNCR*LAGSGTEPWLTDKIR<br>QGVREHKPVLVEILNYKKDGTPFRNAVLVAPIYDDDDELLYFLGSQVEVDDDQPNMGMARRERAAEMLKTLSPRQLEVT<br>TLVASGLRNKEVAARLGLSEKTVKMHRGLVMEKLNLKTSADLVRIAVEAGIA |
| | *A82CNF* | GADDTRVEVQPPAQWVLDLIEASPIASVVSDPRLADNPLIAINQAFTDLTGYSEEECVGRNCRFL*GSGTEPWLTDKIR<br>QGVREHKPVLVEILNYKKDGTPFRNAVLVAPIYDDDDELLYFLGSQVEVDDDQPNMGMARRERAAEMLKTLSPRQLEVT<br>TLVASGLRNKEVAARLGLSEKTVKMHRGLVMEKLNLKTSADLVRIAVEAGIA |
| | *W89CNF* | GADDTRVEVQPPAQWVLDLIEASPIASVVSDPRLADNPLIAINQAFTDLTGYSEEECVGRNCRFLAGSGTEP*LTDKIR<br>QGVREHKPVLVEILNYKKDGTPFRNAVLVAPIYDDDDELLYFLGSQVEVDDDQPNMGMARRERAAEMLKTLSPRQLEVT<br>TLVASGLRNKEVAARLGLSEKTVKMHRGLVMEKLNLKTSADLVRIAVEAGIA |
| | *T91CNF* | GADDTRVEVQPPAQWVLDLIEASPIASVVSDPRLADNPLIAINQAFTDLTGYSEEECVGRNCRFLAGSGTEPWL*DKIR<br>QGVREHKPVLVEILNYKKDGTPFRNAVLVAPIYDDDDELLYFLGSQVEVDDDQPNMGMARRERAAEMLKTLSPRQLEVT<br>TLVASGLRNKEVAARLGLSEKTVKMHRGLVMEKLNLKTSADLVRIAVEAGIA |
| | *I94CNF* | GADDTRVEVQPPAQWVLDLIEASPIASVVSDPRLADNPLIAINQAFTDLTGYSEEECVGRNCRFLAGSGTEPWLTDK*R<br>QGVREHKPVLVEILNYKKDGTPFRNAVLVAPIYDDDDELLYFLGSQVEVDDDQPNMGMARRERAAEMLKTLSPRQLEVT<br>TLVASGLRNKEVAARLGLSEKTVKMHRGLVMEKLNLKTSADLVRIAVEAGIA |
| | *R95CNF* | GADDTRVEVQPPAQWVLDLIEASPIASVVSDPRLADNPLIAINQAFTDLTGYSEEECVGRNCRFLAGSGTEPWLTDKI*<br>QGVREHKPVLVEILNYKKDGTPFRNAVLVAPIYDDDDELLYFLGSQVEVDDDQPNMGMARRERAAEMLKTLSPRQLEVT<br>TLVASGLRNKEVAARLGLSEKTVKMHRGLVMEKLNLKTSADLVRIAVEAGIA |
| | *I108CNF* | GADDTRVEVQPPAQWVLDLIEASPIASVVSDPRLADNPLIAINQAFTDLTGYSEEECVGRNCRFLAGSGTEPWLTDKIR<br>QGVREHKPVLVE*LNYKKDGTPFRNAVLVAPIYDDDDELLYFLGSQVEVDDDQPNMGMARRERAAEMLKTLSPRQLEVT<br>TLVASGLRNKEVAARLGLSEKTVKMHRGLVMEKLNLKTSADLVRIAVEAGIA |
| | *N110CNF* | GADDTRVEVQPPAQWVLDLIEASPIASVVSDPRLADNPLIAINQAFTDLTGYSEEECVGRNCRFLAGSGTEPWLTDKIR<br>QGVREHKPVLVEIL*YKKDGTPFRNAVLVAPIYDDDDELLYFLGSQVEVDDDQPNMGMARRERAAEMLKTLSPRQLEVT<br>TLVASGLRNKEVAARLGLSEKTVKMHRGLVMEKLNLKTSADLVRIAVEAGIA |
| | *Y111CNF* | GADDTRVEVQPPAQWVLDLIEASPIASVVSDPRLADNPLIAINQAFTDLTGYSEEECVGRNCRFLAGSGTEPWLTDKIR<br>QGVREHKPVLVEILN*KKDGTPFRNAVLVAPIYDDDDELLYFLGSQVEVDDDQPNMGMARRERAAEMLKTLSPRQLEVT<br>TLVASGLRNKEVAARLGLSEKTVKMHRGLVMEKLNLKTSADLVRIAVEAGIA |
| | *F118CNF* | GADDTRVEVQPPAQWVLDLIEASPIASVVSDPRLADNPLIAINQAFTDLTGYSEEECVGRNCRFLAGSGTEPWLTDKIR<br>QGVREHKPVLVEILNYKKDGTP*RNAVLVAPIYDDDDELLYFLGSQVEVDDDQPNMGMARRERAAEMLKTLSPRQLEVT<br>TLVASGLRNKEVAARLGLSEKTVKMHRGLVMEKLNLKTSADLVRIAVEAGIA |
| | *N120CNF* | GADDTRVEVQPPAQWVLDLIEASPIASVVSDPRLADNPLIAINQAFTDLTGYSEEECVGRNCRFLAGSGTEPWLTDKIR<br>QGVREHKPVLVEILNYKKDGTPFR*AVLVAPIYDDDDELLYFLGSQVEVDDDQPNMGMARRERAAEMLKTLSPRQLEVT<br>TLVASGLRNKEVAARLGLSEKTVKMHRGLVMEKLNLKTSADLVRIAVEAGIA |
| | *V122CNF* | GADDTRVEVQPPAQWVLDLIEASPIASVVSDPRLADNPLIAINQAFTDLTGYSEEECVGRNCRFLAGSGTEPWLTDKIR<br>QGVREHKPVLVEILNYKKDGTPFRNA*LVAPIYDDDDELLYFLGSQVEVDDDQPNMGMARRERAAEMLKTLSPRQLEVT<br>TLVASGLRNKEVAARLGLSEKTVKMHRGLVMEKLNLKTSADLVRIAVEAGIA |

| | | |
|---|---|---|
| | *L123CNF* | GADDTRVEVQPPAQWVLDLIEASPIASVVSDPRLADNPLIAINQAFTDLTGYSEEECVGRNCRFLAGSGTEPWLTDKIR<br>QGVREHKPVLVEILNYKKDGTPFRNAV*VAPIYDDDDELLYFLGSQVEVDDDQPNMGMARRERAAEMLKTLSPRQLEVT<br>TLVASGLRNKEVAARLGLSEKTVKMHRGLVMEKLNLKTSADLVRIAVEAGIA |
| | *V124CNF* | GADDTRVEVQPPAQWVLDLIEASPIASVVSDPRLADNPLIAINQAFTDLTGYSEEECVGRNCRFLAGSGTEPWLTDKIR<br>QGVREHKPVLVEILNYKKDGTPFRNAVL*APIYDDDDELLYFLGSQVEVDDDQPNMGMARRERAAEMLKTLSPRQLEVT<br>TLVASGLRNKEVAARLGLSEKTVKMHRGLVMEKLNLKTSADLVRIAVEAGIA |
| | *Y128CNF* | GADDTRVEVQPPAQWVLDLIEASPIASVVSDPRLADNPLIAINQAFTDLTGYSEEECVGRNCRFLAGSGTEPWLTDKIR<br>QGVREHKPVLVEILNYKKDGTPFRNAVLVAPI*DDDDELLYFLGSQVEVDDDQPNMGMARRERAAEMLKTLSPRQLEVT<br>TLVASGLRNKEVAARLGLSEKTVKMHRGLVMEKLNLKTSADLVRIAVEAGIA |
| | *Y136CNF* | GADDTRVEVQPPAQWVLDLIEASPIASVVSDPRLADNPLIAINQAFTDLTGYSEEECVGRNCRFLAGSGTEPWLTDKIR<br>QGVREHKPVLVEILNYKKDGTPFRNAVLVAPIYDDDDELL*FLGSQVEVDDDQPNMGMARRERAAEMLKTLSPRQLEVT<br>TLVASGLRNKEVAARLGLSEKTVKMHRGLVMEKLNLKTSADLVRIAVEAGIA |
| | *F137CNF* | GADDTRVEVQPPAQWVLDLIEASPIASVVSDPRLADNPLIAINQAFTDLTGYSEEECVGRNCRFLAGSGTEPWLTDKIR<br>QGVREHKPVLVEILNYKKDGTPFRNAVLVAPIYDDDDELLY*LGSQVEVDDDQPNMGMARRERAAEMLKTLSPRQLEVT<br>TLVASGLRNKEVAARLGLSEKTVKMHRGLVMEKLNLKTSADLVRIAVEAGIA |
| | *S140CNF* | GADDTRVEVQPPAQWVLDLIEASPIASVVSDPRLADNPLIAINQAFTDLTGYSEEECVGRNCRFLAGSGTEPWLTDKIR<br>QGVREHKPVLVEILNYKKDGTPFRNAVLVAPIYDDDDELLYFLG*QVEVDDDQPNMGMARRERAAEMLKTLSPRQLEVT<br>TLVASGLRNKEVAARLGLSEKTVKMHRGLVMEKLNLKTSADLVRIAVEAGIA |
| | *Q141CNF* | GADDTRVEVQPPAQWVLDLIEASPIASVVSDPRLADNPLIAINQAFTDLTGYSEEECVGRNCRFLAGSGTEPWLTDKIR<br>QGVREHKPVLVEILNYKKDGTPFRNAVLVAPIYDDDDELLYFLGS*VEVDDDQPNMGMARRERAAEMLKTLSPRQLEVT<br>TLVASGLRNKEVAARLGLSEKTVKMHRGLVMEKLNLKTSADLVRIAVEAGIA |
| | *Q148CNF* | GADDTRVEVQPPAQWVLDLIEASPIASVVSDPRLADNPLIAINQAFTDLTGYSEEECVGRNCRFLAGSGTEPWLTDKIR<br>QGVREHKPVLVEILNYKKDGTPFRNAVLVAPIYDDDDELLYFLGSQVEVDDD*PNMGMARRERAAEMLKTLSPRQLEVT<br>TLVASGLRNKEVAARLGLSEKTVKMHRGLVMEKLNLKTSADLVRIAVEAGIA |
| | *P149CNF* | GADDTRVEVQPPAQWVLDLIEASPIASVVSDPRLADNPLIAINQAFTDLTGYSEEECVGRNCRFLAGSGTEPWLTDKIR<br>QGVREHKPVLVEILNYKKDGTPFRNAVLVAPIYDDDDELLYFLGSQVEVDDDQ*NMGMARRERAAEMLKTLSPRQLEVT<br>TLVASGLRNKEVAARLGLSEKTVKMHRGLVMEKLNLKTSADLVRIAVEAGIA |
| | *M151CNF* | GADDTRVEVQPPAQWVLDLIEASPIASVVSDPRLADNPLIAINQAFTDLTGYSEEECVGRNCRFLAGSGTEPWLTDKIR<br>QGVREHKPVLVEILNYKKDGTPFRNAVLVAPIYDDDDELLYFLGSQVEVDDDQPN*GMARRERAAEMLKTLSPRQLEVT<br>TLVASGLRNKEVAARLGLSEKTVKMHRGLVMEKLNLKTSADLVRIAVEAGIA |
| | *M153CNF* | GADDTRVEVQPPAQWVLDLIEASPIASVVSDPRLADNPLIAINQAFTDLTGYSEEECVGRNCRFLAGSGTEPWLTDKIR<br>QGVREHKPVLVEILNYKKDGTPFRNAVLVAPIYDDDDELLYFLGSQVEVDDDQPNMG*ARRERAAEMLKTLSPRQLEVT<br>TLVASGLRNKEVAARLGLSEKTVKMHRGLVMEKLNLKTSADLVRIAVEAGIA |
| | *M162CNF* | GADDTRVEVQPPAQWVLDLIEASPIASVVSDPRLADNPLIAINQAFTDLTGYSEEECVGRNCRFLAGSGTEPWLTDKIR<br>QGVREHKPVLVEILNYKKDGTPFRNAVLVAPIYDDDDELLYFLGSQVEVDDDQPNMGMARRERAAE*LKTLSPRQLEVT<br>TLVASGLRNKEVAARLGLSEKTVKMHRGLVMEKLNLKTSADLVRIAVEAGIA |
| | *L166CNF* | GADDTRVEVQPPAQWVLDLIEASPIASVVSDPRLADNPLIAINQAFTDLTGYSEEECVGRNCRFLAGSGTEPWLTDKIR<br>QGVREHKPVLVEILNYKKDGTPFRNAVLVAPIYDDDDELLYFLGSQVEVDDDQPNMGMARRERAAEMLKT*SPRQLEVT<br>TLVASGLRNKEVAARLGLSEKTVKMHRGLVMEKLNLKTSADLVRIAVEAGIA |
| | *L181CNF* | GADDTRVEVQPPAQWVLDLIEASPIASVVSDPRLADNPLIAINQAFTDLTGYSEEECVGRNCRFLAGSGTEPWLTDKIR<br>QGVREHKPVLVEILNYKKDGTPFRNAVLVAPIYDDDDELLYFLGSQVEVDDDQPNMGMARRERAAEMLKTLSPRQLEVT<br>TLVASG*RNKEVAARLGLSEKTVKMHRGLVMEKLNLKTSADLVRIAVEAGIA |
| | *G191CNF* | GADDTRVEVQPPAQWVLDLIEASPIASVVSDPRLADNPLIAINQAFTDLTGYSEEECVGRNCRFLAGSGTEPWLTDKIR<br>QGVREHKPVLVEILNYKKDGTPFRNAVLVAPIYDDDDELLYFLGSQVEVDDDQPNMGMARRERAAEMLKTLSPRQLEVT<br>TLVASGLRNKEVAARL*LSEKTVKMHRGLVMEKLNLKTSADLVRIAVEAGIA |
| | *M199CNF* | GADDTRVEVQPPAQWVLDLIEASPIASVVSDPRLADNPLIAINQAFTDLTGYSEEECVGRNCRFLAGSGTEPWLTDKIR<br>QGVREHKPVLVEILNYKKDGTPFRNAVLVAPIYDDDDELLYFLGSQVEVDDDQPNMGMARRERAAEMLKTLSPRQLEVT<br>TLVASGLRNKEVAARLGLSEKTVK*HRGLVMEKLNLKTSADLVRIAVEAGIA |
| | *M205CNF* | GADDTRVEVQPPAQWVLDLIEASPIASVVSDPRLADNPLIAINQAFTDLTGYSEEECVGRNCRFLAGSGTEPWLTDKIR<br>QGVREHKPVLVEILNYKKDGTPFRNAVLVAPIYDDDDELLYFLGSQVEVDDDQPNMGMARRERAAEMLKTLSPRQLEVT<br>TLVASGLRNKEVAARLGLSEKTVKMHRGLV*EKLNLKTSADLVRIAVEAGIA |
| | *L210CNF* | GADDTRVEVQPPAQWVLDLIEASPIASVVSDPRLADNPLIAINQAFTDLTGYSEEECVGRNCRFLAGSGTEPWLTDKIR<br>QGVREHKPVLVEILNYKKDGTPFRNAVLVAPIYDDDDELLYFLGSQVEVDDDQPNMGMARRERAAEMLKTLSPRQLEVT<br>TLVASGLRNKEVAARLGLSEKTVKMHRGLVMEKLN*KTSADLVRIAVEAGIA |
| | *T212CNF* | GADDTRVEVQPPAQWVLDLIEASPIASVVSDPRLADNPLIAINQAFTDLTGYSEEECVGRNCRFLAGSGTEPWLTDKIR<br>QGVREHKPVLVEILNYKKDGTPFRNAVLVAPIYDDDDELLYFLGSQVEVDDDQPNMGMARRERAAEMLKTLSPRQLEVT<br>TLVASGLRNKEVAARLGLSEKTVKMHRGLVMEKLNLK*SADLVRIAVEAGIA |
| | *L216CNF* | GADDTRVEVQPPAQWVLDLIEASPIASVVSDPRLADNPLIAINQAFTDLTGYSEEECVGRNCRFLAGSGTEPWLTDKIR<br>QGVREHKPVLVEILNYKKDGTPFRNAVLVAPIYDDDDELLYFLGSQVEVDDDQPNMGMARRERAAEMLKTLSPRQLEVT<br>TLVASGLRNKEVAARLGLSEKTVKMHRGLVMEKLNLKTSAD*VRIAVEAGIA |
| | *I225CNF* | GADDTRVEVQPPAQWVLDLIEASPIASVVSDPRLADNPLIAINQAFTDLTGYSEEECVGRNCRFLAGSGTEPWLTDKIR<br>QGVREHKPVLVEILNYKKDGTPFRNAVLVAPIYDDDDELLYFLGSQVEVDDDQPNMGMARRERAAEMLKTLSPRQLEVT<br>TLVASGLRNKEVAARLGLSEKTVKMHRGLVMEKLNLKTSADLVRIAVEAG*A |
| **Double-CNF** | *W31CNF/L35CNF* | GADDTRVEVQPPAQ*VLDLIEASPIASVVSDPRLADNPLIAINQAFTDLTGYSEEECVGRNCRFLAGSGTEPWLTDKIR<br>QGVREHKPVLVEILNYKKDGTPFRNAVLVAPIYDDDDELLYFLGSQVEVDDDQPN*GMARRERAAEMLKTLSPRQLEVT<br>TLVASGLRNKEVAARLGLSEKTVKMHRGLVMEKLNLKTSADLVRIAVEAGIA |
| | *W31CNF/M151CNF* | GADDTRVEVQPPAQ*VLDLIEASPIASVVSDPRLADNPLIAINQAFTDLTGYSEEECVGRNCRFLAGSGTEPWLTDKIR<br>QGVREHKPVLVEILNYKKDGTPFRNAVLVAPIYDDDDELLYFLGSQVEVDDDQPN*GMARRERAAEMLKTLSPRQLEVT<br>TLVASGLRNKEVAARLGLSEKTVKMHRGLVMEKLNLKTSADLVRIAVEAGIA |
| | *L35CNF/M151CNF* | GADDTRVEVQPPAQWVLD*IEASPIASVVSDPRLADNPLIAINQAFTDLTGYSEEECVGRNCRFLAGSGTEPWLTDKIR<br>QGVREHKPVLVEILNYKKDGTPFRNAVLVAPIYDDDDELLYFLGSQVEVDDDQPN*GMARRERAAEMLKTLSPRQLEVT<br>TLVASGLRNKEVAARLGLSEKTVKMHRGLVMEKLNLKTSADLVRIAVEAGIA |
| | *Y136CNF/M151CNF* | GADDTRVEVQPPAQWVLDLIEASPIASVVSDPRLADNPLIAINQAFTDLTGYSEEECVGRNCRFLAGSGTEPWLTDKIR<br>QGVREHKPVLVEILNYKKDGTPFRNAVLVAPIYDDDDELL*FLGSQVEVDDDQPN*GMARRERAAEMLKTLSPRQLEVT<br>TLVASGLRNKEVAARLGLSEKTVKMHRGLVMEKLNLKTSADLVRIAVEAGIA |
| **Triple-CNF** | *W31CNF/L35CNF/ M151CNF* | MGADDTRVEVQPPAQ*VLD*IEASPIASVVSDPRLADNPLIAINQAFTDLTGYSEEECVGRNCRFLAGSGTEPWLTDKI<br>RQGVREHKPVLVEILNYKKDGTPFRNAVLVAPIYDDDDELLYFLGSQVEVDDDQPN*GMARRERAAEMLKTLSPRQLEV<br>TTLVASGLRNKEVAARLGLSEKTVKMHRGLVMEKLNLKTSADLVRIAVEAGIA |

| | | |
|---|---|---|
| | *L35CNF/Y136CNF/ M151CNF* | GADDTRVEVQPPAQWVLD*IEASPIASVVSDPRLADNPLIAINQAFTDLTGYSEEECVGRNCRFLAGSGTEPWLTDKIR<br>QGVREHKPVLVEILNYKKDGTPFRNAVLVAPIYDDDDELL*FLGSQVEVDDDQPN*GMARRERAAEMLKTLSPRQLEVT<br>TLVASGLRNKEVAARLGLSEKTVKMHRGLVMEKLNLKTSADLVRIAVEAGIA |
| **Other EL222 variants** | *W31CNF/S140Y* | GADDTRVEVQPPAQ*VLDLIEASPIASVVSDPRLADNPLIAINQAFTDLTGYSEEECVGRNCRFLAGSGTEPWLTDKIR<br>QGVREHKPVLVEILNYKKDGTPFRNAVLVAPIYDDDDELLYFLGYQVEVDDDQPNMGMARRERAAEMLKTLSPRQLEVT<br>TLVASGLRNKEVAARLGLSEKTVKMHRGLVMEKLNLKTSADLVRIAVEAGIA |
| | *L35CNF/L123K* | GADDTRVEVQPPAQWVLD*IEASPIASVVSDPRLADNPLIAINQAFTDLTGYSEEECVGRNCRFLAGSGTEPWLTDKIR<br>QGVREHKPVLVEILNYKKDGTPFRNAVKVAPIYDDDDELLYFLGSQVEVDDDQPNMGMARRERAAEMLKTLSPRQLEVT<br>TLVASGLRNKEVAARLGLSEKTVKMHRGLVMEKLNLKTSADLVRIAVEAGIA |
| | *LOV* | GADDTRVEVQPPAQWVLDLIEASPIASVVSDPRLADNPLIAINQAFTDLTGYSEEECVGRNCRFLAGSGTEPWLTDKIR<br>QGVREHKPVLVEILNYKKDGTPFRNAVLVAPIYDDDDELLYFLGSQVEVDDDQPNMGMARRERAAEMLKTLA |
| | *LOV-W31CNF* | GADDTRVEVQPPAQ*VLDLIEASPIASVVSDPRLADNPLIAINQAFTDLTGYSEEECVGRNCRFLAGSGTEPWLTDKIR<br>QGVREHKPVLVEILNYKKDGTPFRNAVLVAPIYDDDDELLYFLGSQVEVDDDQPNMGMARRERAAEMLKTLA |
| | *LOV-L35CNF* | GADDTRVEVQPPAQWVLD*IEASPIASVVSDPRLADNPLIAINQAFTDLTGYSEEECVGRNCRFLAGSGTEPWLTDKIR<br>QGVREHKPVLVEILNYKKDGTPFRNAVLVAPIYDDDDELLYFLGSQVEVDDDQPNMGMARRERAAEMLKTLA |
| | *AQTRIP (V44I/L55I/A82Q/ V124I)* | GADDTRVEVQPPAQWVLDLIEASPIASIVSDPRLADNPIIAINQAFTDLTGYSEEECVGRNCRFLQGSGTEPWLTDKIR<br>QGVREHKPVLVEILNYKKDGTPFRNAVLIAPIYDDDDELLYFLGSQVEVDDDQPNMGMARRERAAEMLKTLSPRQLEVT<br>TLVASGLRNKEVAARLGLSEKTVKMHRGLVMEKLNLKTSADLVRIAVEAGIA |
| **OTS** | *MjCNFRS* | MDEFEMIKRNTSEIISEEELREVLKKDEKSALIGFEPSGKIHLGHYLQIKKMIDLQNAGFDIIIVLADLHAYLNQKGEL<br>DEIRKIGDYNKKVFEAMGLKAKYVYGSEWMLDKDYTLNVYRLALKTTLKRARRSMELIAREDENPKVAEVIYPIMQVNG<br>PHYLGVDVAVGGMEQRKIHMLARELLPKKVVCIHNPVLTGLDGEGKMSSSKGNFIAVDDSPEEIRAKIKKAYCPAGVVE<br>GNPIMEIAKYFLEYPLTIKRPEKFGGDLTVNSYEELESLFKNKELHPMDLKNAVAEELIKILEPIRKRL |
| | *MjtRNA$_{CUA}$* | ccggcggtagttcagcagggcagaacggcggactCTAaatccgcatggcgctggttcaaatccggcccgccggacca |

* = CNF (4-cyano-L-phenylalanine)

### TABLE S3. Protein quality control.

| Protein | | Yield (mg/L)[a] | FMN/ Protein ( %)[b] | Concentration (mM)[c] | Oligomeric state[d] | Expected mass (Da)[e] | Observed mass (Da)[f] |
|---|---|---|---|---|---|---|---|
| **EL222 WT** | *WT* | 5.1 | 104 | 1.7 | Monomer | 23157.00 | 23157.01 |
| **Single-CNF** | *A18CNF* | 3.2 | 101 | 1.8 | Monomer | 23258.02 | 23258.03 |
| | *W31CNF* | 2.9 | 146 | 1.9 | Monomer | 23142.98 | 23142.99 |
| | *L35CNF* | 4.6 | 150 | 1.3 | Monomer | 23215.98 | 23216.08 |
| | *I36CNF* | 2.4 | 94 | 1.4 | Monomer | 23215.98 | 23216.03 |
| | *A42CNF* | 0.9 | 6 | 0.4 | Dimer | 23258.02 | 15232.59 |
| | *V44CNF* | 0.9 | 0 | 0.5 | Monomer | 23229.99 | 23230.01 |
| | *S46CNF* | 2.7 | 4 | 1.5 | Monomer | 23242.03 | 23242.12 |
| | *N53CNF* | 3.2 | 138 | 1.9 | Monomer | 23215.02 | 23215.13 |
| | *L55CNF* | 1.4 | 24 | 0.8 | Monomer | 23215.98 | 23215.99 |
| | *F62CNF* | 3.2 | 96 | 1.3 | Monomer | 23181.99 | 23182.11 |
| | *Y68CNF* | 4.2 | 121 | 1.8 | Monomer | 23166.00 | 23166.12 |
| | *C78CNF* | 0.9 | 0 | 0.5 | Dimer | 23226.05 | 23226.21 |
| | *F80CNF* | 1.4 | 70 | 0.8 | Monomer | 23181.99 | 23182.10 |
| | *A82CNF* | 2.0 | 70 | 1.5 | Monomer | 23258.02 | 23258.04 |
| | *W89CNF* | 2.7 | 132 | 1.8 | Monomer | 23142.98 | 23143.12 |
| | *T91CNF* | 2.6 | 101 | 1.5 | Monomer | 23228.01 | 23228.12 |
| | *I94CNF* | 1.0 | 63 | 0.8 | Monomer | 23215.98 | 23216.00 |
| | *R95CNF* | 1.7 | 57 | 0.7 | Monomer | 23172.96 | 23173.12 |
| | *I108CNF* | 0.9 | 63 | 0.9 | Monomer | 23215.98 | 23215.96 |
| | *N110CNF* | 1.8 | 5 | 1.0 | Monomer | 23215.02 | 23215.15 |
| | *Y111CNF* | 3.1 | 57 | 1.5 | Monomer | 23166.00 | 26166.14 |
| | *F118CNF* | 3.0 | 89 | 1.1 | Monomer | 23181.99 | 23182.12 |
| | *N120CNF* | 2.1 | 47 | 1.3 | Monomer | 23215.02 | 23215.16 |
| | *V122CNF* | 1.6 | 12 | 1.6 | Monomer | 23229.99 | 23230.00 |
| | *L123CNF* | 2.5 | 219 | 1.2 | Dimer? | 23215.98 | 23216.11 |
| | *V124CNF* | 1.1 | 29 | 1.1 | Monomer | 23229.99 | 23230.01 |
| | *Y128CNF* | 2.9 | 187 | 1.5 | Monomer | 23166.00 | 23166.00 |
| | *Y136CNF* | 2.0 | 155 | 1.1 | Monomer | 23166.00 | 23165.10 |
| | *F137CNF* | 1.8 | 155 | 1.0 | Monomer | 23181.99 | 23182.15 |
| | *S140CNF* | 1.1 | 140 | 1.1 | Multimer | 23242.03 | 23242.18 |
| | *Q141CNF* | 1.4 | 92 | 0.8 | Dimer | 23201.00 | 23201.16 |
| | *Q148CNF* | 2.3 | 126 | 1.7 | Monomer | 23201.00 | 23201.15 |
| | *P149CNF* | 0.4 | 67 | 0.6 | Monomer | 23232.01 | 23222.14 |
| | *M151CNF* | 3.1 | 108 | 0.9 | Monomer | 23198.02 | 23198.15 |
| | *M153CNF* | 2.5 | 172 | 1.5 | Monomer | 23198.02 | 23197.98 |
| | *M162CNF* | 0.7 | 188 | 0.4 | Monomer | 23198.02 | 23198.04 |
| | *L166CNF* | 1.5 | 101 | 1.5 | Monomer | 23215.98 | 23216.11 |
| | *L181CNF* | 2.2 | 150 | 1.4 | Monomer | 23215.98 | 23215.99 |
| | *G191CNF* | 2.0 | 133 | 1.5 | Monomer | 23272.04 | 23272.17 |
| | *M199CNF* | 2.6 | 74 | 1.5 | Monomer | 23198.02 | 23198.09 |
| | *M205CNF* | 2.8 | 91 | 1.1 | Monomer | 23198.02 | 23198.09 |
| | *L210CNF* | 3.4 | 106 | 1.9 | Monomer | 23215.98 | 23218.99 |
| | *T212CNF* | 2.8 | 132 | 1.3 | Monomer | 23228.01 | 23228.16 |
| | *L216CNF* | 3.4 | 131 | 1.5 | Monomer | 23215.98 | 23216.10 |
| | *I225CNF* | 2.4 | 97 | 1.5 | Monomer | 23215.98 | 23216.12 |
| **Double-CNF** | *W31CNF/L35CNF* | 0.73 | 109 | 1.2 | Monomer | 23201.96 | 23201.10 |
| | *W31CNF/M151CNF* | 2.0 | 142 | 1.5 | Monomer | 23184.00 | 23183.15 |
| | *L35CNF/M151CNF* | 2.5 | 108 | 1.5 | Monomer | 23257.00 | 23256.13 |
| **Triple-CNF[f]** | *L35CNF/Y136CNF/M151CNF* | 0.31 | 164 | 1.5 | Monomer | 23266.00 | 23266.13 |
| | *W31CNF/L35CNF/M151CNF* | 0.94 | 126 | 1.1 | Monomer | 23242.98 | 23243.13 |
| **Other EL222 mutants** | *W31CNF/S140Y* | 1.3 | 77 | 1.3 | Multimer | 23219.01 | 23219.21 |
| | *L35CNF/L123K* | 0.9 | 165 | 1.2 | Dimer? | 23231.00 | 23229.11 |
| | *LOV* | 3.2 | 24 | 1.1 | Monomer | 16726.41 | 16778.41 |
| | *LOV W31CNF* | 2.6 | 34 | 1.2 | Monomer | 16795.53 | 16764.39 |
| | *LOV L35CNF* | 1.8 | 30 | 1.8 | Monomer | 16837.39 | 16838.50 |
| | *AQTRIP* | 2.6 | 76 | 1.2 | Monomer | 23242.19 | 23241.05 |

[a] All variants were expressed in TB media except W89CNF and I94CNF (both glucose-based minimal media)
[b] FMN concentrations were estimated by UV/Vis spectroscopy while protein concentrations were estimated by FTIR.
[c] The reported concentration applies to FTIR experiments. For UV/Vis experiments, proteins were diluted to a concentration between 10-40 µM.
[d] Estimated by size-exclusion chromatography (elution volumes interpolated in a calibration curve with proteins of known molecular weight) in the dark state.
[e] Expected monoisotopic masses were computed from https://web.expasy.org/peptide_mass/. Observed masses were determined by MS.
[f] The two triple mutants were expressed in C321.ΔA.exp(DE3). All other constructs were expressed in BL21(DE3).

#### TABLE S4. Extracted peak parameters of the CNF bands.

| | DARK | | | LIT | | |
|---|---|---|---|---|---|---|
| **Variant** | $\nu_{max}$ [a] **(cm$^{-1}$)** | $\lvert\vec{F}\rvert$ [b] **(MV/cm)** | $\Delta\nu_{HB}$ [c] **(cm$^{-1}$)** | $\nu_{max}$ [a] **(cm$^{-1}$)** | $\lvert\vec{F}\rvert$ [b] **(MV/cm)** | $\Delta\nu_{HB}$ [c] **(cm$^{-1}$)** |
| *A18CNF* | 2235.8 | -27 | 10 | 2235.7 | -27 | 9 |
| *W31CNF* | 2236.8 | -37 | 12 | 2235.4 | -36 | 11 |
| *L35CNF* | 2232.6 | -66 | 14 | 2233.7 | -68 | 15 |
| *I36CNF* | 2229.9 | -47 | 7 | 2230.2 | -46 | 7 |
| *N53CNF* | 2235.5 | -37 | 11 | 2235.1 | -37 | 11 |
| *L55CNF* | 2237.1 | -58 | 17 | 2237.1 | -58 | 17 |
| *F62CNF* | 2231.1 | -62 | 11 | 2231.2 | -62 | 12 |
| *Y68CNF* | 2236.6 | -54 | 15 | 2236.4 | -49 | 14 |
| *F80CNF* | 2236.2 | -51 | 14 | 2236.0 | -53 | 15 |
| *A82CNF* | 2233.4 | -76 | 16 | 2232.8 | -75 | 16 |
| *T91CNF* | 2234.3 | -52 | 13 | 2232.6 | -54 | 12 |
| *I94CNF* | 2227.0 | -76 | 10 | 2229.3 | -73 | 12 |
| *R95CNF* | 2234.5 | -80 | 18 | 2232.8 | -74 | 15 |
| *I108CNF* | 2229.9 | -54 | 9 | 2230.1 | -52 | 9 |
| *Y111CNF* | 2234.7 | -32 | 9 | 2234.7 | -33 | 10 |
| *F118CNF* | 2230.1 | -51 | 8 | 2230.0 | -54 | 9 |
| *N120CNF* | 2223.4 | -40 | 0 | 2222.8 | -40 | -1 |
| *V122CNF* | 2230.2 | -88 | 15 | 2230.3 | -88 | 16 |
| *L123CNF* | 2234.3 | -71 | 16 | 2228.5 | -85 | 13 |
| *V124CNF* | 2231.5 | -54 | 10 | 2230.5 | -53 | 9 |
| *Y128CNF* | 2232.6 | -92 | 19 | 2232.5 | -92 | 19 |
| *Y136CNF* | 2235.3 | -83 | 20 | 2234.3 | -87 | 19 |
| *F137CNF* | 2235.1 | -108 | 24 | 2233.2 | -123 | 25 |
| *S140CNF* | 2232.7 | -84 | 17 | 2234.0 | -85 | 19 |
| *Q141CNF* | 2231.8 | -99 | 19 | 2228.2 | -98 | 15 |
| *Q148CNF* | 2235.3 | -45 | 13 | 2234.0 | -38 | 10 |
| *P149CNF* | 2235.3 | -16 | 7 | 2234.7 | -15 | 6 |
| *M151CNF* | 2234.7 | -52 | 13 | 2234.3 | -52 | 13 |
| *M153CNF* | 2235.9 | -22 | 9 | 2235.9 | -22 | 9 |
| *M162CNF* | 2235.8 | -84 | 20 | 2235.6 | -85 | 20 |
| *L166CNF* | 2232.6 | -32 | 7 | 2234.3 | -36 | 10 |
| *L181CNF* | 2236.4 | -58 | 16 | 2236.5 | -57 | 16 |
| *G191CNF* | 2235.1 | -39 | 11 | 2235.1 | -39 | 11 |
| *M199CNF* | 2235.3 | -21 | 8 | 2235.3 | -19 | 8 |
| *M205CNF* | 2233.1 | -81 | 17 | 2233.1 | -79 | 17 |
| *L210CNF* | 2233.9 | -45 | 11 | 2233.3 | -44 | 10 |
| *T212CNF* | 2235.3 | -30 | 10 | 2235.0 | -32 | 10 |
| *L216CNF* | 2229.0 | -36 | 5 | 2229.1 | -42 | 6 |
| *I225CNF* | 2235.5 | -39 | 12 | 2235.5 | -37 | 11 |

[a] Peak maxima.
[b] Environmental electric field inferred from the peak areas.
[c] Frequency shift due to H-bonding inferred from the electric fields and field-frequency calibration curves (see **Note S3**).

### TABLE S5: CNF rotamers.

| Variant | $X_1$ (°)[a] | $X_2$ (°)[a] |
|---|---|---|
| *A18CNF* | -55 | 103.1 |
| *W31CNF* | -173.2 | 73.5 |
| *L35CNF* | -180 | 77.8 |
| *I36CNF* | -65.4 | 107.7 |
| *A42CNF* | -176.1 | 71.9 |
| *V44CNF* | 65.8 | 91.7 |
| *S46CNF* | -63.1 | 97.7 |
| *N53CNF* | -57.2 | 110.2 |
| *L55CNF* | -71.3 | 103 |
| *F62CNF* | -178.1 | 77.2 |
| *Y68CNF* | -69.6 | 93.7 |
| *C78CNF* | -67.4 | 99.6 |
| *F80CNF* | -70.4 | 106.7 |
| *A82CNF* | -71.5 | 100.1 |
| *W89CNF* | -73.5 | 105.9 |
| *T91CNF* | -179.1 | 78.3 |
| *I94CNF* | 177 | 79.6 |
| *R95CNF* | -74 | 109.6 |
| *I108CNF* | -66.5 | 96 |
| *Y111CNF* | -61.4 | 100.4 |
| *F118CNF* | 61 | 93.2 |
| *N120CNF* | -66.8 | 92 |
| *V122CNF* | -67.3 | 94 |
| *L123CNF* | -67.8 | 95.3 |
| *V124CNF* | -177.4 | 77.7 |
| *Y128CNF* | -66.5 | 96 |
| *Y136CNF* | 64.1 | 92.2 |
| *F137CNF* | -64.2 | 90.8 |
| *S140CNF* | -63 | 91.1 |
| *Q141CNF* | -61 | 90.7 |
| *Q148CNF* | -176.2 | 77.2 |
| *P149CNF* | -71.3 | 103 |
| *M151CNF* | -63.2 | 116.2 |
| *M153CNF* | -70 | 106 |
| *M162CNF* | -180 | 77.8 |
| *L166CNF* | -71.5 | 100.1 |
| *L181CNF* | -67.5 | 99.6 |
| *G191CNF* | -61.1 | 101.2 |
| *M199CNF* | 177 | 79.6 |
| *M205CNF* | -178.1 | 77.2 |
| *L210CNF* | -69.8 | 97.8 |
| *T212CNF* | -61.4 | 93.4 |
| *L216CNF* | 177 | 79.6 |
| *I225CNF* | -66.6 | 99.6 |

[a] $X_1$ is the torsion angle around the Cα-Cβ axes defined by the atoms N-Cα-Cβ-Cγ.
[b] $X_2$ is the torsion angle around the Cβ-Cγ axes defined by the atoms Cα-Cβ-Cγ-Cδ.

### TABLE S6: Kinetic parameters for the lit-to-dark transition of EL222.

Lifetimes (τ) at 20°C and $H_2O$-based buffer (except where indicated otherwise) corresponding to the peaks with the largest *D* values calculated from lifetime distribution analysis using the maximum entropy method in different spectral regions. Lifetimes are indicated as mean ± standard deviation (n=3). Errors below two seconds are omitted. If more than one peak is observed, the relative areas are indicated in parenthesis

| **Variant** | | $\tau_{FMN}$ **(s)** [a] | $\tau_{Amide}$ **(s)** [b] | $\tau_{CNF}$ **(s)** [c] |
|---|---|---|---|---|
| **EL222** | *WT* | 41 | 57±3 | N.P. |
| | *WT ($D_2O$)* | 56 (19%) /177±10 (70%) | 223±4 | N.P. |
| | *LOV* | 23 | 34 | N.P. |
| | *LOV ($D_2O$)* | 104 | 42±2 (20%) / 142±4 (64%) | N.P. |
| **1x CNF** | *A18CNF* | 38 | | |
| | *W31CNF* | 85±4 (66%) / 455±17 (30%) | 40±3 (35%) / 221±9 (34%) | 41 (64%) / 196±19 (36%) |
| | *L35CNF* | 39 | 104±4 | 24 (39%) / 98±3 (54%) |
| | *I36CNF* | 22 | | |
| | *N53CNF* | 48 | 53±3 | 46 |
| | *L55CNF* | 40±13 | | |
| | *F62CNF* | 8 | | |
| | *Y68CNF* | 22 | | |
| | *F80CNF* | 30 | | |
| | *A82CNF* | 9 | | |
| | *W89CNF* | 45±14 | | |
| | *T91CNF* | 117±4 | | |
| | *I94CNF* | 145±11 | | |
| | *R95CNF* | 61 | | |
| | *I108CNF* | 39 | | |
| | *Y111CNF* | 72 | | |
| | *F118CNF* | 18 | | |
| | *N120CNF* | 368±4 | | |
| | *L123CNF* | 86 | | |
| | *Y128CNF* | 47 | | |
| | *Y136CNF* | 44±3 | 59±2 | 63±4 |
| | *F137CNF* | 29 | | |
| | *S140CNF* | 298±18 | | |
| | *Q141CNF* | 162±17 | | |
| | *Q148CNF* | 29 | | |
| | *P149CNF* | 40 | | |
| | *M151CNF* | 39 | 60±2 | 78±9 |
| | *M153CNF* | 36 | | |
| | *M162CNF* | 62±34 | | |
| | *L166CNF* | 22 | | |
| | *L181CNF* | 42 | | |
| | *G191CNF* | 32 | | |
| | *M199CNF* | 36 | | |
| | *M205CNF* | 24 | | |
| | *L210CNF* | 21 | | |
| | *T212CNF* | 18 | | |
| | *L216CNF* | 21 (30%) /44 (59%) | 47±4 | 40 |
| | *I225CNF* | 38 | | |
| **2xCNF** | *W31CNF/L35CNF* | 97±6 | 87±6 | 88±3 |
| | *W31CNF/M151CNF* | 7 (21%) / 21 (65%) | 32±3 (66%) / 324±23 (34%) | 28±2 (52%) / 358±23 (48%) |
| | *L35CNF/M151CNF* | 32±2 (63%) / 58±4 (21%) | 177±2 | 39±4 (43%) / 181±3 (48%) |
| | *Y136CNF/M151CNF* | 19±2 (23%) / 44±4 (66%) | 45±2 (70%) /552±11 (28%) | 53±5 (40%) / 480±75 (52%) |
| **3x CNF** | *L35CNF/Y136CNF/M151CNF* | 29 (52%) / 132±3 (29%) | 105±19 | 99±10 |
| | *W31CNF/L35CNF/M151CNF* | 93±8 (50%) /672±170 (50%) | 112±26 | 115±3 (55%) / 588±22 (30%) |

[a] 320-450 nm. [b] 1750-1500 $cm^{-1}$. [c] 2270-2200 $cm^{-1}$.

N.P. indicates a measurement that is not possible due to the sample lacking absorption bands in the target region.

### TABLE S7: Activation energies for the lit-to-dark transition of EL222.

Activation energies ($E_a$) in $H_2O$-based buffers (except where indicated otherwise) calculated from the slopes of Arrhenius plots (natural logarithm of the kinetic rate constants, taken as the inverse of the lifetimes, as a function of the temperature). The used lifetimes were extracted from the peaks with the largest *D* values calculated from lifetime distribution analysis using the maximum entropy method in different spectral regions. Only peaks with relative area larger than 15% were analyzed. Values in parenthesis were calculated from the slowest kinetic events in those cases were two components were found at all temperatures.

| **Variant** | $\boldsymbol{E_a^{FMN}}$ **(kcal/mol)** [a] | $\boldsymbol{E_a^{Amide}}$ **(kcal/mol)** [b] | $\boldsymbol{E_a^{CNF}}$ **(kcal/mol)** [c] |
|---|---|---|---|
| *wt EL222* | 16 | 18 | N.P |
| *wt EL222 ($D_2O$)* | 14 | 18 | N.P. |
| *EL222 W31CNF* | 17 (19) | 14 (13) | 12 |
| *EL222 L35CNF* | 16 | 12 (13) | 13 (16) |
| *EL222 M151CNF* | 12 | 17 | ~0 |

[a] 320-450 nm. [b] 1750-1500 $cm^{-1}$. [c] 2270-2200 $cm^{-1}$.

N.P. indicates a measurement that is not possible due to the sample lacking absorption bands in the target region.

### TABLE S8. Photoconversion yields.

| **Variant** | $f_{A_{390}}$ [a] |
|---|---|
| *wt EL222* | 0.83 |
| *wt EL222 ($D_2O$)* | 0.91 |
| *A18CNF* | 0.93 |
| *W31CNF* | 0.83 |
| *L35CNF* | 0.92 |
| *I36CNF* | 0.89 |
| *N53CNF* | 0.90 |
| *L55CNF* | 0.96 |
| *F62CNF* | 0.88 |
| *Y68CNF* | 0.85 |
| *F80CNF* | 0.51 |
| *A82CNF* | 0.99 |
| *T91CNF* | 0.81 |
| *I94CNF* | 1.00 |
| *R95CNF* | 0.92 |
| *I108CNF* | 0.95 |
| *Y111CNF* | 0.89 |
| *F118CNF* | 0.90 |
| *N120CNF* | 0.96 |
| *V122CNF* | 0.87 |
| *L123CNF* | 1.00 |
| *V124CNF* | 0.90 |
| *Y128CNF* | 0.91 |
| *Y136CNF* | 0.91 |
| *F137CNF* | 0.91 |
| *S140CNF* | 0.97 |
| *Q141CNF* | 0.92 |
| *Q148CNF* | 0.82 |
| *P149CNF* | 0.89 |
| *M151CNF* | 0.82 |
| *M153CNF* | 0.87 |
| *M162CNF* | 0.87 |
| *L166CNF* | 0.81 |
| *L181CNF* | 0.87 |
| *G191CNF* | 0.91 |
| *M199CNF* | 0.93 |
| *M205CNF* | 0.99 |
| *L210CNF* | 0.90 |
| *T212CNF* | 0.82 |
| *L216CNF* | 0.88 |
| *I225CNF* | 0.91 |

[a] Fraction of EL222 proteins in the lit state upon illumination based on FMN absorbance (FMN-C78 adduct state or, equivalently, $A_{390}$ species, see **Note S4**). All data were taken in $H_2O$-based buffer except otherwise indicated.

### TABLE S9: Regularization values.

Optimum regularization parameter ($\lambda_{opt}$), given as $log_{10}(\lambda_{opt})$, determined using the L-curve method.

| Variant | | $\lambda_{opt}^{FMN}$ [a] | $\lambda_{opt}^{Amide}$ [b] | $\lambda_{opt}^{CNF}$ [c] |
|---|---|---|---|---|
| **EL222** | *WT* | -6.9 | -6.2 | N.P. |
| | *WT ($D_2O$)* | -5.8 | -6.0 | N.P. |
| | *LOV* | -6.5 | -5.8 | N.P. |
| | *LOV ($D_2O$)* | -6.5 | -6.7 | N.P. |
| **1x CNF** | *W31CNF* | -6.3 | -4.6 | -6.1 |
| | *L35CNF* | -6.9 | -4.7 | -6.6 |
| | *N53CNF* | -6.6 | -5.2 | -6.1 |
| | *Y136CNF* | -6.0 | -4.6 | -6.5 |
| | *M151CNF* | -6.4 | -5.3 | -6.4 |
| | *L216CNF* | -6.6 | -4.7 | -6.0 |
| **2xCNF** | *W31CNF/L35CNF* | -5.7 | -6.4 | -6.4 |
| | *W31CNF/M151CNF* | -6.9 | -5.4 | -6.4 |
| | *L35CNF/M151CNF* | -6.3 | -5.0 | -6.4 |
| | *Y136CNF/M151CNF* | -5.5 | -5.4 | -6.0 |
| **3x CNF** | *L35CNF/Y136CNF/M151CNF* | -5.1 | -5.2 | -5.9 |
| | *W31CNF/L35CNF/M151CNF* | -4.0 | -6.2 | -6.4 |

[a] 320-450 nm. [b] 1750-1500 $cm^{-1}$. [c] 2270-2200 $cm^{-1}$.

N.P. indicates a measurement that is not possible due to the sample lacking absorption bands in the target region.

### FIGURE S1.

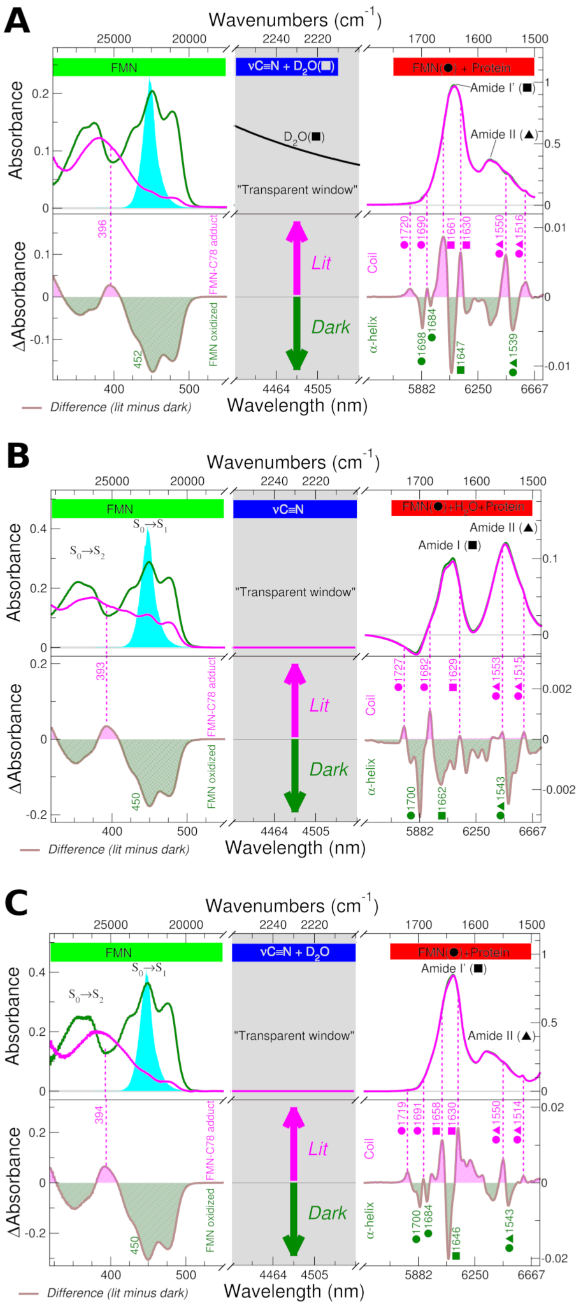

**Electronic and vibrational spectra of wt EL222 and its isolated LOV domain in different solvents.** UV/Visible absorbance (top left) and difference (bottom left) spectra of dark (green) and lit (magenta) states of EL222. FTIR absorbance (top right) and difference (bottom right) spectra of dark (green) and lit (magenta) states of EL222. The main bands in the difference spectra are labeled and assigned to either FMN or protein. In the middle panel, the solvent absorbance is indicatied. Notice that $D_2O$ overlaps and masks the signal arising from the cyano moiety of CNF. **(A)** wt EL222 ($D_2O$), **(B)** EL222-LOV ($H_2O$), **(C)** EL222-LOV ($D_2O$).

#### FIGURE S2.

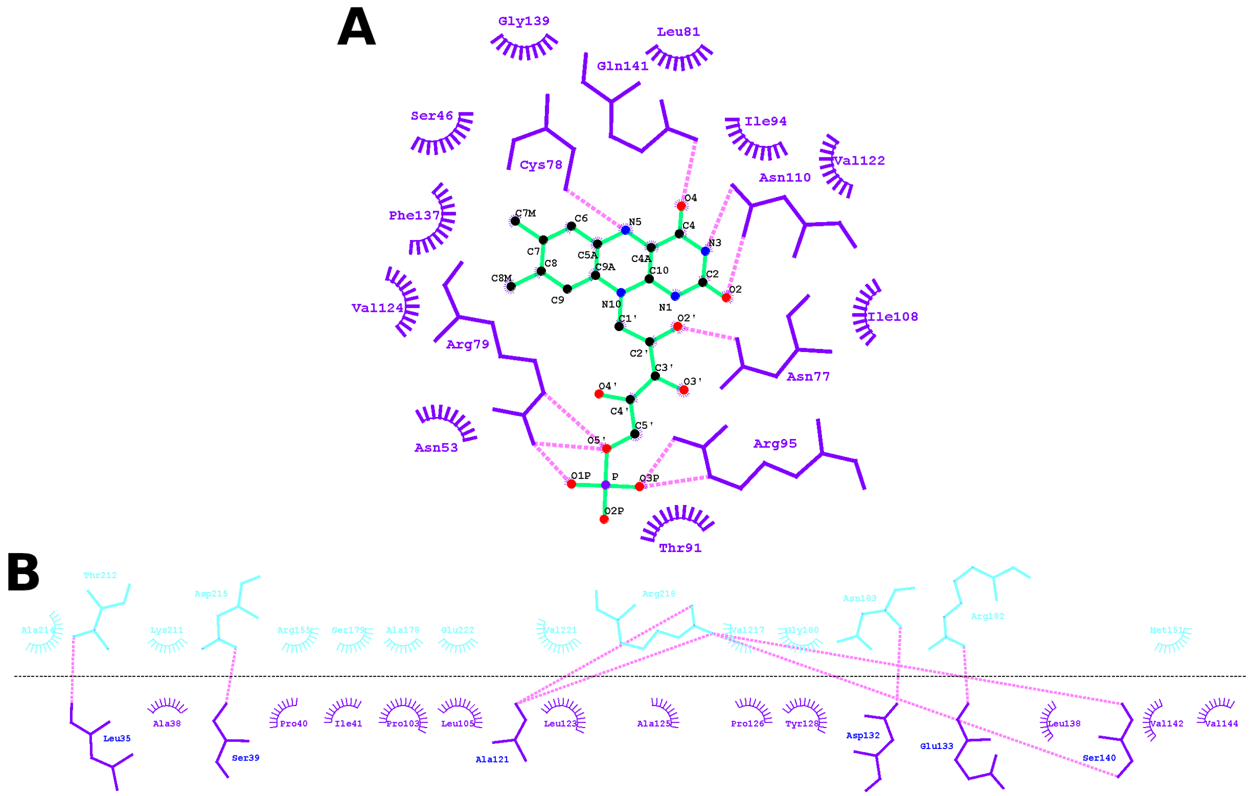

**Schematic representation of protein-chromophore and intra-protein interactions taking place in the dark state of EL222. (A)** Interactions between the LOV domain and the FMN cofactor. **(B)** Interactions between the LOV and HTH domains. The sketches are based on the crystal structure of EL222 (3P7N) (*2*) and rendered with LigPlot (*97*). FMN is shown in green, LOV domain in violet, HTH domain in cyan and hydrogen bonds as magenta dotted lines.

### FIGURE S3.

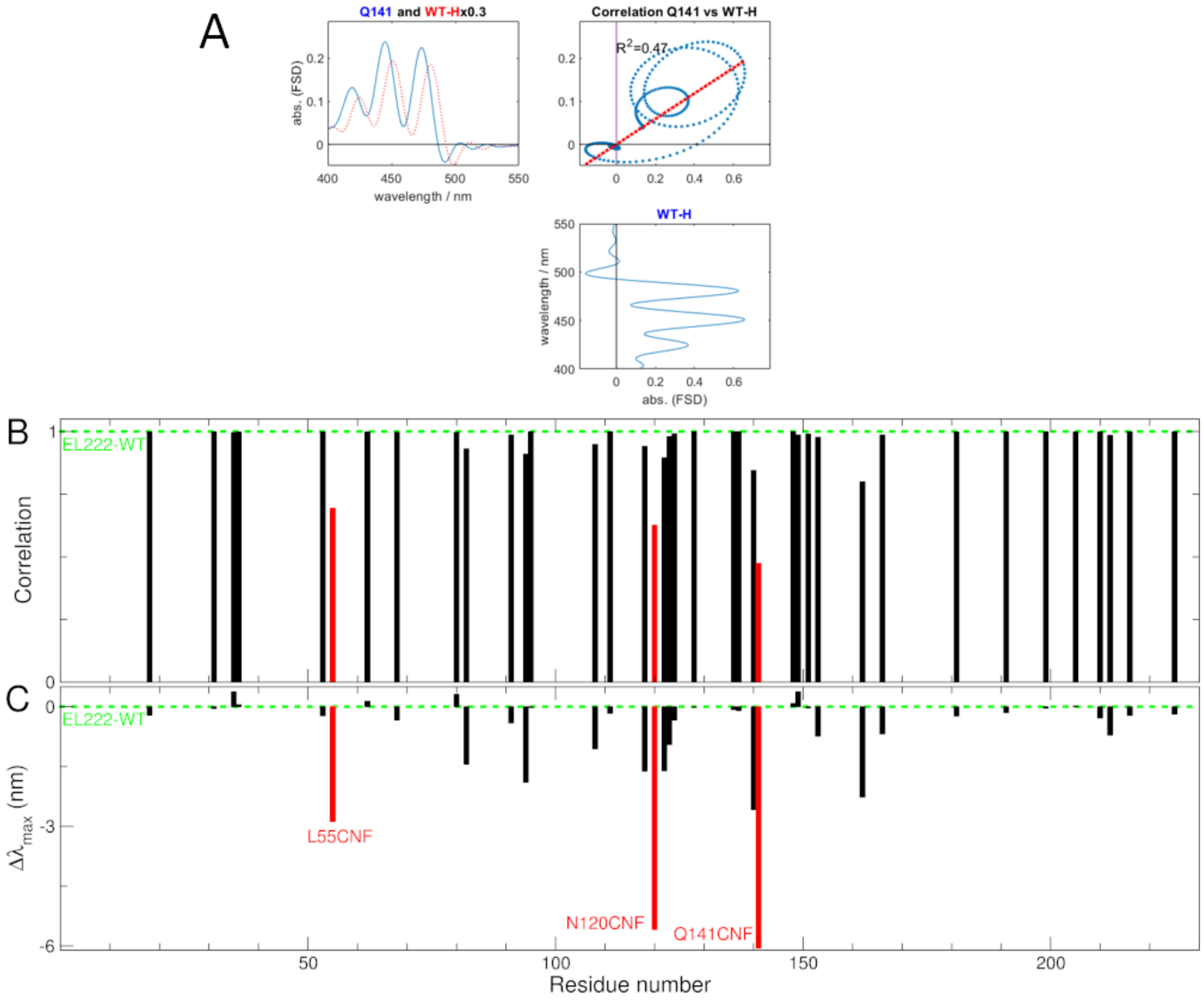

**Comparison between the UV/Vis dark spectra of different CNF mutants and the wt**. **(A)** The similarity between band-narrowed UV/Vis spectrum of Q141CNF and wt was quantified by the correlation coefficient of a plot of the spectrum of Q141CNF *vs.* wt EL222 ($R^2$=1 for identical spectra, $R^2$=0 for spectra without any correlation between them) as previously done (*98*). **(B)** Correlation coefficient for different variants with respect to wt EL222. Variants with a correlation below 0.7, highlighted in red, display significant spectral alterations. **(C)** Shift of the maximum absorption wavelength ($\Delta\lambda_{max}$) of the main peak in the UV/Vis spectrum between different variants and wt EL222 (451 nm). Variants with a shift above 3 nm are labeled in red.

#### FIGURE S4.

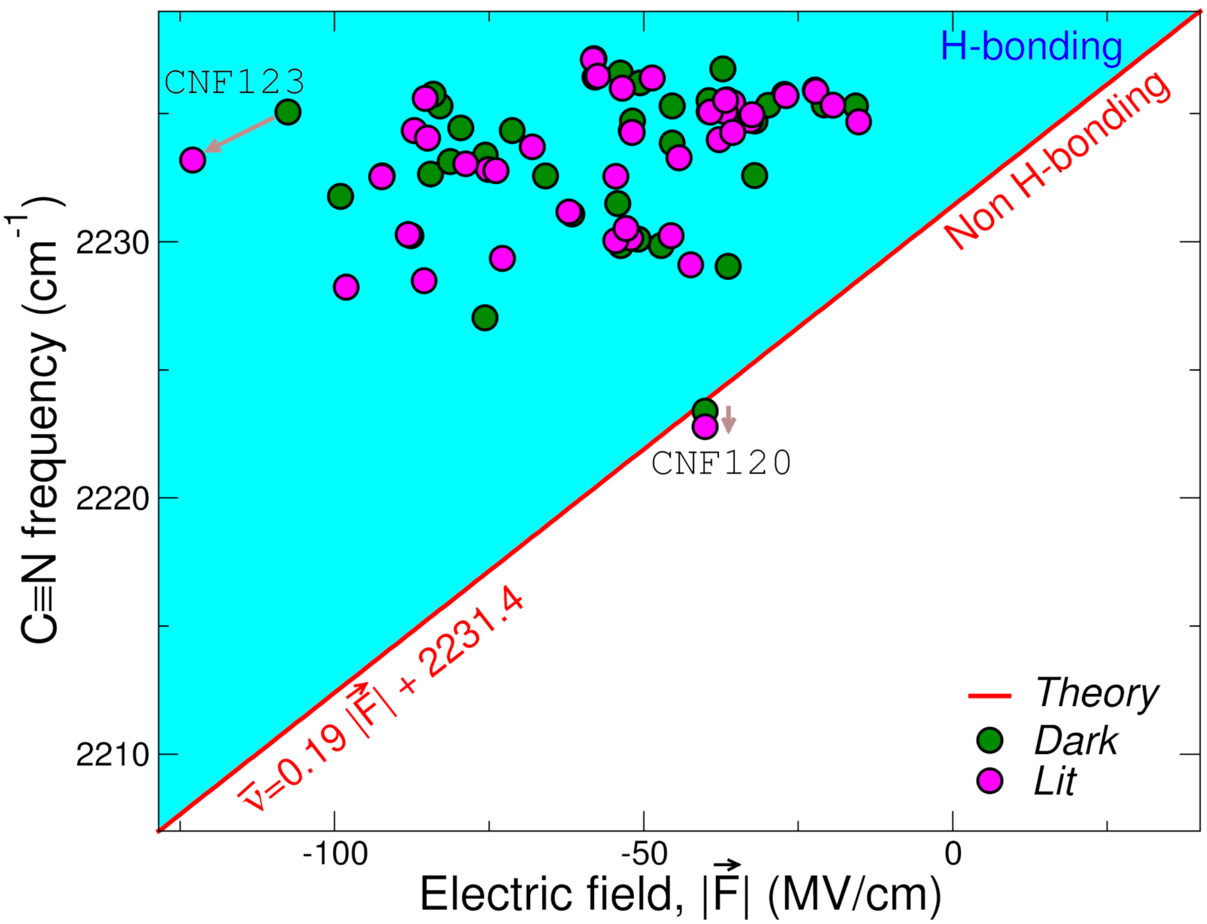

**Peak frequency of CNF bands as a function of solvent electric field.** Experimental data of 39 EL222 variants in the dark (green circles) and lit (magenta circles) states. C≡N frequencies have been taken from the peak maxima ($\nu_{max}$) while electric fields have been estimated from the peak areas (Table S4). The red line is the aprotic (non H-bonding) solvent field-frequency calibration curve for ortho-tolunitrile using polarizable force fields (EQ. 5, shown in the graph) (*45*). The cyan-shaded area denotes the blue-shifts caused by H-bonding. Two residue positions with extreme behaviors are shown. On one hand, CNF120 follows the theoretical prediction. On the other hand, CN123 displays the largest deviation, thus suggesting a strong H-bonding environment around the nitrile probe.

#### FIGURE S5.

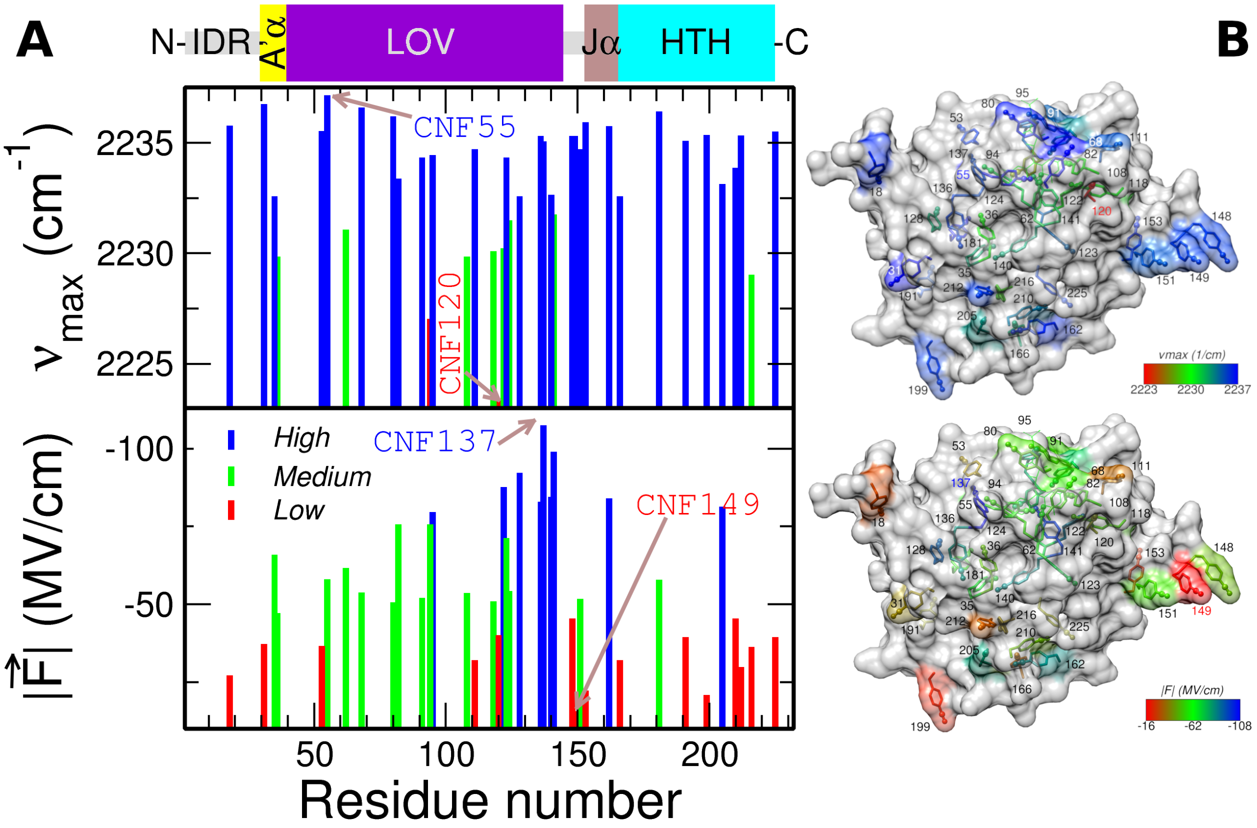

**Mapping site-specific parameters reported by the CNF probe of dark state EL222 variants in 1D and 3D.** **(A)** Plots of maximum absorbance ($\nu_{max}$, top), and electric field ($|\vec{F}|$, bottom) as a function of the residue number. A scheme of different EL222 regions in its dark state conformation is shown on top. From the N- to the C-terminus we have: an intrinsically dis-ordered region (IDR, grey-colored), the A'α extension (yellow), the LOV domain (purple), another IDR (grey), the Jα bridging helix (brown), and the HTH domain (cyan). **(B)** Structural models of dark-adapted EL222 (PDB: 3P7N) containing 39 CNF residues (see Note S2) color-coded according to the same three criteria as in panel A: $\nu_{max}$ (top), and $|\vec{F}|$ (bottom). In both panels, the residues with the lowest and highest values are highlighted in red and blue, respectively.

#### FIGURE S6.

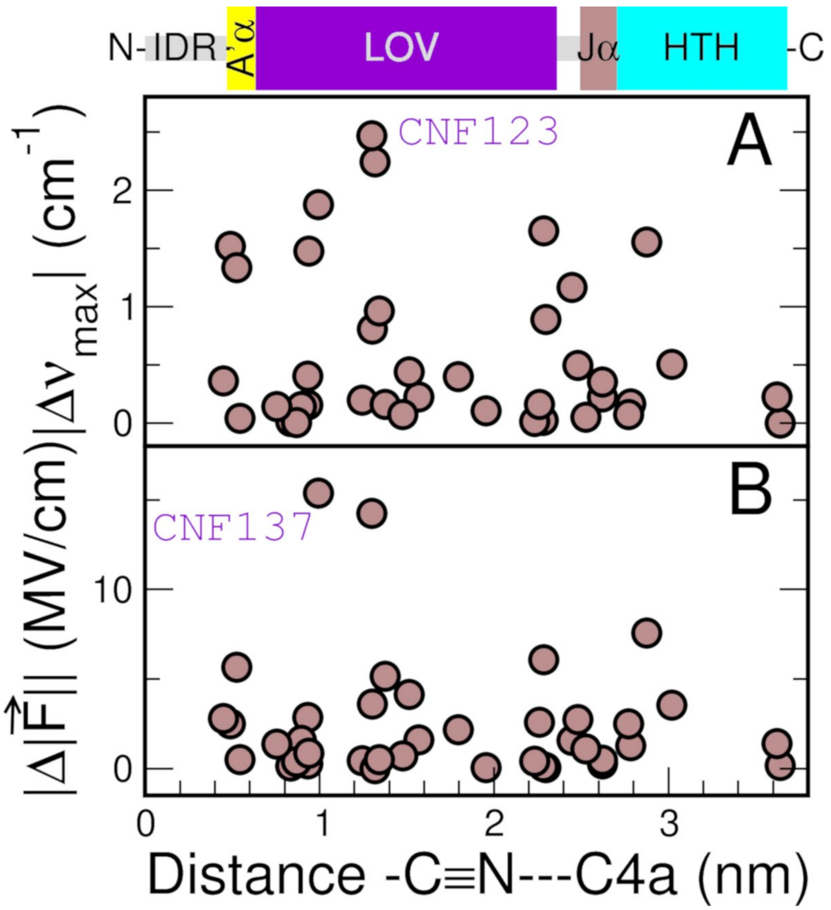

**Light-induced changes in CNF spectra and derived parameters as a function of CNF-to-FMN distance.** Distances are calculated between the N atom of the nitrile moiety of the CNF residue and C4a atom of FMN of structural models of EL222 in the dark state (Note S2). **(A)** Shift in C≡N absorption maxima between dark and lit states. **(B)** Change in electric field, calculated from the areas of the C≡N absorption bands, between dark and lit states. The residues with the largest spectral changes are indicated. A scheme of different EL222 regions in its dark state conformation is shown on top. From the N- to the C-terminus we have: an intrinsically dis-ordered region (IDR, grey-colored), the A'α extension (yellow), the LOV domain (purple), another IDR (grey), the Jα bridging helix (brown), and the HTH domain (cyan). Note the absence of any correlation between the changes in the nitrile vibration and the distance to the FMN.

### FIGURE S7.

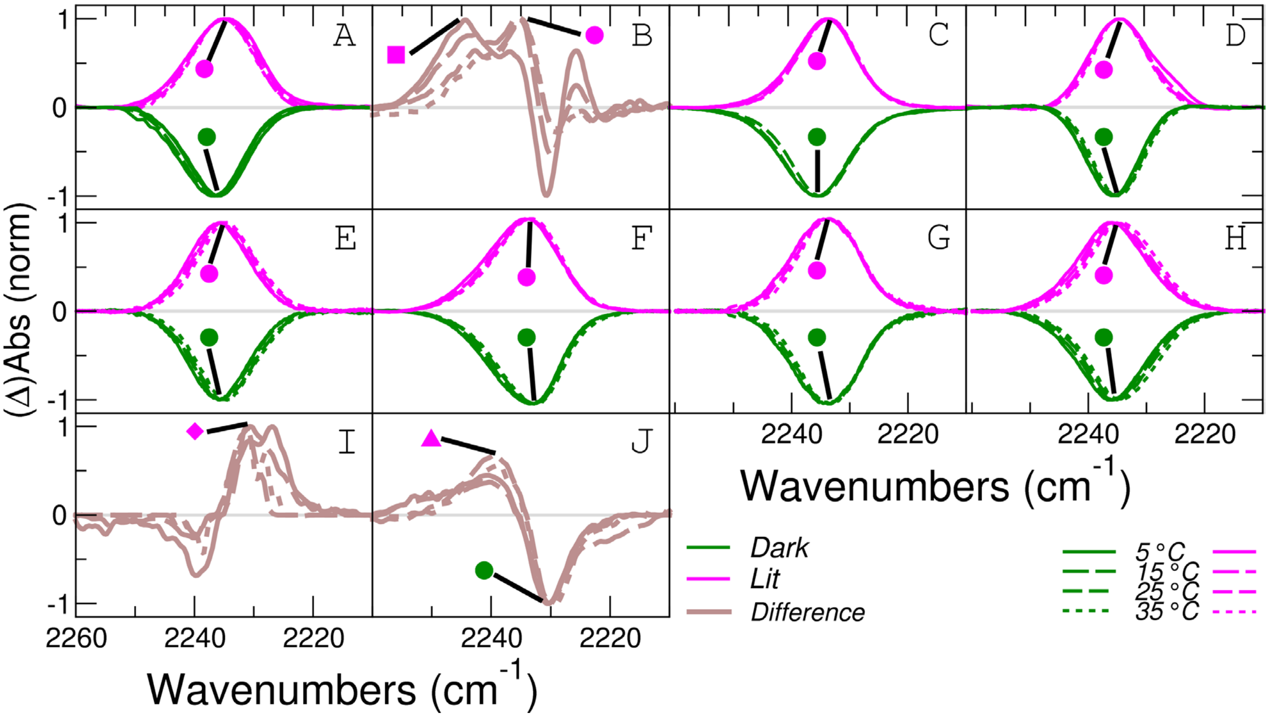

**FTLS analyses of EL222 variants.** C≡N absorption spectra at four different temperatures (5-35 °C in 10 °C steps) were used to calculate frequency-temperature line slope (FTLS) plots. In general, absolute spectra of dark (green lines) and lit (magenta lines) samples were employed. Exceptionally, difference spectra (brown lines) were needed in the case of EL222 L35CNF variant in order to separate the two species found under continuous illumination. The sign of dark spectra has been inverted to facilitate their visualization. Notice that negative bands in difference spectra correspond to the dark state and positive bands to the lit state. The tracked bands are indicated with symbols. In panel I) the negative band is not present at all temperatures and therefore no dark-state band could be tracked. The EL222 variants are as follows. A) EL222 W31CNF. B) EL222 L35CNF (photostationary spectra). C) EL222 Y136CNF. D) EL222 M151CNF. E) LOV-W31CNF. F) LOV-L35CNF. G) EL222 W31CNF/S140Y. H) EL222 L35CNF/L123K. I) “Fast” transient component of EL222 L35CNF (CNF35-A). J) “Slow” transient component of EL222 L35CNF (CNF35-B).

#### FIGURE S8.

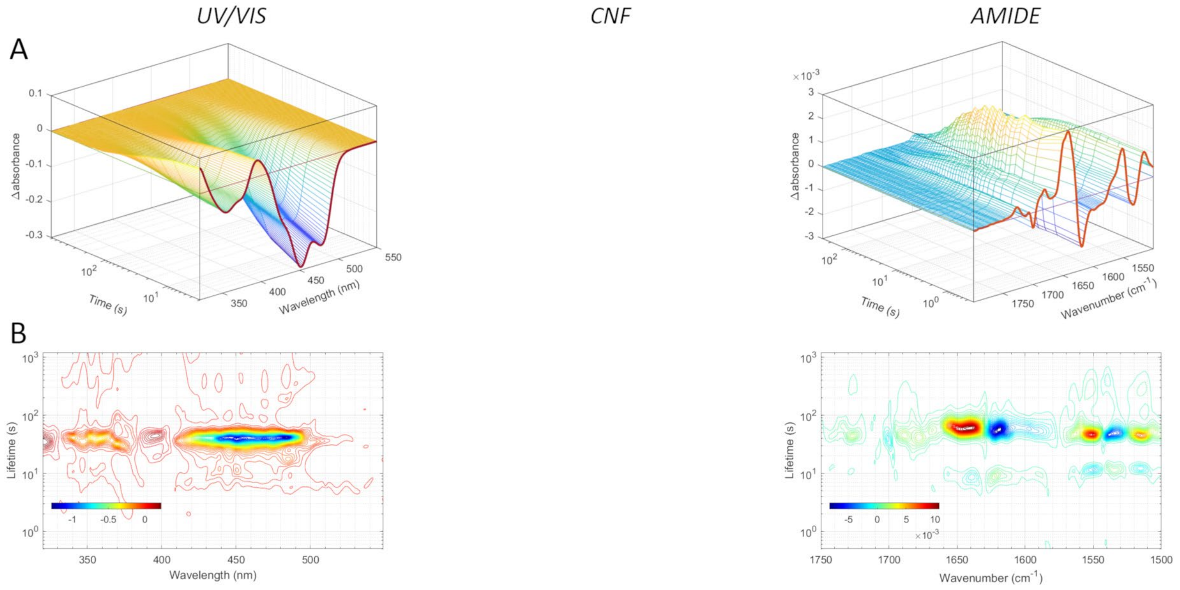

**Time-resolved UV/VIS/IR spectroscopy of wt EL222 in $H_2O$. (A)** 3D pre-processed datasets of differential absorbance as a function of time and frequency. **(B)** 2D lifetime distribution plots using the maximum entropy method. The left, middle and right panels show the FMN (320-550 nm) and amide (1750-1500 cm$^{-1}$) probes, respectively.

#### FIGURE S9.

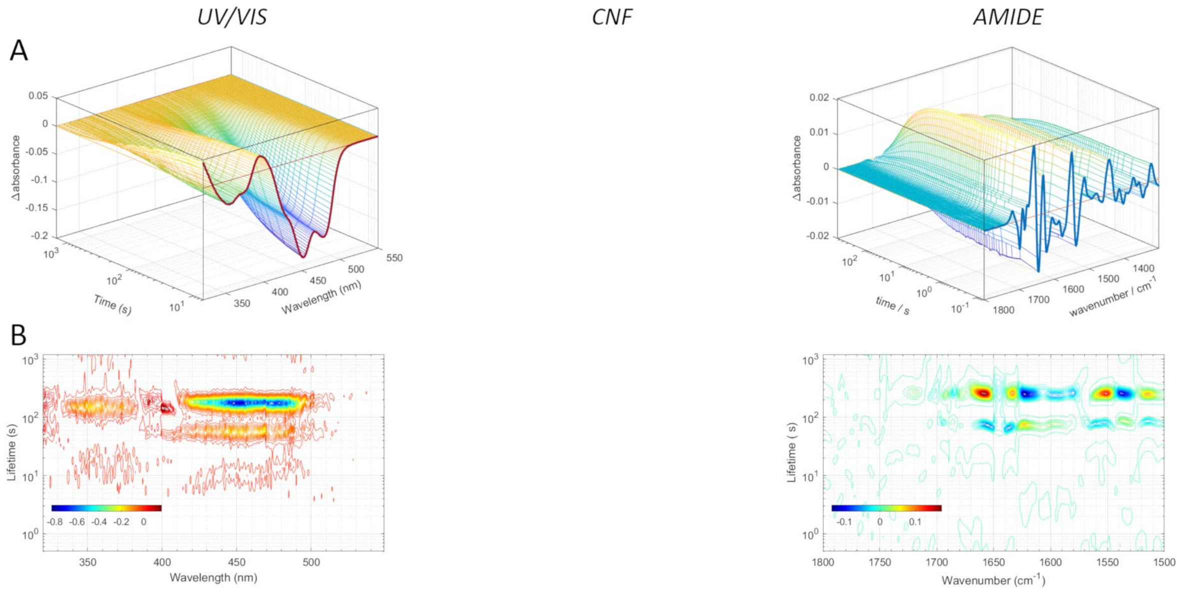

**Time-resolved UV/VIS/IR spectroscopy of wt EL222 in $D_2O$. (A)** 3D pre-processed datasets of differential absorbance as a function of time and frequency. **(B)** 2D lifetime distribution plots using the maximum entropy method. The left, middle and right panels show the FMN (320-550 nm) and amide (1750-1500 cm$^{-1}$) probes, respectively.

#### FIGURE S10.

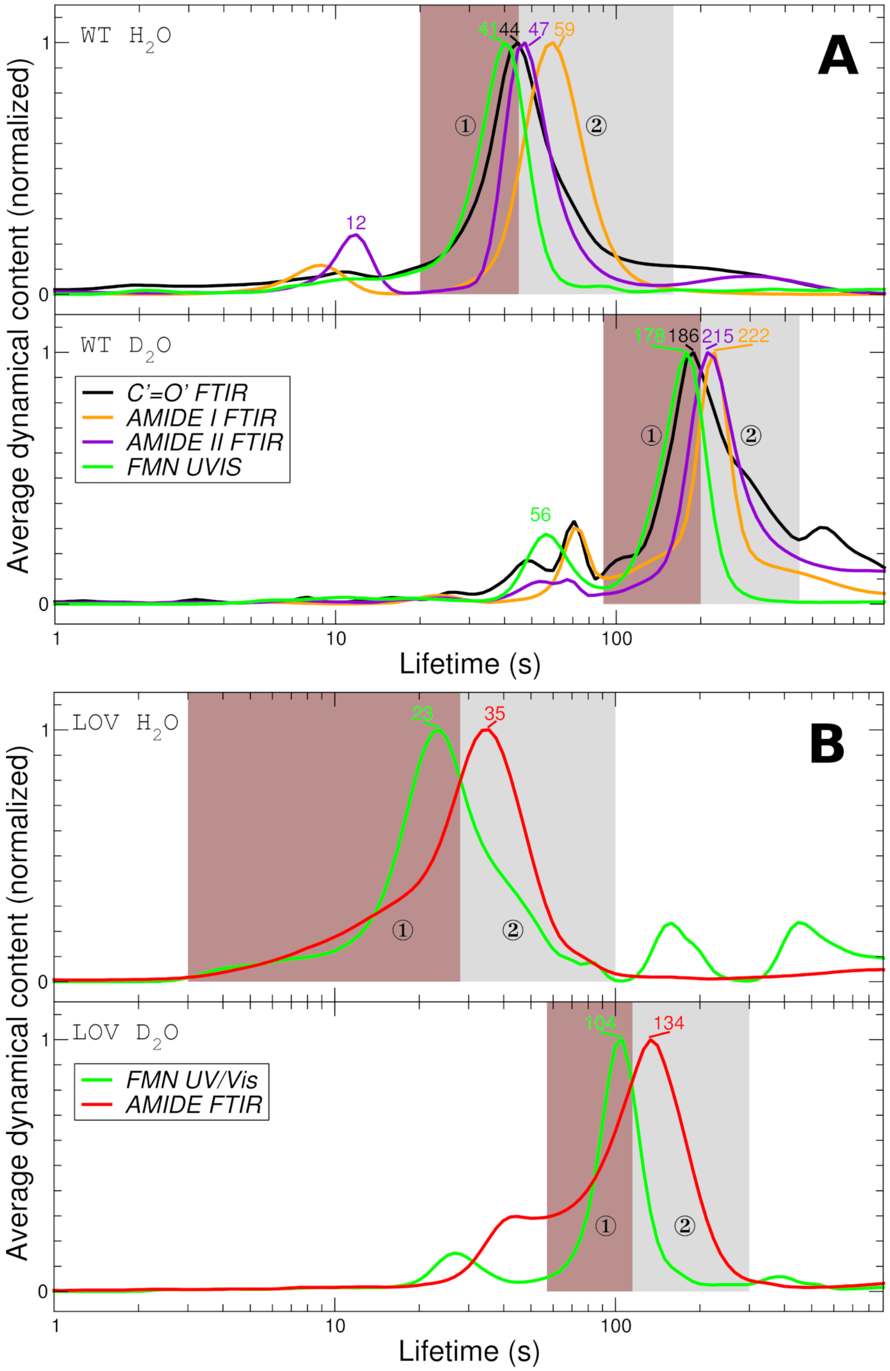

**Average dynamical content of the main EL222 variants. (A)** Wt EL222 in $H_2O$ (top) vs $D_2O$ (bottom). The IR region has been divided into three: FMN carbonyls (1750-1680 $cm^{-1}$), amide I (1680-1600 $cm^{-1}$), and amide II (1600-1500 $cm^{-1}$). **(B)** Isolated LOV domain of EL222 in $H_2O$ (top) *vs.* $D_2O$ (bottom). Two probes have been used: FMN (320-550 nm) and amide bands (1750-1500 $cm^{-1}$). All datasets arise from lifetime distribution analysis using the maximum entropy method. The two shaded areas highlight the main kinetic event sensed by the FMN (number one in the figure) and amide (number two in the figure) probes.

### FIGURE S11.

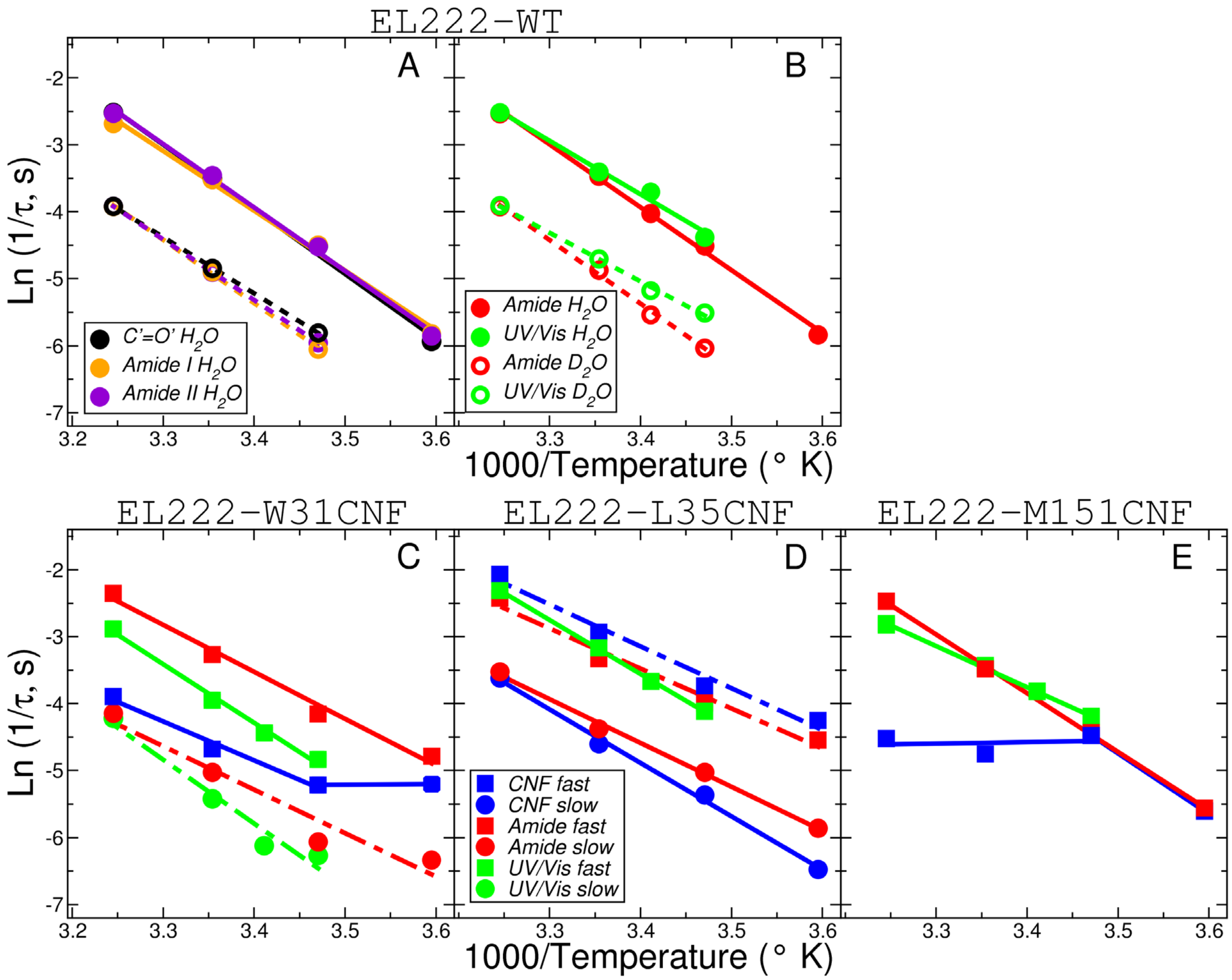

**Effect of temperature on the lit-to-dark kinetics of wt EL222 and single-CNF mutants. (A)** Arrhenius plots of wt EL222 at three different IR probe frequencies: 1750-1680 $cm^{-1}$ (FMN carbonyls, black), 1680-1600 $cm^{-1}$ (protein amide I band, orange), and 1600-1500 $cm^{-1}$ (protein amide II band among others, violet). **(B)** Arrhenius plot of the main kinetic component at two probe frequencies: 1750-1500 $cm^{-1}$ (amide bands, red) and 320-550 nm (FMN UV/Vis, green). In (A) and (B), samples measured in $H_2O$ and $D_2O$ are shown as open circles and closed circles, respectively. Lines are linear fits to the data (solid lines for data in $H_2O$ and dotted lines for data in $D_2O$). **(C)** Arrhenius plots of EL222 W31CNF at three probe frequencies. **(D)** Arrhenius plots of EL222 L35CNF at three probe frequencies. **(E)** Arrhenius plots of EL222 M151CNF at three probe frequencies. In (C), (D) and (E) the monitored frequencies/wavelengths are 2200-2270 $cm^{-1}$ (nitrile band of CNF residue, blue) 1750-1500 $cm^{-1}$ (amide bands, red) and 320-550 nm (FMN UV/Vis, green). The fastest kinetic components are shown as squares while the slowest kinetic events are shown as circles (if only a major kinetic component is present, squares are used). Lines are linear fits to the data (solid lines for the most abundant kinetic event and dotted lines for the least abundant kinetic event).

#### FIGURE S12.

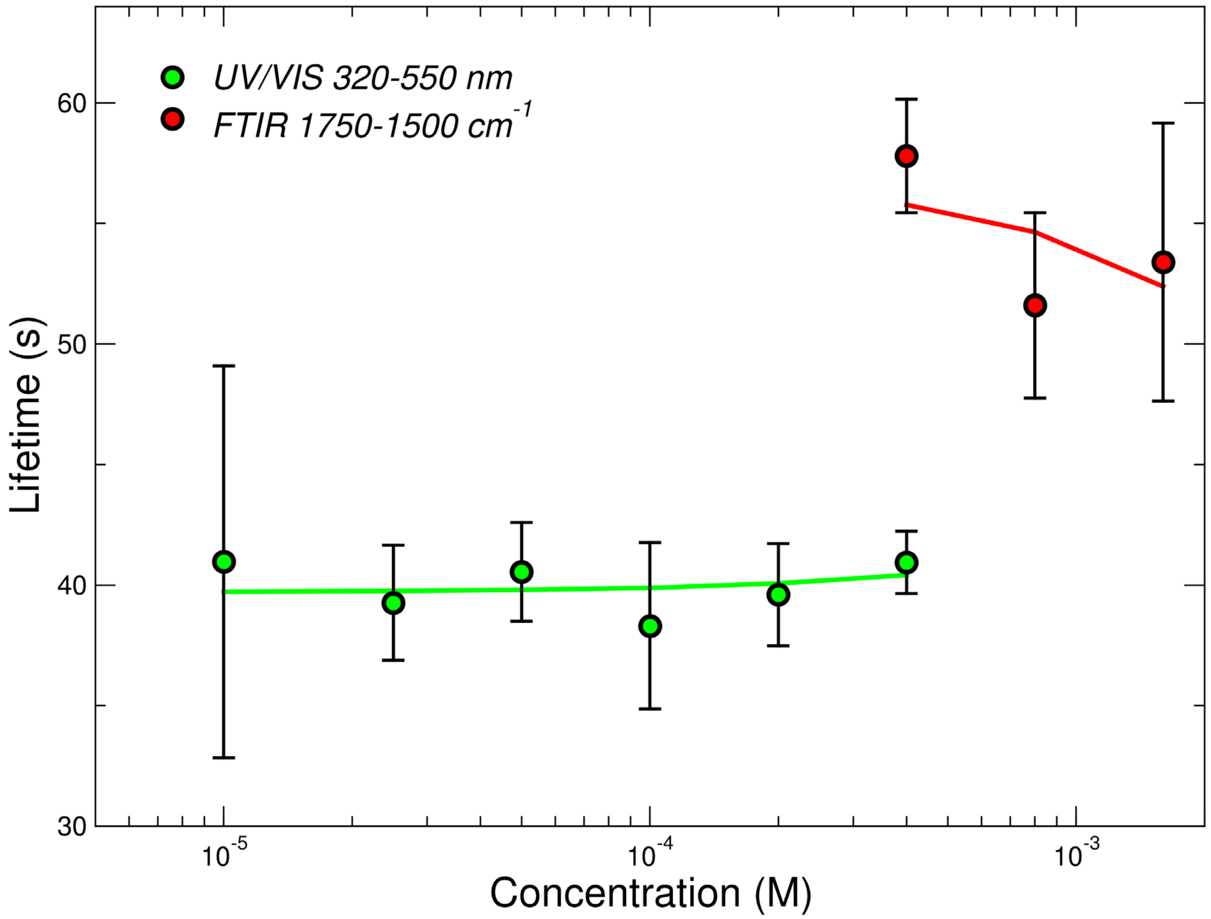

**Effect of protein concentration on the lit-to-dark kinetics of wt EL222 in $H_2O$.** Recovery lifetimes are plotted as a function of concentration for two probe frequencies: 1750-1500 $cm^{-1}$ (amide bands, red circles) and 320-550 nm (FMN, green circles). Solid lines are linear fits to the data. Error bars represent the standard deviation of 3 independent measurements.

### FIGURE S13.

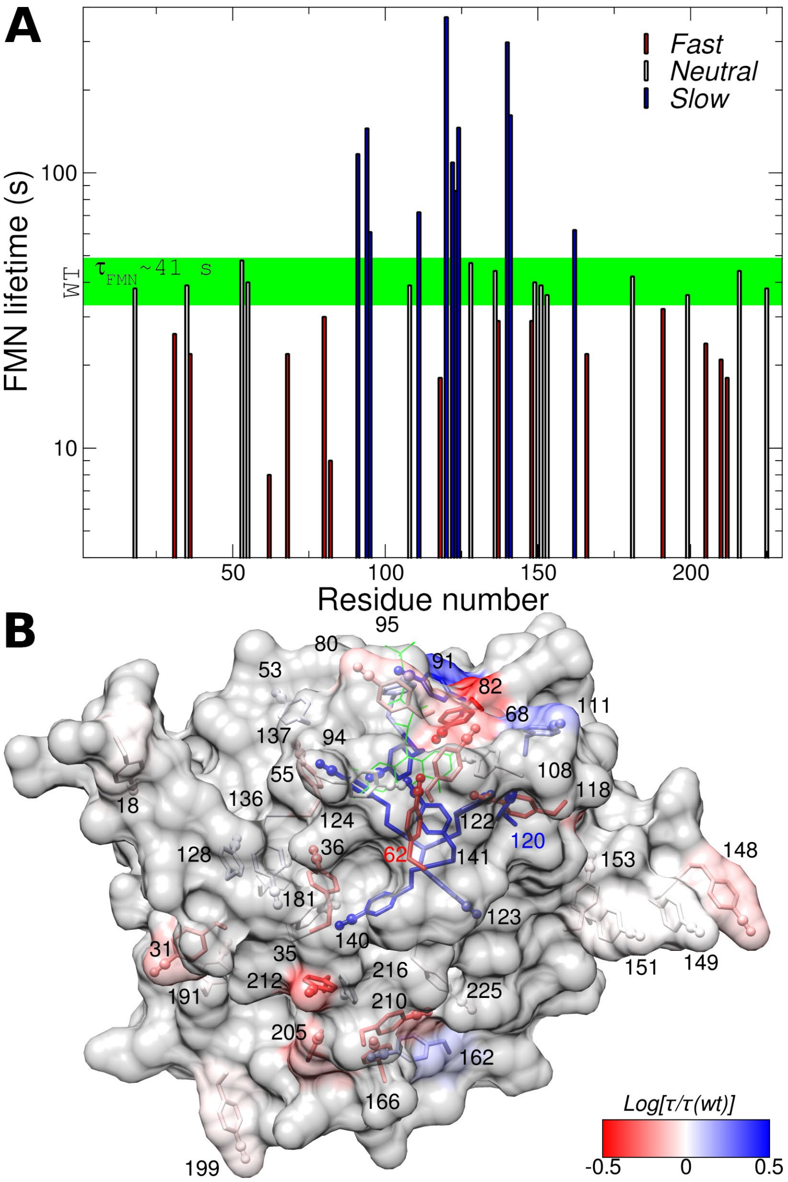

**Mapping FMN photorecovery lifetimes in 1D and 3D. (A)** FMN lifetimes measured by time-resolved UV/Vis spectroscopy as a function of residue number. Neutral mutants are defined as those having similar lifetimes as wild-type EL222 ($\tau_{FMN} = 41 \pm 7$ s). **(B)** Structure of dark-adapted EL222 containing 39 CNF residues color-coded according to the FMN recovery lifetime ($log[\tau_{FMN}/\tau_{FMN}(wt)]$) modeled onto PDB id 3P7N. In both panels, the residues with the fastest and slowest lifetimes are highlighted in red and blue, respectively.

### FIGURE S14.

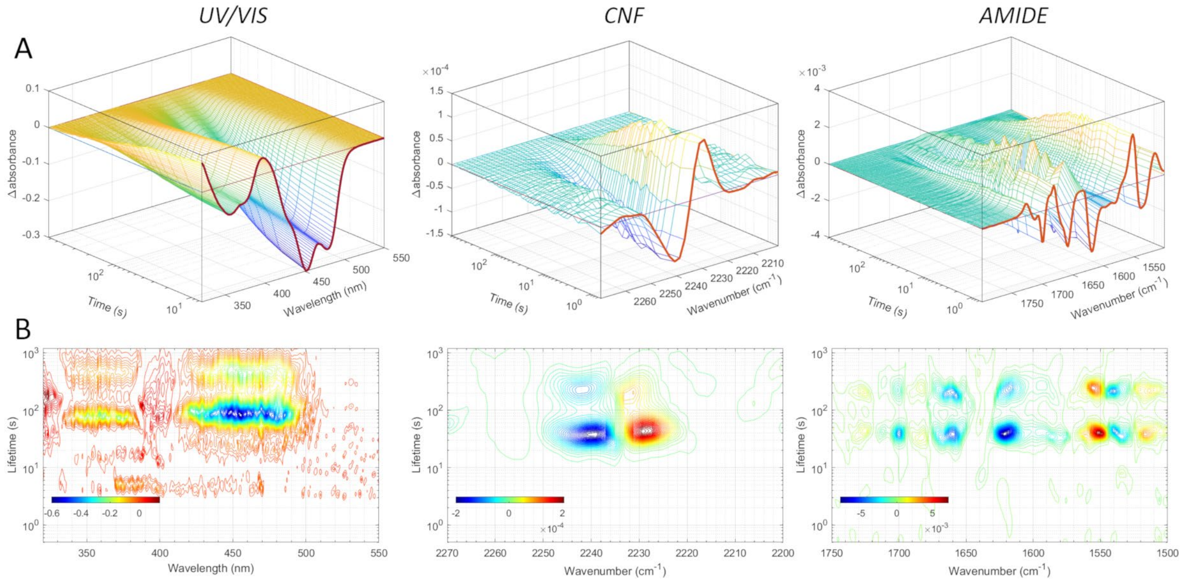

**Time-resolved UV/VIS/IR spectroscopy of EL222 W31CNF. (A)** 3D pre-processed datasets of differential absorbance as a function of time and frequency. **(B)** 2D lifetime distribution plots using the maximum entropy method. The left, middle and right panels show the FMN (320-550 nm), CNF (2270-2200 $cm^{-1}$), and amide (1750-1500 $cm^{-1}$) probes, respectively.

### FIGURE S15.

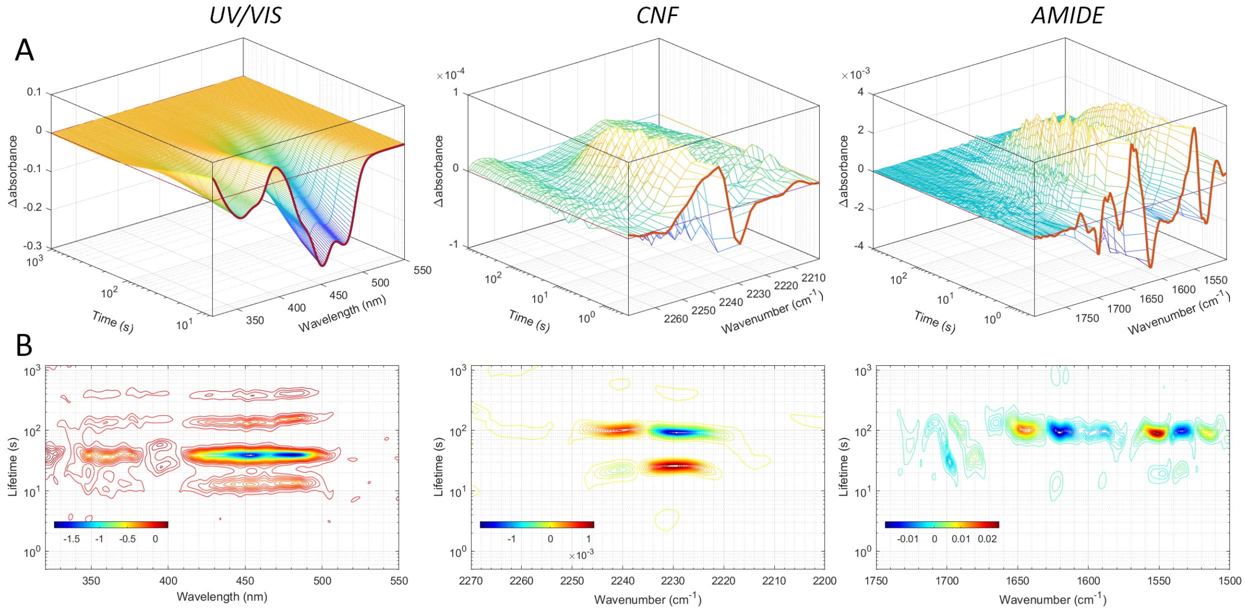

**Time-resolved UV/VIS/IR spectroscopy of EL222 L35CNF. (A)** 3D pre-processed datasets of differential absorbance as a function of time and frequency. **(B)** 2D lifetime distribution plots using the maximum entropy method. The left, middle and right panels show the FMN (320-550 nm), CNF (2270-2200 $cm^{-1}$), and amide (1750-1500 $cm^{-1}$) probes, respectively.

#### FIGURE S16.

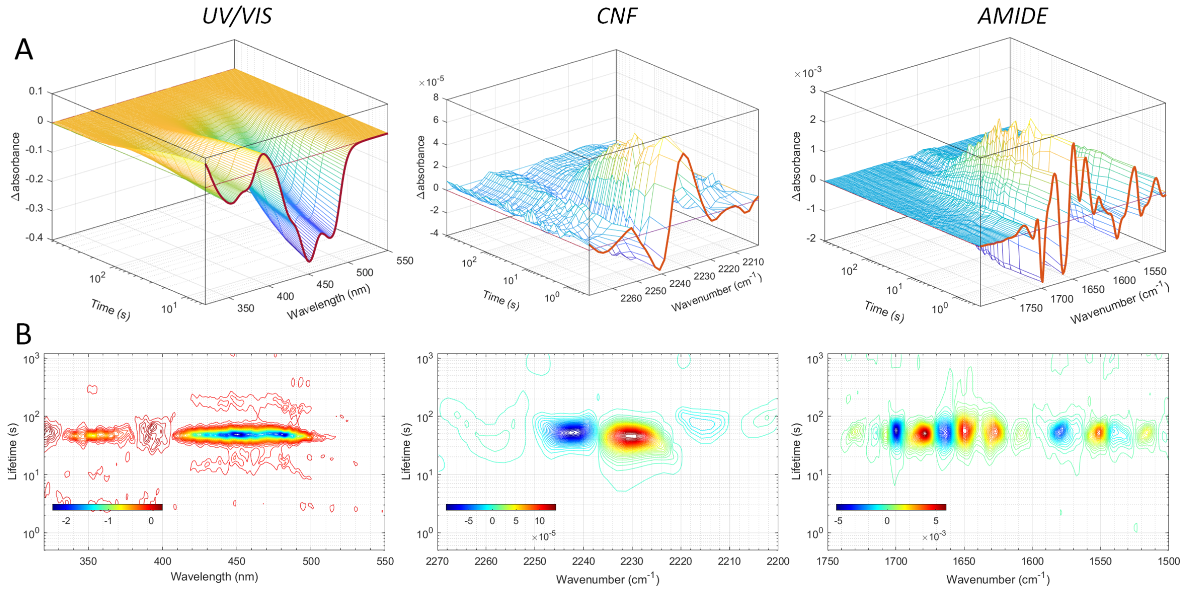

**Time-resolved UV/VIS/IR spectroscopy of EL222 N53CNF. (A)** 3D pre-processed datasets of differential absorbance as a function of time and frequency. **(B)** 2D lifetime distribution plots using the maximum entropy method. The left, middle and right panels show the FMN (320-550 nm), CNF (2270-2200 cm$^{-1}$), and amide (1750-1500 cm$^{-1}$) probes, respectively.

#### FIGURE S17.

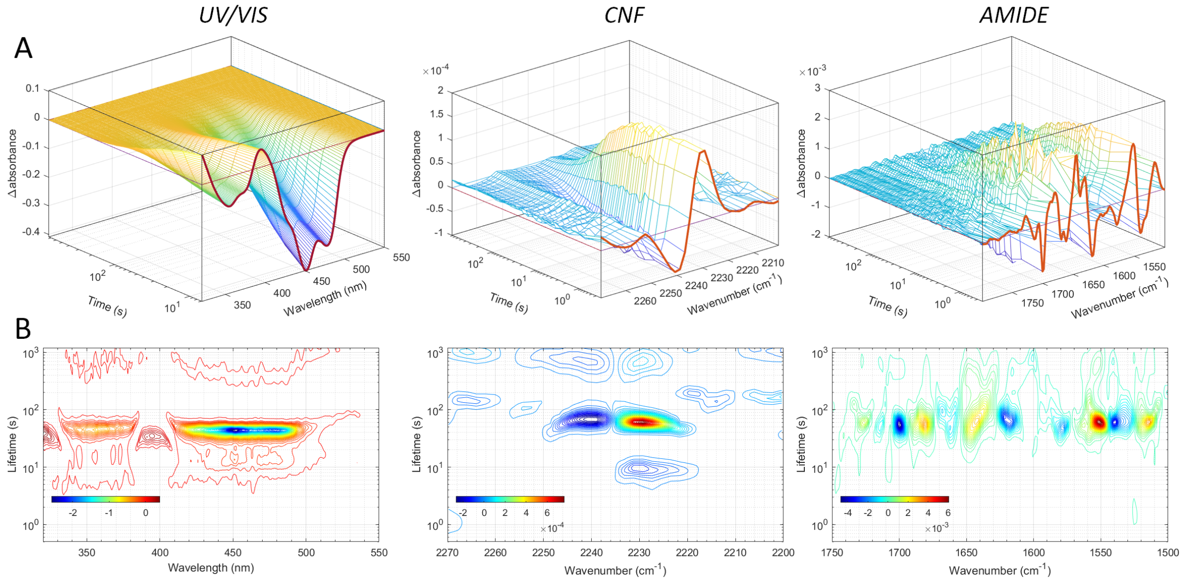

**Time-resolved UV/VIS/IR spectroscopy of EL222 Y136CNF. (A)** 3D pre-processed datasets of differential absorbance as a function of time and frequency. **(B)** 2D lifetime distribution plots using the maximum entropy method. The left, middle and right panels show the FMN (320-550 nm), CNF (2270-2200 cm$^{-1}$), and amide (1750-1500 cm$^{-1}$) probes, respectively.

### FIGURE S18.

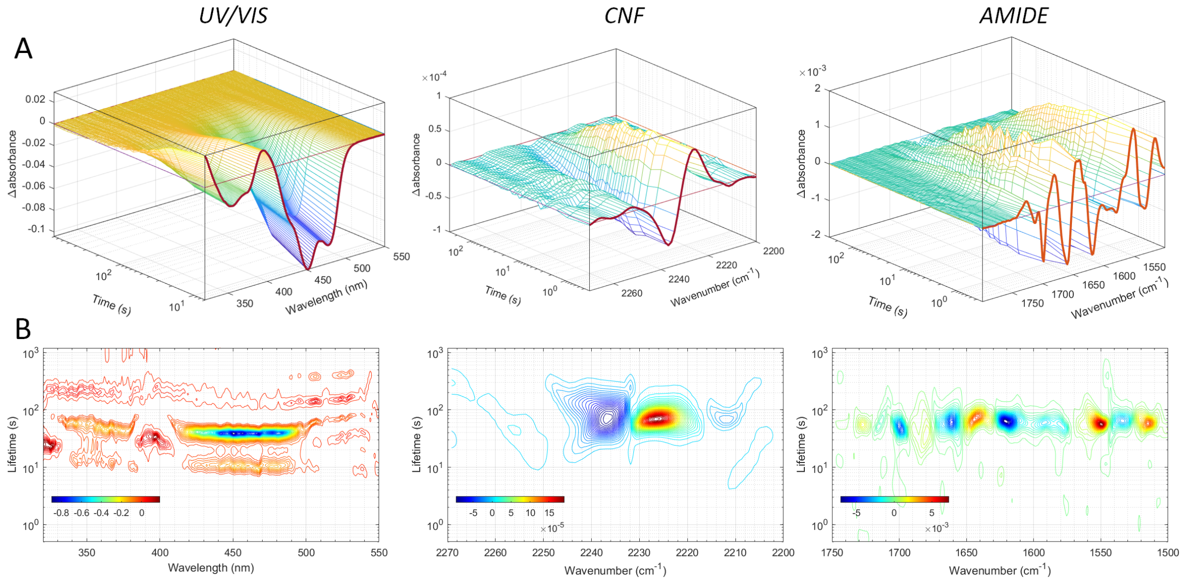

**Time-resolved UV/VIS/IR spectroscopy of EL222 M151CNF. (A)** 3D pre-processed datasets of differential absorbance as a function of time and frequency. **(B)** 2D lifetime distribution plots using the maximum entropy method. The left, middle and right panels show the FMN (320-550 nm), CNF (2270-2200 cm$^{-1}$), and amide (1750-1500 cm$^{-1}$) probes, respectively.

### FIGURE S19.

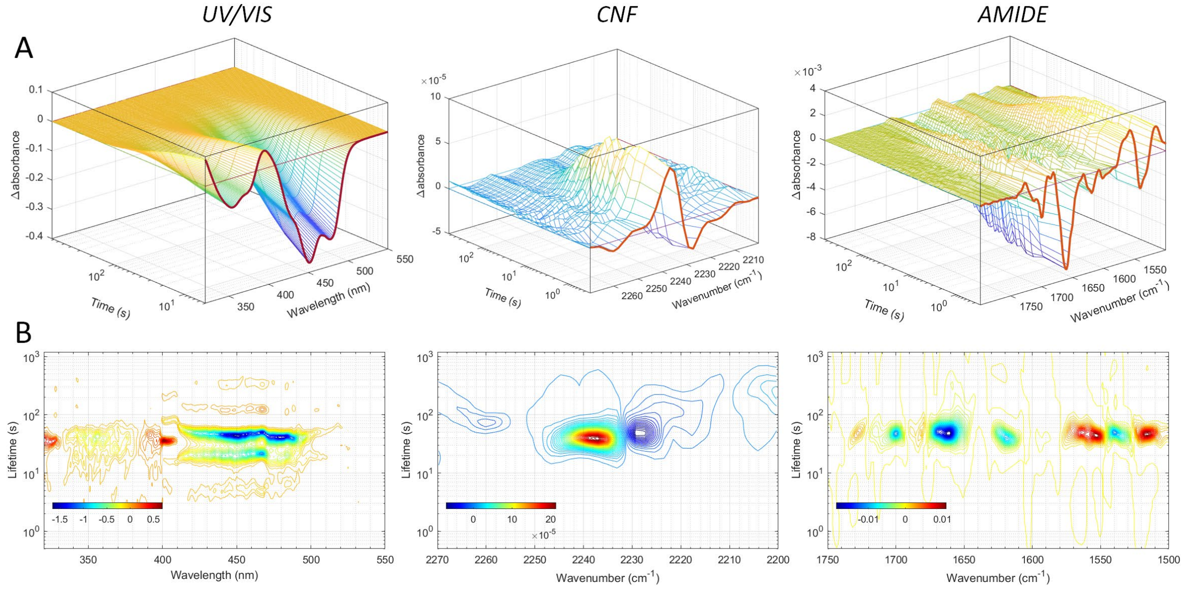

**Time-resolved UV/VIS/IR spectroscopy of EL222 L216CNF. (A)** 3D pre-processed datasets of differential absorbance as a function of time and frequency. **(B)** 2D lifetime distribution plots using the maximum entropy method. The left, middle and right panels show the FMN (320-550 nm), CNF (2270-2200 cm$^{-1}$), and amide (1750-1500 cm$^{-1}$) probes, respectively.

### FIGURE S20.

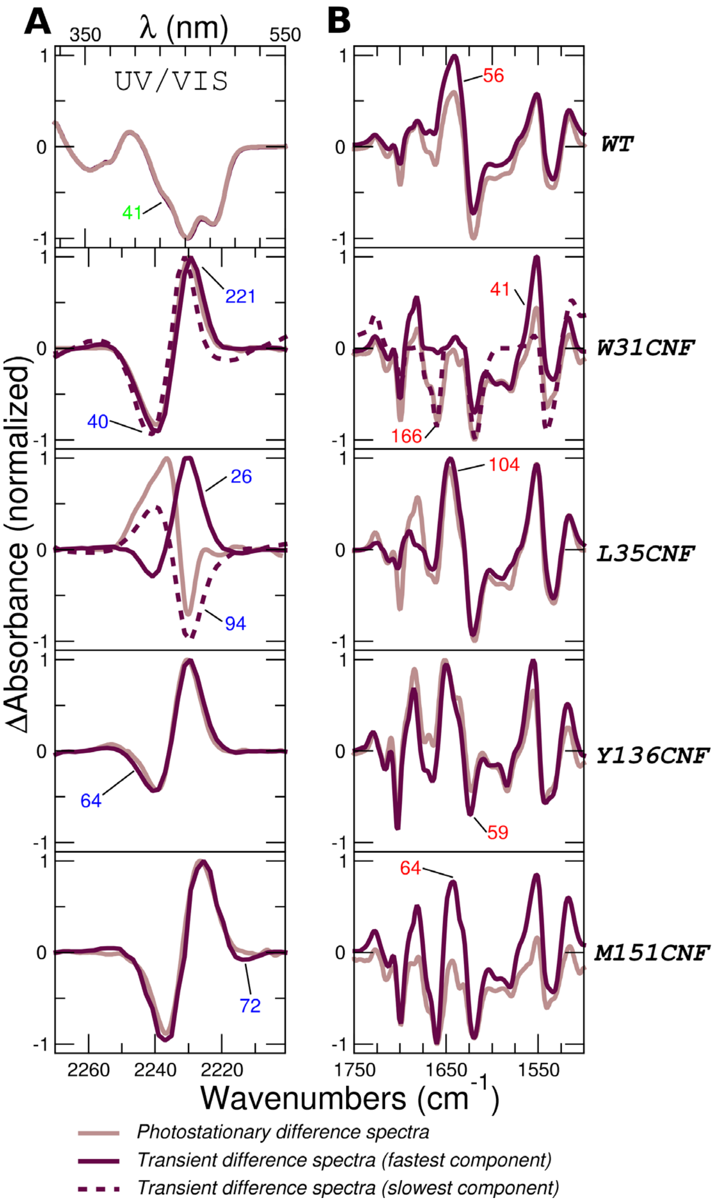

**Lifetime distribution analysis of time-resolved spectroscopy data of wt EL222 and single-CNF variants. (A)** Comparison between the transient difference spectra (decay-associated difference spectra or DADS) and photostationary difference spectra (steady-state difference spectra or SSDS) in the CNF absorption region (normalized from the spectra shown in Fig. 2A). In the case of wt EL222, which lacks CNF bands, transient and steady-stated UV/Vis spectra are shown instead (top row). **(B)** Same as panel A but with transient and photostationary spectra showing the amide region. In most cases, a single component (i.e. mono-exponential function) suffices to describe the data and thus transient and stationary spectra resemble each other. Two components are needed to fit the kinetics of W31CNF and L35CNF mutants. The numbers indicate the lifetimes (in seconds) of the corresponding dynamical events color-coded according to the probe (green for FMN, red for amide, and blue for CNF).

#### FIGURE S21.

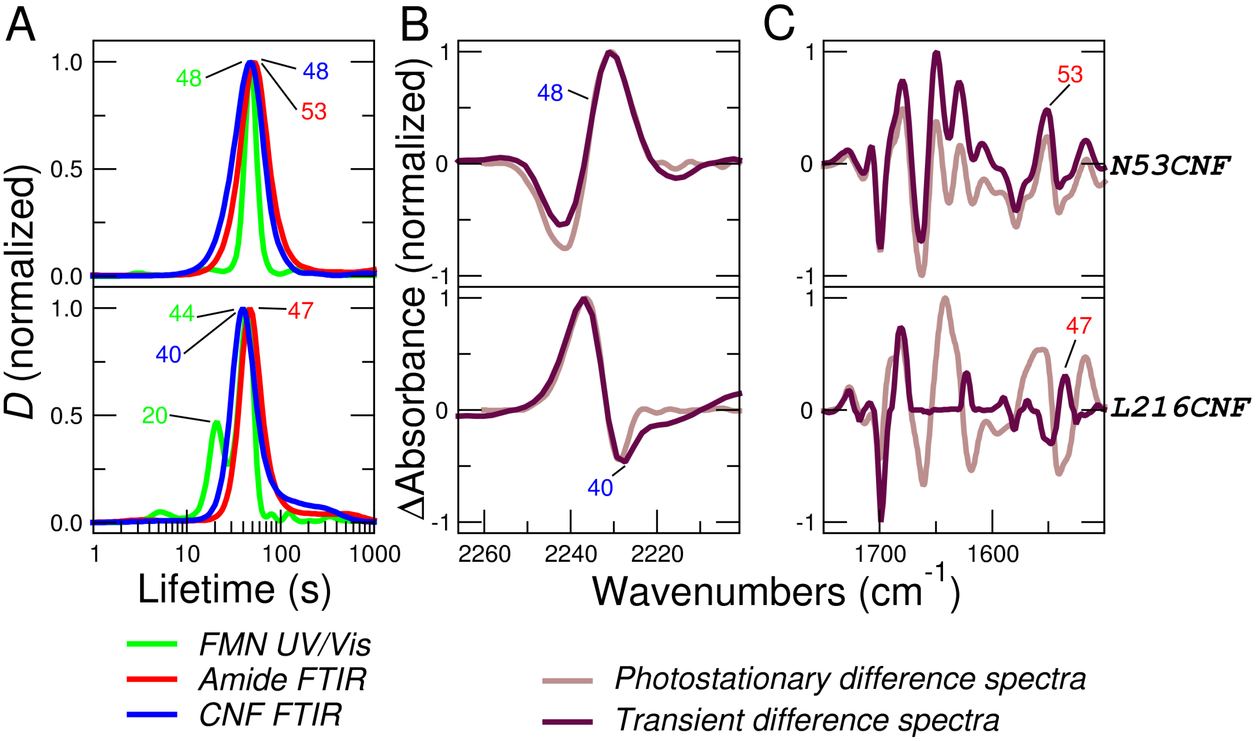

**Lifetime distribution analysis of time-resolved spectroscopy data of other single-CNF variants. (A)** Average dynamical content (*D*) as a function of lifetime for two single-CNF mutants (N53CNF and L216CNF) and three probes: FMN (green lines), amide (red lines) and the cyano moiety of the CNF residue (blue lines). The numbers report the lifetimes (in seconds) of the main dynamical events (peak maxima) again color-coded as green, red and blue for FMN, amide, and CNF probes, respectively. **(B)** Comparison between the integrated transient difference spectra (equivalent to a decay-associated difference spectra or DADS) and steady-state difference spectra in the CNF absorption region (SSDS, normalized from the spectra shown in Fig. 2A). **(C)** Same as panel B but with transient and photostationary spectra showing the amide region.

### FIGURE S22.

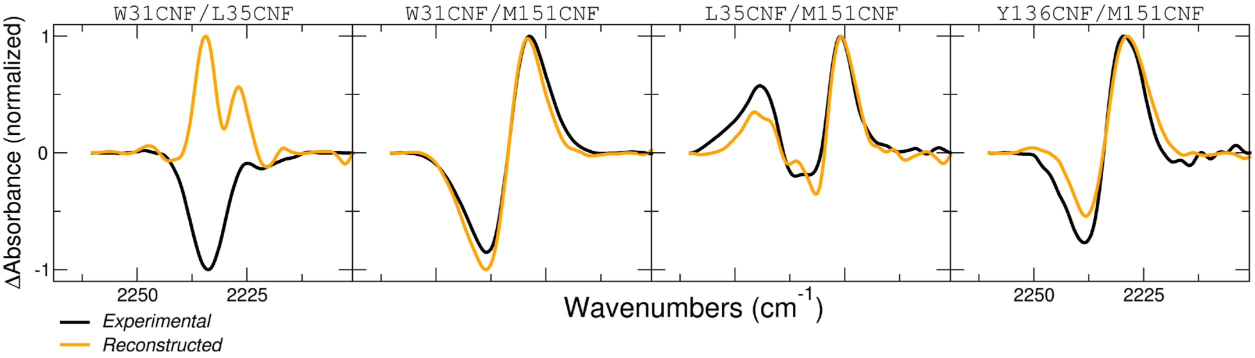

**Predictions of steady-state difference spectra (SSDS) of double-CNF variants.** Experimental SSDS (black lines) of the double-CNF mutants, indicated on top of the panels, were reconstructed by a linear combination of the two corresponding single-CNF SSDS. The only adjustable parameter was the fraction of each spectra. Best fist spectra (lowest $\chi^2$) are colored in orange.

### FIGURE S23.

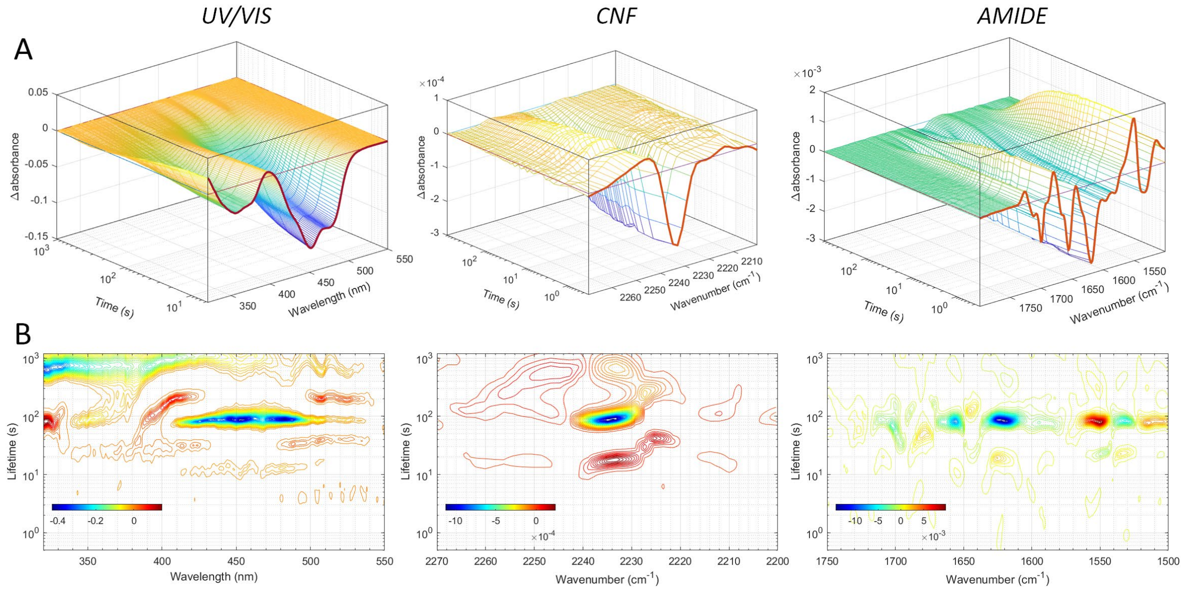

**Time-resolved UV/VIS/IR spectroscopy of EL222 W31CNF/L35CNF. (A)** 3D pre-processed datasets of differential absorbance as a function of time and frequency. **(B)** 2D lifetime distribution plots using the maximum entropy method. The left, middle and right panels show the FMN (320-550 nm), CNF (2270-2200 cm$^{-1}$), and amide (1750-1500 cm$^{-1}$) probes, respectively.

### FIGURE S24.

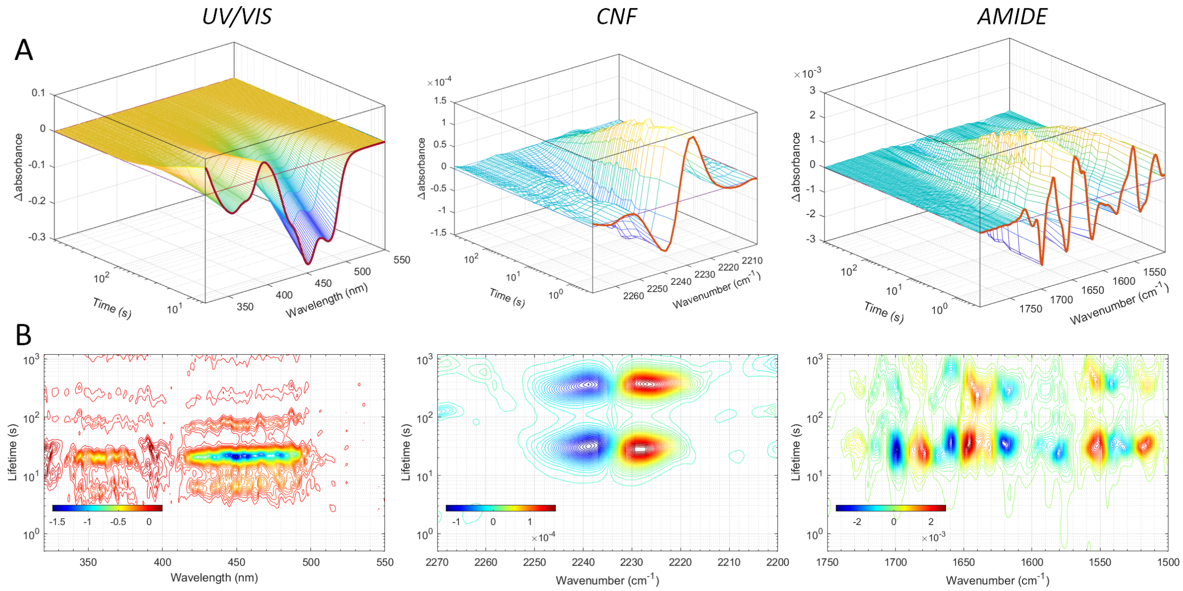

**Time-resolved UV/VIS/IR spectroscopy of EL222 W31CNF/M151CNF. (A)** 3D pre-processed datasets of differential absorbance as a function of time and frequency. **(B)** 2D lifetime distribution plots using the maximum entropy method. The left, middle and right panels show the FMN (320-550 nm), CNF (2270-2200 cm$^{-1}$), and amide (1750-1500 cm$^{-1}$) probes, respectively.

### FIGURE S25.

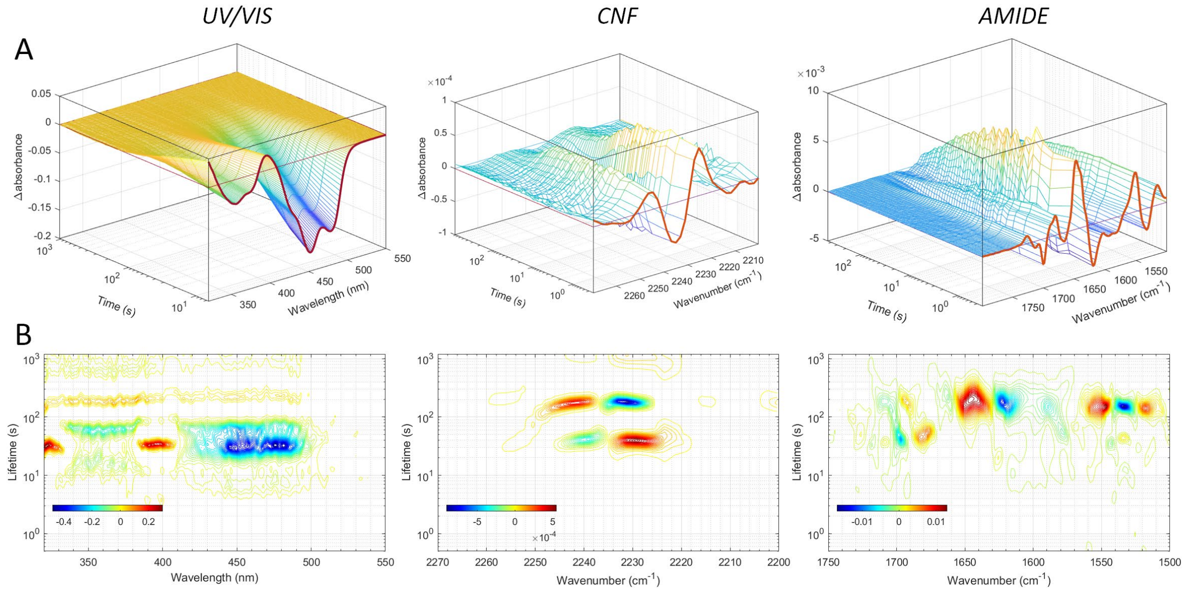

**Time-resolved UV/VIS/IR spectroscopy of EL222 L35CNF/M151CNF. (A)** 3D pre-processed datasets of differential absorbance as a function of time and frequency. **(B)** 2D lifetime distribution plots using the maximum entropy method. The left, middle and right panels show the FMN (320-550 nm), CNF (2270-2200 $cm^{-1}$), and amide (1750-1500 $cm^{-1}$) probes, respectively.

### FIGURE S26.

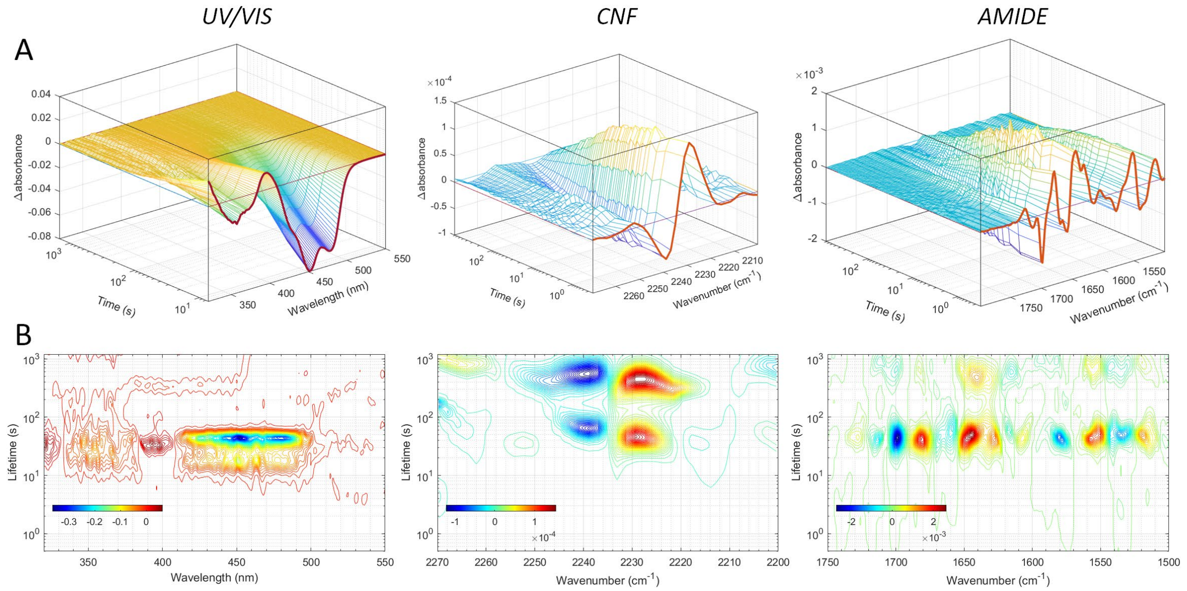

**Time-resolved UV/VIS/IR spectroscopy of EL222 Y136CNF/M151CNF. (A)** 3D pre-processed datasets of differential absorbance as a function of time and frequency. **(B)** 2D lifetime distribution plots using the maximum entropy method. The left, middle and right panels show the FMN (320-550 nm), CNF (2270-2200 $cm^{-1}$), and amide (1750-1500 $cm^{-1}$) probes, respectively.

### FIGURE S27.

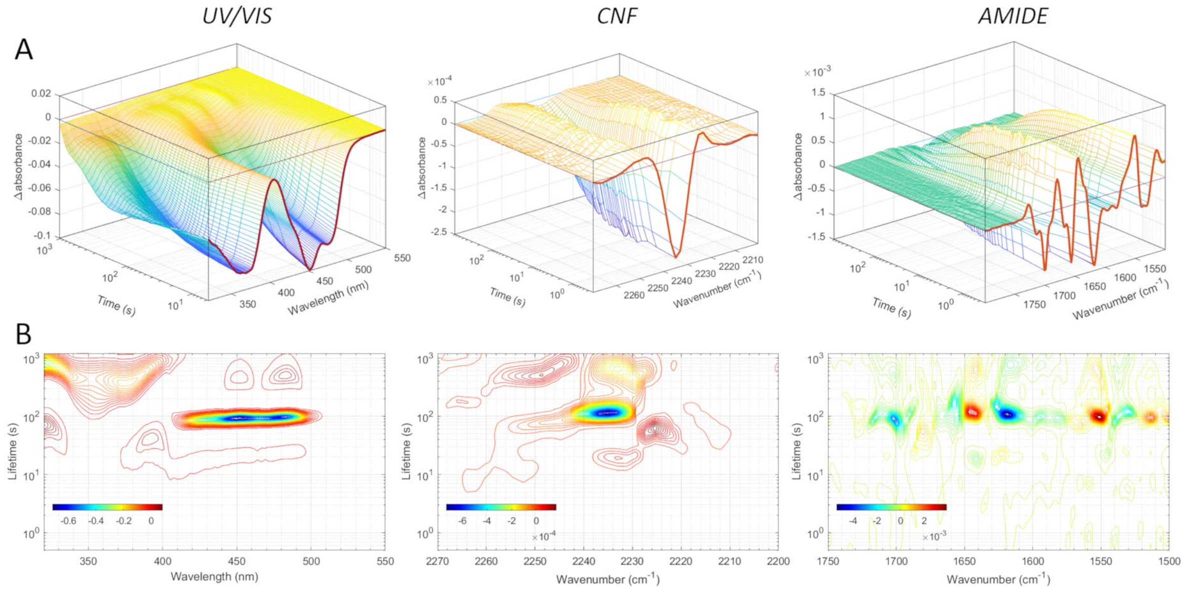

**Time-resolved UV/VIS/IR spectroscopy of EL222 W31CNF/L35CNF/M151CNF**. **(A)** 3D pre-processed datasets of differential absorbance as a function of time and frequency. **(B)** 2D lifetime distribution plots using the maximum entropy method. The left, middle and right panels show the FMN (320-550 nm), CNF (2270-2200 $cm^{-1}$), and amide (1750-1500 $cm^{-1}$) probes, respectively.

### FIGURE S28.

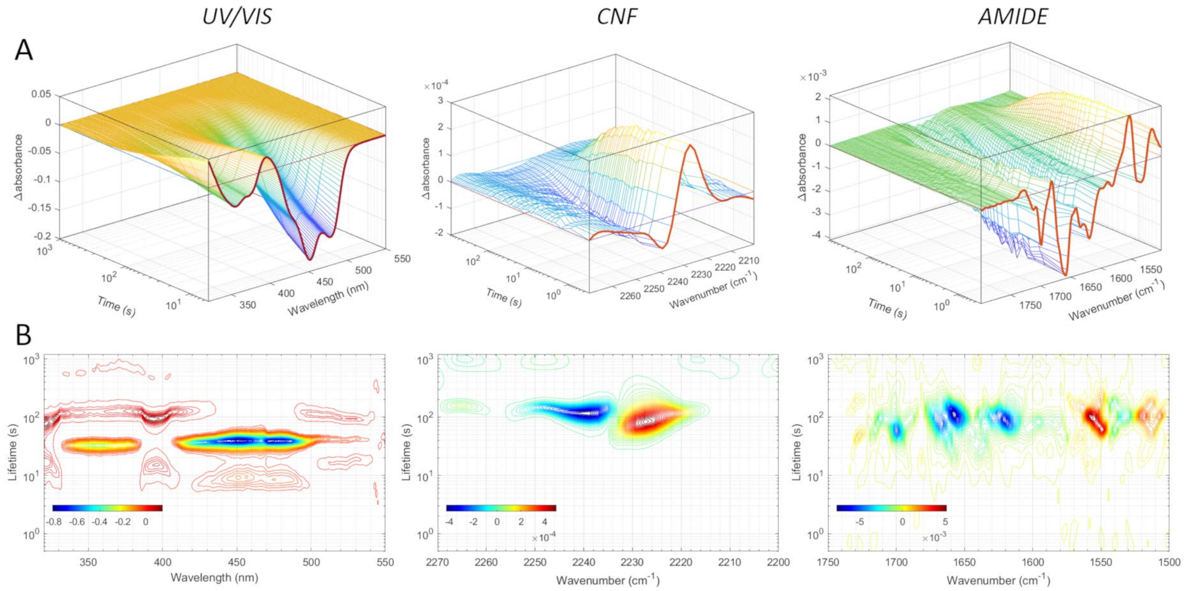

**Time-resolved UV/VIS/IR spectroscopy of EL222 L35CNF/Y136CNF/M151CNF. (A)** 3D pre-processed datasets of differential absorbance as a function of time and frequency. **(B)** 2D lifetime distribution plots using the maximum entropy method. The left, middle and right panels show the FMN (320-550 nm), CNF (2270-2200 $cm^{-1}$), and amide (1750-1500 $cm^{-1}$) probes, respectively.

### FIGURE S29.

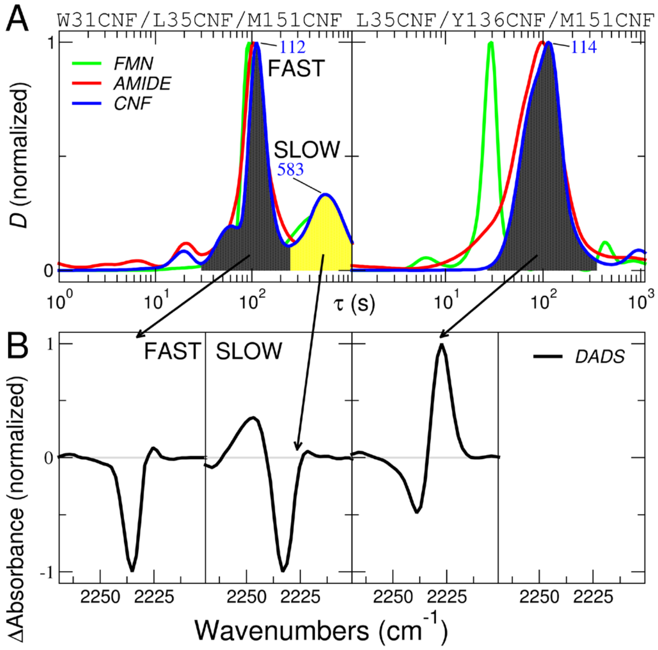

**Disentangling residue-by-residue relaxation rates with triple-CNF EL222 mutants. (A)** Average dynamical content ($D$) as a function of lifetime ($\tau$), derived from lifetime distribution analysis (Figure S13-S24 B) for two triple-CNF mutants (W31CNF/L35CNF/M151CNF and L35CNF/Y136CNF/M151CNF). Curves were calculated for the relaxation of the FMN (320-450 nm, green lines), amide bands (including also contributions from FMN, 1750-1500 cm$^{-1}$, red lines), and the cyano moiety of the CNF residue (2270-2200 cm$^{-1}$, blue lines). The peak maxima, corresponding to the most probable lifetime (in seconds) are indicated in the case of CNF relaxation. **(B)** The transient spectra associated with the kinetic events seen in the lit-to-dark transition of the cyano moiety were derived by integration over the shaded areas (black for the fastest event and cyan for the slowest event).

### FIGURE S30.

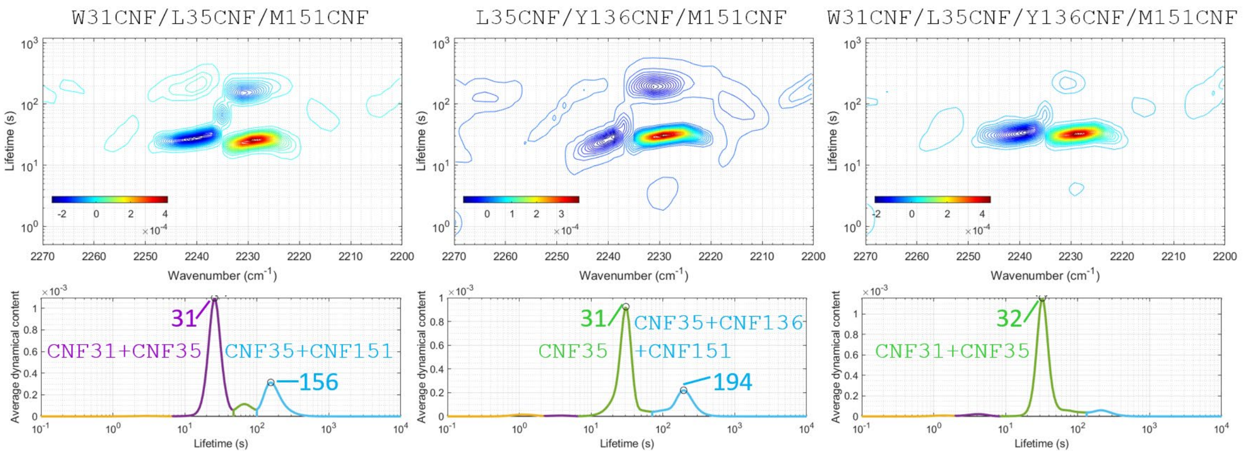

**Simulations of lifetime density maps and average dynamical contents (*D*).** Synthetic datasets of triple- and quadruple-CNF EL222 variants were generated by averaging the experimental datasets of the corresponding single-CNF variants. Subsequently, they were analyzed using the maximum entropy method following the same procedures as for the experimental data. The lifetimes (in seconds) of the main peaks are indicated together with the contributing residues.

#### FIGURE S31.

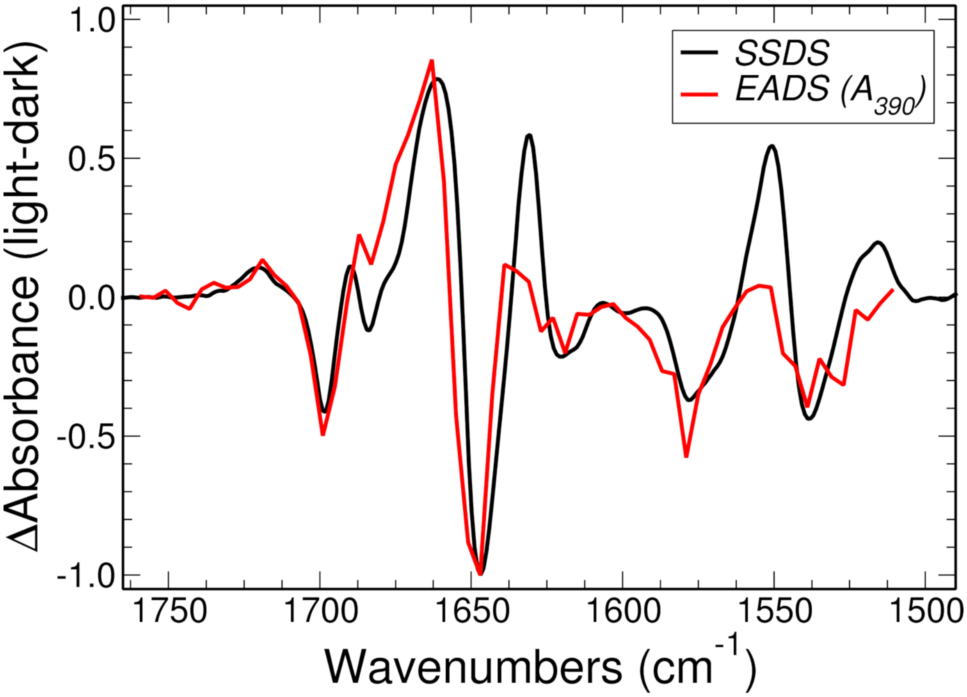

**Comparison of transient and stationary IR spectra of wt EL222 in $D_2O$.** The evolution-associated difference spectra (EADS) of the adduct state (corresponding to the $A_{390}$ species), which is formed 5 µs after photoexcitation is shown in red, while the steady-state difference spectrum (SSDS) is shown in black.

#### SUPPLEMENTARY REFERENCES.

81. J. S. Beckwith, C. A. Rumble, E. Vauthey, Data analysis in transient electronic spectroscopy – an experimentalist's view. International Reviews in Physical Chemistry 39, 135-216 (2020).
82. C. Slavov, H. Hartmann, J. Wachtveitl, Implementation and Evaluation of Data Analysis Strategies for Time-Resolved Optical Spectroscopy. Analytical Chemistry 87, 2328-2336 (2015).
83. V. A. Lórenz-Fonfría, H. Kandori, Transformation of Time-Resolved Spectra to Lifetime-Resolved Spectra by Maximum Entropy Inversion of the Laplace Transform. Applied Spectroscopy 60, 407-417 (2016).
84. V. A. Lórenz-Fonfría, H. Kandori, Practical Aspects of the Maximum Entropy Inversion of the Laplace Transform for the Quantitative Analysis of Multi-Exponential Data. Applied Spectroscopy 61, 74-84 (2016).
85. P. C. Hansen, D. P. O'Leary, The Use of the L-Curve in the Regularization of Discrete Ill-Posed Problems. SIAM Journal on Scientific Computing 14, 1487-1503 (1993).
86. B. Webb, A. Sali, Comparative Protein Structure Modeling Using MODELLER. Current Protocols in Bioinformatics 54,  (2016).
87. D. Gfeller, O. Michielin, V. Zoete, SwissSidechain: a molecular and structural database of non-natural sidechains. Nucleic Acids Res 41, D327-332 (2013).
88. W. Humphrey, A. Dalke, K. Schulten, VMD: Visual molecular dynamics. Journal of Molecular Graphics 14, 33-38 (1996).
89. T. R. Alderson, L. E. Kay, NMR spectroscopy captures the essential role of dynamics in regulating biomolecular function. Cell 184, 577-595 (2021).
90. M. Maj, J. P. Lomont, K. L. Rich, A. M. Alperstein, M. T. Zanni, Site-specific detection of protein secondary structure using 2D IR dihedral indexing: a proposed assembly mechanism of oligomeric hIAPP. Chemical Science 9, 463-474 (2018).
91. M. Amiram, A. D. Haimovich, C. Fan, Y.-S. Wang, H.-R. Aerni, I. Ntai, D. W. Moonan, N. J. Ma, A. J. Rovner, S. H. Hong, N. L. Kelleher, A. L. Goodman, M. C. Jewett, D. Söll, J. Rinehart, F. J. Isaacs, Evolution of translation machinery in recoded bacteria enables multi-site incorporation of nonstandard amino acids. Nature Biotechnology 33, 1272-1279 (2015).
92. D. L. Dunkelmann, S. B. Oehm, A. T. Beattie, J. W. Chin, A 68-codon genetic code to incorporate four distinct non-canonical amino acids enabled by automated orthogonal mRNA design. Nature Chemistry 13, 1110-1117 (2021).
93. P. Fischer, S. Mukherjee, E. Peter, M. Broser, F. Bartl, P. Hegemann, The inner mechanics of rhodopsin guanylyl cyclase during cGMP-formation revealed by real-time FTIR spectroscopy. eLife 10,  (2021).
94. J. E. Hart, K. H. Gardner, Lighting the way: Recent insights into the structure and regulation of phototropin blue light receptors. Journal of Biological Chemistry 296,  (2021).
95. K. Magerl, B. Dick, Dimerization of LOV domains of Rhodobacter sphaeroides (RsLOV) studied with FRET and stopped-flow experiments. Photochemical & Photobiological Sciences 19, 159-170 (2020).
96. D. Nohr, R. Rodriguez, S. Weber, E. Schleicher, How can EPR spectroscopy help to unravel molecular mechanisms of flavin-dependent photoreceptors? Frontiers in Molecular Biosciences 2,  (2015).
97. R. A. Laskowski, M. B. Swindells, LigPlot+: Multiple Ligand–Protein Interaction Diagrams for Drug Discovery. Journal of Chemical Information and Modeling 51, 2778-2786 (2011).
98. M. Granell, X. León, G. Leblanc, E. Padrós, V. A. Lórenz-Fonfría, Structural insights into the activation mechanism of melibiose permease by sodium binding. Proceedings of the National Academy of Sciences 107, 22078-22083 (2010).
